# Identification of the orphan GPR25 as a receptor of the chemokine CXCL17

**DOI:** 10.1101/2024.09.21.614233

**Authors:** Wen-Feng Hu, Jie Yu, Juan-Juan Wang, Ru-Jiao Sun, Yong-Shan Zheng, Teng Zhang, Ya-Li Liu, Zeng-Guang Xu, Zhan-Yun Guo

**Author notes:** **Correspondence**, Z.-Y. Guo, Research Center for Translational Medicine at East Hospital, School of Life Sciences and Technology, Tongji University, 1239 Siping Road, Shanghai 200092, China. Wen-Feng Hu and Jie Yu contributed to equally to this work.

## Abstract

C-X-C motif chemokine ligand 17 (CXCL17) is a small secretory protein primarily expressed in mucosal tissues and likely functions as a chemoattractant, but its receptor is still controversial. In the present study, we identified the rarely studied orphan G protein-coupled receptor 25 (GPR25) as a receptor of CXCL17 via prediction by the newly developed AlphaFold 3 algorithm and validation by the NanoBiT-based β-arrestin recruitment assay. In the β-arrestin recruitment assay, recombinant human CXCL17 could activate human GPR25 in transfected human embryonic kidney (HEK) 293T cells with an EC_50_ value around 100 nM, but it had no activation effect on 17 other tested G protein-coupled receptors (GPCRs). Deletion of three conserved C-terminal residues of human CXCL17 almost abolished its activity, and alanine replacement of W95 or R178 of human GPR25, two conserved residues in the predicted orthosteric ligand binding pocket, almost abolished its response towards CXCL17. These results are consistent with the AlphaFold 3-predicted binding model in which the highly conserved C-terminal fragment of CXCL17 inserts into the orthosteric ligand binding pocket of the receptor GPR25. According to the expression pattern of CXCL17 and GPR25 shown at the Human Protein Atlas, CXCL17 might be an endogenous agonist of GPR25 in human and other mammals, but this hypothesis needs to be tested in future studies via more assays. The present deorphanization paves the way for further functional characterization of the orphan receptor GPR25 and the orphan ligand CXCL17.

## Introduction

C-X-C motif chemokine ligand 17 (CXCL17) is a small secretory protein first identified in 2006 [1□6]. It was formerly named dendritic cell and monocyte chemokine-like protein (DMC) or VEGF correlated chemokine 1 (VCC-1) in early studies [1,2]. CXCL17 is primarily expressed in some mucosal tissues, such as lung, stomach, esophagus, and salivary gland, according to the data shown at Human Protein Atlas (https://www.proteinatlas.org/ENSG00000189377-CXCL17), and likely functions as a chemoattractant for monocytes, macrophages, and dendritic cells [1,3□13]. Knockout of *Cxcl17* gene in mouse resulted in a significantly reduced number of macrophages in lung, and abnormal functions of T cells [8, 11]. In addition, CXCL17 likely plays a role in angiogenesis and relevant to tumor development [2,14□20].

The gene of CXCL17 is widely present in mammals from Monotremata to Eutheria (Table S1), but it is not found in lower vertebrates, suggesting it emerged in the early stage of mammalian evolution. CXCL17 is synthesized in vivo as a precursor containing an N-terminal signal peptide. As shown in Fig. S1, most residues of CXCL17 orthologs from different species are variable, but there are six absolutely conserved Cys residues that are expected to form three disulfide bonds. Some C-terminal residues are also conserved (Fig. S1), implying that the C-terminal fragment might be important for CXCL17 function. Despite classification as a chemokine, it is not sure whether CXCL17 share a similar structure with other chemokines, because its three-dimensional structure has not yet been experimentally determined. As predicted by the AlphaFold 2 algorithm, the structure of CXCL17 is different from that of other chemokines (Fig. 1A), but only the C-terminal α-helix has high prediction confidence. Moreover, the algorithm predicts three disulfide bonds, C50□C75, C52□C110, and C77□C103, in human CXCL17 (Fig. 1A).

**Fig. 1.**
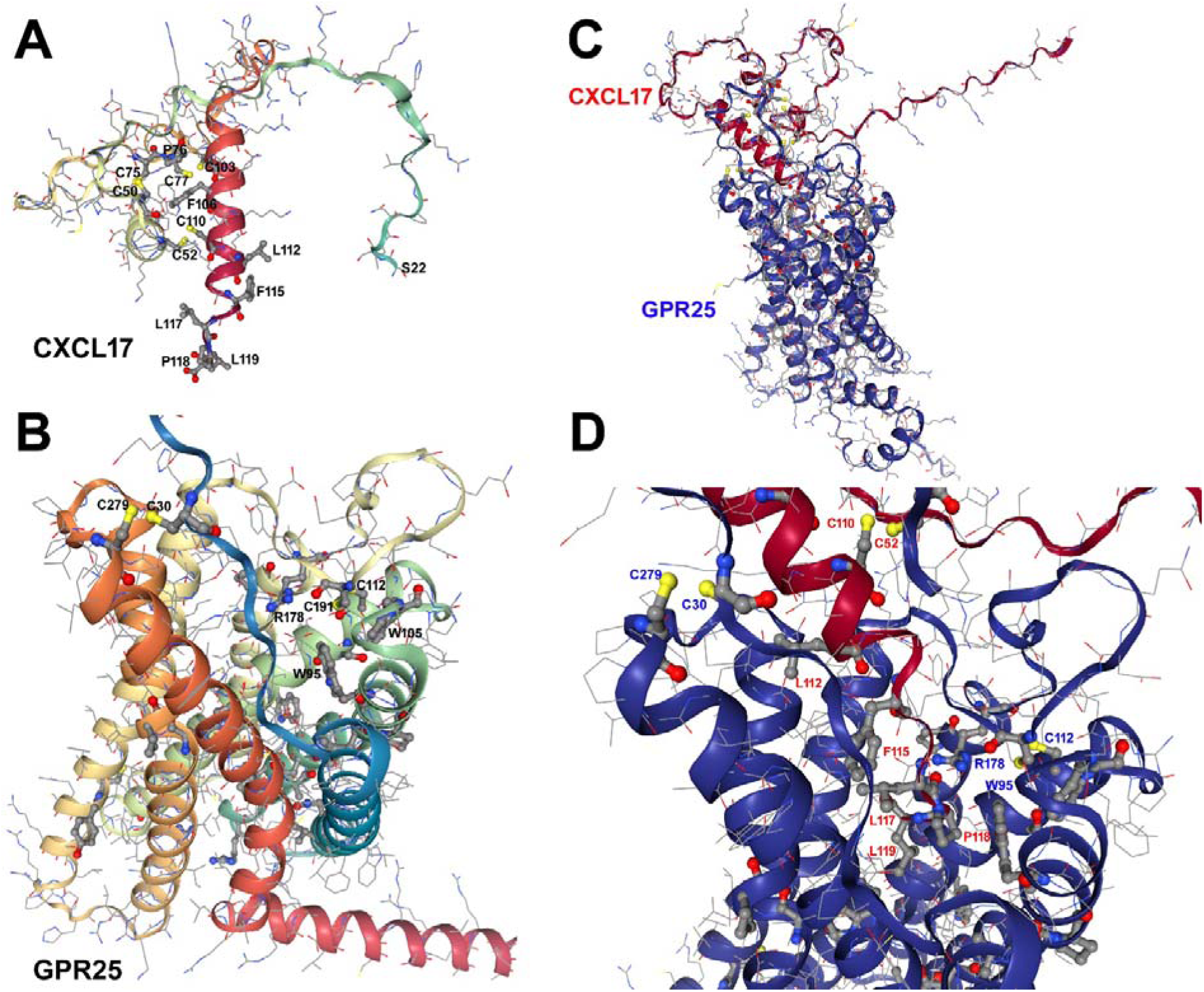
Predicted structures of human CXCL17, human GPR25, and their binding complex. (**A**) Structure of the mature human CXCL17 (residue 22□119) predicted by AlphaFold algorithm. The structure was viewed by an online server (nglviewer.org/ngl) according to the AlphaFold-predicted structure of the full-length human CXCL17 (AF-Q6UXB2-F1-model_v4). The highly conserved residues are shown as balls-and-sticks, and other residues are shown as lines. (**B**) Structure of human GPR25 predicted by AlphaFold algorithm. The structure was viewed by an online server (nglviewer.org/ngl) according to the AlphaFold predicted structure of human GPR25 (AF-O00155-F1-model_v4). The highly conserved residues are shown as balls-and-sticks, and other residues are shown as lines. (**C,D**) The overall structure (C) and close-up view (D) of the binding complex of mature human CXCL17 with human GPR25 predicted by AlphaFold 3. The binding complex was generated by the online AlphaFold 3 server (https://alphafoldserver.com) and viewed by an online server (nglviewer.org/ngl). CXCL17 is show in red and GPR25 in blue.

The function of CXCL17 is expected to be mediated by a plasma membrane receptor, but identity of this receptor is still controversial. The orphan G protein-coupled receptor 35 (GPR35) was first reported as a receptor of CXCL17 in 2015 [21], but this pairing couldn’t be reproduced by other laboratories [22□24]. Recently, two A-class G protein-coupled receptors (GPCRs), the chemokine receptor CXCR4 and the orphan receptor MRGPRX2, were both reported as CXCL17 receptors [24,25]. In another recent paper, CXCL17 was reported to modulate chemokine signaling via binding to extracellular glycosaminoglycans [26].

G protein-coupled receptor 25 (GPR25) is a rarely studied A-class orphan GPCR. Since its discovery in 1997 [27], only six papers can be retrieved from Pubmed using GPR25 as a keyword (https://pubmed.ncbi.nlm.nih.gov). Previous studies suggested that GPR25 might be associated with regulation of blood pressure [28] and expression of PD-1/PD-L1 [29]. According to the data shown at the Human Protein Atlas, GPR25 is predominantly expressed in stomach and lung at tissue level and primarily expressed by T-cells, B-cells, and plasma cells, and some other immune cells at single cell level (https://www.proteinatlas.org/ENSG00000170128-GPR25). Thus, it seems that GPR25 might be responsible for immunity.

GPR25 is widely present in vertebrates, from fishes to mammals (Table S2), and amino acid sequence alignment identified some conserved residues that are likely responsible for structural integrity, downstream signaling, and ligand binding. Predicted by the AlphaFold 2 algorithm, human GPR25 folds into a typical GPCR structure with seven transmembrane domains (TMDs), as well as a short extracellular N-terminus and a short intracellular C-terminus (Fig. 1B). The conserved C112 and C191 form a disulfide bond tethering the extracellular loop 2 to the extracellular end of TMD2 (Fig. 1B). This disulfide bond is present in all GPR25 orthologs (Fig. S2) and most of other GPCRs. C30 and C279 form another disulfide bond tethering the extracellular N-terminal region to the extracellular end of TMD7 (Fig. 1B). This disulfide bond is present in most GPR25 orthologs, but only present in a few of other GPCRs, such as chemokine receptors. The conserved W95, L92, W105, R178, and W257 are located in the expected orthosteric ligand pocket formed by the seven TMDs (Fig. 1B), and might be responsible for binding ligand.

In the present study, we identified the orphan GPR25 as a receptor of CXCL17 via prediction by the newly developed AlphaFold 3 algorithm [30] and subsequent validation by the NanoLuc Binary Technology (NanoBiT)-based β-arrestin recruitment assay in the transfected human embryonic kidney (HEK) 293T cells. Considering their expression pattern, CXCL17 might be an endogenous agonist of GPR25 in human and other mammals, but this hypothesis needs to be tested in future. The present pairing paves the way for functional studies of the rarely studied orphan receptor GPR25 and the orphan chemokine ligand CXCL17.

## Results

### Prediction of possible binding of CXCL17 with GPCRs by AlphaFold 3

We predicted possible binding of mature human CXCL17 with all human chemokine receptors and most of orphan GPCRs using the online AlphaFold 3 server (https://alphafoldserver.com). As shown in Table S3, human CXCL17 binds to human GPR25 with ipTM values of ∼0.7, but it binds to other receptors, including the previously reported CXCL17 receptors (GPR35, MRGPRX2, and CXCR4) with ipTM values always below the confident threshold of 0.6. For some CXCL17□GPR25 pairs from other mammals, AlphaFold 3 prediction also gave ipTM values around 0.6□0.7 (Table S4). Thus, it seems that the orphan GPR25 might be a receptor of CXCL17.

In the binding model predicted by AlphaFold 3, mature CXCL17 binds to the extracellular region of GPR25 with its C-terminus inserting into the receptor’s central hole, the expected orthosteric ligand pocket (Fig. 1C), consistent with the fact that the C-terminal residues of CXCL17 are highly conserved in evolution (Fig. S1). According to the predicted structure (Fig. 1D), the conserved W95 of human GPR25 likely forms hydrophobic interactions with the conserved P118 of human CXCL17, and the positively charged R178 of the receptor likely forms hydrogen bond with the carboxyl oxygen of L117 of the ligand.

### Preparation of mature human CXCL17 peptide

To test whether CXCL17 is a ligand of GPR25 by experiments, we needed mature human CXCL17 peptide that has 98 amino acids including six conserved Cys residues probably forming three disulfide bonds (Fig. S1). To quickly prepare CXCL17, we employed bacterial overexpression using *Escherichia coli* as host cells. For this purpose, we designed a CXCL17 precursor that carries an N-terminal 6×His tag followed by an enterokinase cleavage site (Fig. S3). The 6×His tag will facilitate purification after overexpression, and the enterokinase cleavage site is designed for removal of the tag after in vitro refolding.

After the transformed *E. coli* cells were induced by isopropyl β-D-thiogalactoside (IPTG), a strong band slightly larger than 12 kDa appeared on reducing SDS/PAGE (Fig. 2A), consistent with the theoretical value (13.1 kDa) of the overexpressed CXCL17 precursor. After the bacteria were lysed by sonication, the CXCL17 precursor was mainly present in the pellet as analyzed by SDS/PAGE (Fig. 2A), suggesting it formed inclusion bodies. After solubilized via an *S*-sulfonation approach, the CXCL17 precursor was purified by immobilized metal ion affinity chromatography (Ni^2+^ column). As analyzed by reducing SDS/PAGE (Fig. 2A), the eluted fraction contained a strong band slightly larger than 12 kDa (presumably the monomeric precursor, indicated by an asterisk) and a weak band slightly larger than 30 kDa (presumably the dimeric precursor, indicated by an octothorpe), suggesting that CXCL17 is prone to crosslinking via intermolecular isopeptide bonds. Estimated from the band density, ∼50 mg of the precursor could be typically obtained from 1 L of culture broth after purification by the Ni^2+^ column.

**Fig. 2.**
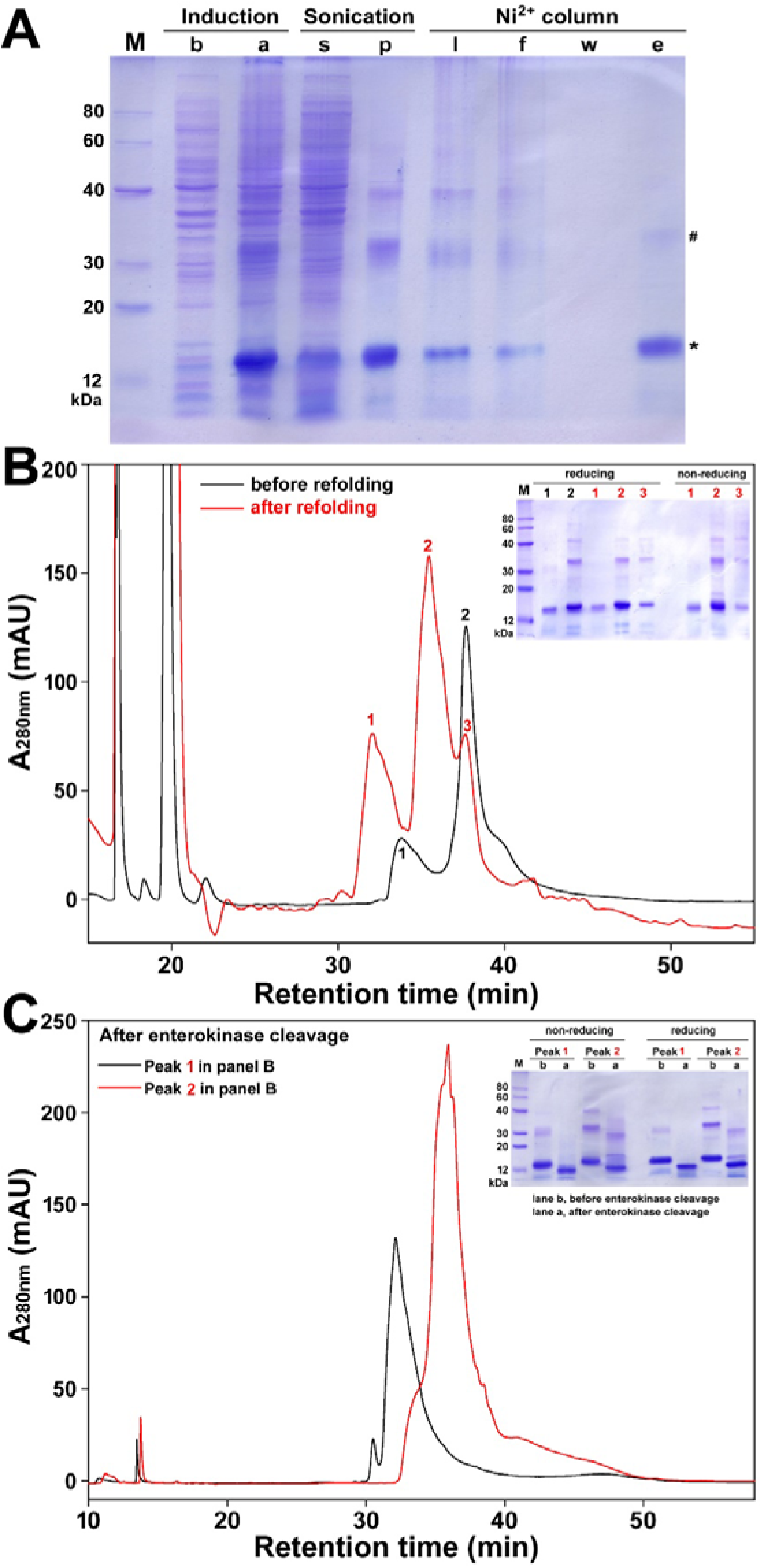
Preparation of the recombinant human CXCL17 peptide. (**A**) SDS/PAGE analysis of the overexpressed CXCL17 precursor at different preparation steps. Lane (M), protein ladder; lane (b), before IPTG induction; lane (a), after IPTG induction; lane (s), supernatant after sonication; lane (p), pallet after sonication; lane (l), sample loading to the Ni^2+^ column after S-sulfonation; lane (f), flow-through faction; lane (w), fraction washed by 10 mM imidazole; lane (e), fraction eluted by 250 mM imidazole. The samples were mixed with SDS loading buffer containing 100 mM DTT and loaded onto a 15% SDS-gel after boiling. After electrophoresis, the gel was stained by Coomassie Brilliant Blue R250. (**B**) HPLC analysis of the CXCL17 precursor before and after in vitro refolding. Black trace, 0.2 mL of the eluent from the Ni^2+^ column was treated by DTT and then loaded on to a C_18_ reverse-phase column; red trace, 0.4 mL of the eluent from the Ni^2+^ column was subjected to in vitro refolding and then loaded on to a C_18_ reverse-phase column. **Inner panel**, SDS/PAGE analysis of the eluted peaks before and after refolding. After lyophilization, samples of these eluted peaks were mixed with SDS loading buffer either containing 100 mM DTT or not, and loaded onto a 15% SDS-gel after boiling. After electrophoresis, the gel was stained by Coomassie Brilliant Blue R250. (**C**) HPLC analysis of the refolded CXCL17 precursor fractions after cleavage by enterokinase. The refolded CXCL17 precursor fractions (peak 1 and peak 2 in panel B) were treated with recombinant enterokinase and then loaded onto a C_18_ reverse-phase column. **Inner panel**, SDS/PAGE analysis before and after enterokinase cleavage. The refolded fractions (peak 1 and 2 after refolding in panel B) before and after enterokinase cleavage were mixed with SDS loading buffer either containing 100 mM DTT or not, and loaded onto a 15% SDS-gel after boiling. After electrophoresis, the gel was stained by Coomassie Brilliant Blue R250.

The CXCL17 precursor eluted from the Ni^2+^ column was further analyzed by high performance liquid chromatography (HPLC) after removal of the reversibly modified sulfonate moieties by dithiothreitol (DTT) treatment: two peaks were eluted from a C_18_ reverse-phase column by an acidic acetonitrile gradient (Fig. 2B, black trace). As analyzed by reducing SDS/PAGE (Fig. 2B, inner panel), the higher peak 2 contained CXCL17 monomer as well as small amount of dimer and trimer; while the lower peak 1 only contained CXCL17 monomer, but it seemed slightly smaller than the monomer in peak 2. Thus, it seemed that the peak 1 may be partially degraded products of the overexpressed CXCL17 precursor.

After in vitro refolding, three peaks were eluted from the C_18_ reverse-phase column (Fig. B). The peak 1 was likely the refolding product of the partially degraded CXCL17 precursor, because it was slightly smaller than the monomer of peak 2 as analyzed by SDS/PAGE either under reducing condition or non-reducing condition (Fig. 2B, inner panel). The largest peak 2 and the smallest peak 3 were likely the refolding products of the intact CXCL17 precursor, but both peaks contained some dimmers and trimers as analyzed by SDS/PAGE either under reducing condition or under non-reducing condition (Fig. 2B, inner panel), suggesting these oligomers are crosslinked via intermolecular isopeptide bonds.

After in vitro refolding, the refolded peak 1 and peak 2 shown in Fig. 2B were subjected to enterokinase cleavage to remove the N-terminal tag of the precursor. As analyzed by HPLC (Fig. 2C), a broad major peak was eluted from the C_18_ reverse-phase column for each fraction after enterokinase treatment. As analyzed by SDS/PAGE, enterokinase treatment significantly decreased the molecular weight of both fractions (Fig. 2C, inner panel), implying removal of their N-terminal tag. After enterokinase treatment, the peak 1 was also slightly smaller than peak 2 on SDS/PAGE either under reducing condition or non-reducing condition (Fig. 2C, inner panel), suggesting the conserved C-terminus of peak 1 might be truncated when overexpressed in *E. coli*. Thus, only the correct peak 2 was used for later activity assay after enterokinase cleavage.

To test importance of the conserved C-terminus, we prepared a truncated mutant peptide, designated as [desC3]CXCL17, in which three C-terminal residues of human CXCL17 were removed (Fig. S3). The mutant peptide was prepared using the same procedure for preparation of the wild-type (WT) CXCL17 peptide. This mutant precursor had only one elution peak on HPLC before and after in vitro refolding.

### CXCL17 activates GPR25 in **β**-arrestin recruitment assays

If CXCL17 functions as an agonist for GPR25, it would activate this receptor and cause β-arrestin recruitment, a general phenomenon for GPCRs [31,32]. Thus, we developed a β-arrestin recruitment assay for human GPR25 based on the bioluminescent NanoBiT approach [33], that has been used to test reported possible ligands of GPR83 and S1PR2 in our recent studies [34□36]. A large NanoLuc fragment (LgBiT) was genetically fused to the intracellular C-terminus of human GPR25 and a low-affinity SmBiT complementation tag was genetically fused to the N-terminus of human β-arrestin 2 (ARRB2) or human β-arrestin 1 (ARRB1). The resultant GPR25-LgBiT and the SmBiT-fused β-arrestin were coexpressed in HEK293T cells under a controllable manner via a doxycycline (Dox)-response bi-directional promoter. If CXCL17 is an agonist, its binding would activate GPR25-LgBiT and induce binding of SmBiT-fused β-arrestin, thus proximity effect would cause complementation of the β-arrestin-fused low-affinity SmBiT tag with the receptor-fused LgBiT and restore NanoLuc luciferase activity. This NanoBiT-based β-arrestin recruitment assay was unlikely affected by endogenously expressed CXCL17 receptors, because they have no LgBiT fusion.

After the transfected HEK293T cells were induced to coexpress GPR25-LgBiT and SmBiT-ARRB2, low bioluminescence was detected after addition of the NanoLuc substrate (Fig. 3A), confirming coexpression of GPR25-LgBiT and SmBiT-ARRB2 and their background complementation. After addition of the recombinant WT CXCL17 peptide, the measured bioluminescence increased quickly in a dose-dependent manner (Fig. 3A). According to the peak bioluminescence after addition of CXCL17, a roughly sigmoidal activation curve was obtained with an EC_50_ value of ∼70 nM (Fig. 3B, inner panel). Thus, it seemed that CXCL17 can activate GPR25 and cause recruitment of β-arrestin 2.

**Fig. 3.**
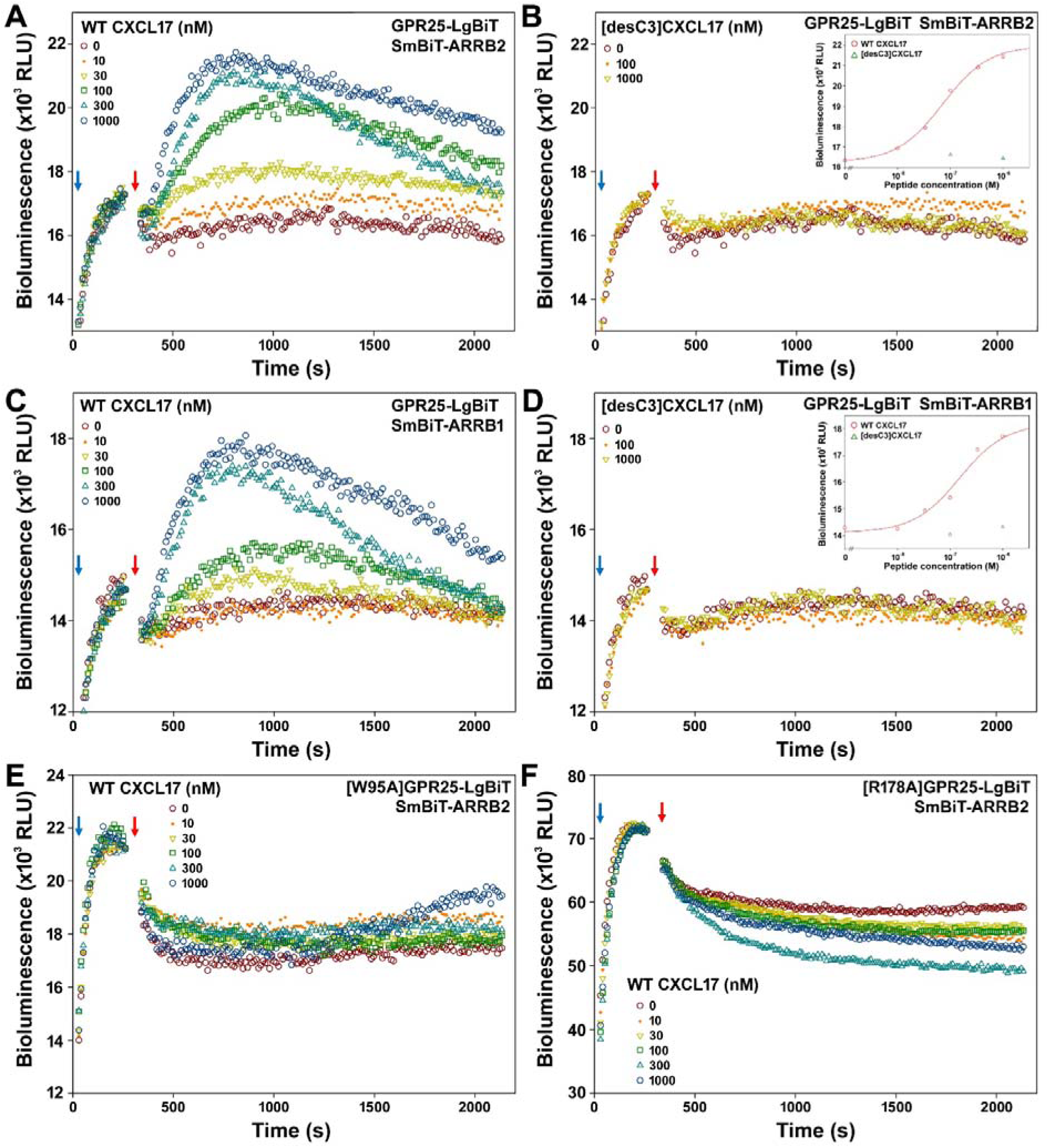
CXCL17 activates GPR25 in NanoBiT-based β-arrestin recruitment assays. (**A**) A typical bioluminescence change after sequential addition of NanoLuc substrate and WT CXCL17 to living HEK293T cells coexpressing GPR25-LgBiT and SmBiT-ARRB2. (**B**) A typical bioluminescence change after sequential addition of NanoLuc substrate and [desC3]CXCL17 to living HEK293T cells coexpressing GPR25-LgBiT and SmBiT-ARRB2. Inner panel, dose curve of WT or mutant CXCL17 activating GPR25 assayed as β-arrestin 2 recruitment. The peak bioluminescence data in panel A and B are plotted with peptide concentrations using SigmaPlot 10.0 software. (**C**) A typical bioluminescence change after sequential addition of NanoLuc substrate and WT CXCL17 to living HEK293T cells coexpressing GPR25-LgBiT and SmBiT-ARRB1. (**D**) A typical bioluminescence change after sequential addition of NanoLuc substrate and [desC3]CXCL17 to living HEK293T cells coexpressing GPR25-LgBiT and SmBiT-ARRB1. Inner panel, dose curve of WT or mutant CXCL17 activating GPR25 assayed as β-arrestin 1 recruitment. The peak bioluminescence data in panel C and D are plotted with peptide concentrations using SigmaPlot 10.0 software. (**E**) A typical bioluminescence change after sequential addition of NanoLuc substrate and WT CXCL17 to living HEK293T cells coexpressing [W95A]GPR25-LgBiT and SmBiT-ARRB2. (**F**) A typical bioluminescence change after sequential addition of NanoLuc substrate and WT CXCL17 to living HEK293T cells coexpressing [R178A]GPR25-LgBiT and SmBiT-ARRB2. In these panels, blue arrows indicate addition of NanoLuc substrate and red arrows indicate addition of peptide.

After three C-terminal residues were deleted, the resultant [desC3]CXCL17 mutant peptide had no detectable activation effect on human GPR25 in the β-arrestin 2 recruitment assay (Fig. 3B). Thus, it seemed that the C-terminal fragment is important for CXCL17 activating GPR25, consistent with the fact that C-terminal residues are highly conserved among CXCL17 orthologs from different species (Fig. S1).

Towards living HEK293T cells coexpressing GPR25-LgBiT and SmBiT-ARRB1, addition of WT CXCL17 also caused significant increase of bioluminescence in a dose-dependent manner (Fig. 3C), with a calculated EC_50_ value of ∼150 nM (Fig. 3D, inner panel). Deletion of three C-terminal residues also abolished the activity of CXCL17 on GPR25 in the β-arrestin 1 recruitment assay (Fig. 3D). Thus, it seemed that CXCL17 can activate GPR25 and cause recruitment of both β-arrestin 2 and β-arrestin 1 in the transfected HEK293T cells.

After the large aromatic W95 or the positively charged R178, two conserved residues in the expected orthosteric ligand binding pocket of human GPR25 (Fig. 1D), was replaced by an alanine residue, WT CXCL17 almost had no activation effect on both mutant receptors in the β-arrestin recruitment assay (Fig. 3E,F). Thus, both W95 and R178 seemed important for GPR25 interacting with CXCL17, consistent with the predicted binding model in which W95 and R178 contact with C-terminal residues of CXCL17.

### CXCL17 has no activation effects on other tested GPCRs

As shown above, CXCL17 could activate GPR25 in the NanoBiT-based β-arrestin recruitment assays and this activation effect relied on three ligand residues at C-terminus and two receptor residues (W95 and R178) in the expected orthosteric ligand binding pocket. To further test whether CXCL17 is specific to GPR25, we measured its activation effect on 17 other GPCRs using the NanoBiT-based β-arrestin recruitment assay (Fig. 4). These human GPCRs were genetically fused with a LgBiT at their intracellular C-terminus and coexpressed with an N-terminally SmBiT-fused human β-arrestin 2 (ARRB2) via a Dox-response bi-directional promoter in transfected HEK293T cells.

**Fig. 4.**
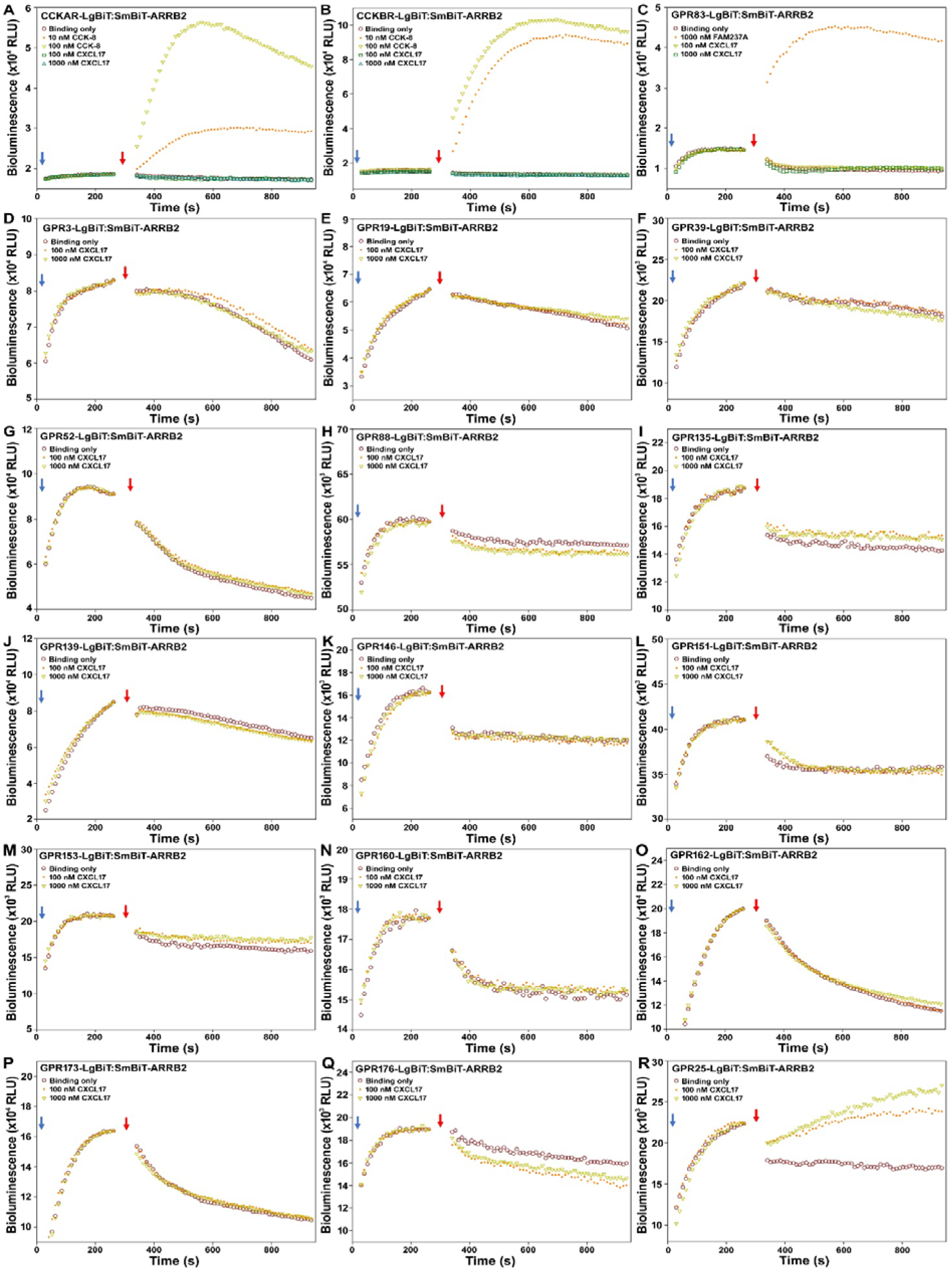
CXCL17 has no activation effects on other tested GPCRs. In the NanoBiT-based β-arrestin recruitment assays, NanoLuc substrate and indicated peptides were sequentially added to living HEK293T cells coexpressing a C-terminally LgBiT-fused GPCR and an N-terminally SmBiT-fused human β-arrestin 2 (ARRB2). The blue arrows indicate addition of NanoLuc substrate and red arrows indicate addition of peptide.

Towards CCKAR and CCKBR, two receptors of the peptide hormone cholecystokinin (CCK), addition of their cognate agonist CCK-8 peptide caused quick increase of bioluminescence in a dose-dependent manner (Fig. 4A,B). In contrast, up to 1.0 μM of recombinant WT CXCL17 had no detectable effect on both receptors (Fig. 4A,B). Towards the officially orphan receptor GPR83, addition of the recombinant FAM237A protein, a recently discovered agonist [34,35,37], caused quick increase of bioluminescence, but addition of WT CXCL17 had no detectable effect (Fig. 4C). Thus, these receptors can be normally activated by their cognate agonist, but had no response to CXCL17.

Towards 14 other orphan GPCRs, including GPR3, GPR19, GPR39, GPR52, GPR88, GPR135, GPR139, GPR146, GPR151, GPR153, GPR160, GPR162, GPR173 and GPR176, addition of WT CXCL17 had no significant effect on the measured bioluminescence (Fig. 4D□Q), implying that CXCL17 cannot activate these receptors. When assayed these orphan receptors, GPR25 was used as a positive control (Fig. 4R): significant bioluminescence increase was observed after addition of WT CXCL17. Thus, it seemed that CXCL17 specifically activates the orphan receptor GPR25.

## Discussion

In the present study, the newly developed AlphaFold 3 algorithm [30] was employed to identify possible receptor for CXCL17. This is the first report for receptor deorphanization using the AlphaFold 3 algorithm. As a powerful tool for predicting protein[protein interactions, the AlphaFold 3 algorithm will significantly accelerate deorphanization of receptors and peptides in future studies. After prediction, we experimentally demonstrated that the recombinant human CXCL17 could specifically activate human GPR25 in transfected HEK293T cells and their interaction relies on three conserved residues at the C-terminus of the ligand and two conserved residues (W95 and R178) in the expected orthosteric ligand binding pocket of the receptor. The experimental results are consistent with the predicted binding model in which the conserved C-terminus of CXCL17 inserts into the expected orthosteric ligand binding pocket of the receptor GPR25.

GPR25 is a rarely studied orphan receptor and its biological functions are largely unknown. No natural or synthetic ligands (agonists or antagonists) have yet been identified for this receptor. Lack of known ligands seriously hampered its functional studies in the past. To the best of our knowledge, CXCL17 is the so far only known ligand (agonist) for this receptor. Thus, CXCL17 or its analogs are valuable tools to study the biological functions of GPR25 in future. GPR25 is likely responsible for immunity and might be a therapeutic target for some diseases, our present pairing opened the door for characterization of this rarely studied receptor.

CXCL17 has been reported to activate some other receptors. For an example, it was reported to activate GPR35 in 2015 based on the observation that overexpression of GPR35 in BA/F3 cells or HEK293 cells could increase the response of these cells to CXCL17 treatment in calcium mobilization assay [21]. However, this result couldn’t be reproduced by three other laboratories [22□24]. At present stage, it is quite sure that GPR35 is not a receptor of CXCL17. Recently, CXCL17 was reported to activate the orphan receptor MRGPRX2, but quite high concentration (typically 300 nM) of CXCL17 was needed [25]. As an orphan MAS-related receptor, MRGPRX2 can be activated by a variety of peptides and compounds at micromolar concentrations [38]. Another recent study reported that CXCL17 is an allosteric inhibitor of the chemokine receptor CXCR4 signaling [24]. In future, more experiments are needed to confirm the real receptor of CXCL17 using various assays, since independent confirmation is a minimum criterion for ligand[receptor pairing [39,40].

According to the data shown at the Human Protein Atlas, CXCL17 is primarily expressed in lung by club cells, basal respiratory cells, and alveolar cells, in stomach by gastric mucus-secreting cells, in esophagus by squamous epithelial cells, and in salivary gland by serous glandular cells; while GPR25 is primarily expressed by T-cells, B-cells, plasma cells, and some other immune cells. Thus, these mucosal tissues might affect the function of these GPR25-expressing immune cells via their secreted CXCL17. However, more studies are needed to test whether CXCL17 is an endogenous ligand of GPR25 in future.

Our recombinant human CXCL17 displayed EC_50_ values at ∼100 nM in our assays, that seems too high for an endogenous ligand. Two aspects might be considered in future work. (a) Improving quality of the recombinant CXCL17 peptide. Mature CXCL17 probably has three disulfide bonds and a highly flexible structure as predicted by AlphaFold (Fig. 1A). After the bacteria overexpressed CXCL17 precursor was subjected to in vitro refolding, we always obtained a broad peak on HPLC, implying the peptide might fold into several disulfide isomers. SDS-PAGE analysis indicated that CXCL17 is prone to formation of dimmers and trimers via intermolecular isopeptide bonds. In future, other expression systems, such as yeast or mammalian cells, will be used to produce high quality CXCL17 peptide for activity assays. (b) Is the CXCL17 peptide further processed? In the present work, we used the full-length mature human CXCL17 (amino acids 22□118) for activity assays. However, some previous studies suggested that CXCL17 might be further processed in cells or after secretion. Lee et al. reported that CXCL17 might be cleaved at 61□63 position when synthesized in mammalian cells [18]. The secreted CXCL17 might also be further processed by extracellular chymase, like some other chemokines [23]. These possibilities will also be tested in future studies. CXCL17 might display higher activity towards GPR25 after high quality peptide or correct isoforms was used for activity assays.

The gene of CXCL17 is widely present in extant mammals, but absent from extant lower vertebrates, such as birds, reptiles, amphibians, and fishes, implying that it originated in the early stage of mammalian evolution. However, the gene GPR25 is widely present in extant vertebrates, from fishes to mammals, implying that it emerged in ancient fishes and then spread to all vertebrate lineages. Thus, GPR25 has a much earlier origination than CXCL17. It is possible that the later emerged CXCL17 used an existent receptor to mediate its function when it originated in ancient mammals. However, GPR25 might have other ligand(s) that has earlier origination than CXCL17. This possibility needs to be tested in future studies. For searching other ligands for GPR25, CXCL17 may be used as a positive control in various screening assays.

## Materials and methods

### Preparation of mature human CXCL17

The nucleotide sequence encoding the designed CXCL17 precursor (6×His-CXCL17) was chemically synthesized at Tsingke Biotechnology (Beijing, China) using the *E. coli*-biased codons (Fig. S3). After cleavage with restriction enzymes NdeI and EcoRI, the synthetic DNA fragment was ligated into a pET vector, resulting in the bacteria expression construct pET/6×His-CXCL17. The expression construct pET/6×His-[desC3]CXCL17, encoding the C-terminally truncated mutant precursor, was generated by site-directed mutagenesis via the QuikChange approach .

Thereafter, the wild-type or mutant CXCL17 precursors were overexpressed in *E. coli* strain BL21(DE3) according to standard protocols. The transformed bacteria were induced by 1.0 mM IPTG at 37 °C for 6−8 h, harvested by centrifugation (5000 *g*, 10 min), re-suspended in lysis buffer (20 mM phosphate, pH 7.4, 0.5 M NaCl), and lysed by sonication. Thereafter, inclusion bodies were collected via centrifugation (15000 *g*, 30 min), resuspended in solubilizing buffer (lysis buffer plus 6 M guanidine chloride), and subjected to *S*-sulfonation by addition of sodium sulfite (Na_2_SO_3_) and sodium tetrathionate (Na_2_S_4_O_6_) to the final concentration of 100 mM and 80 mM, respectively. After overnight shaking at room temperature, the supernatant was applied to an immobilized metal ion affinity chromatography, and the *S*-sulfonated CXCL17 precursor was eluted by 250 mM imidazole (in solubilizing buffer) from the Ni^2+^ column. The samples at each step were analyzed by reducing SDS/PAGE.

Thereafter, the *S*-sulfonated wild-type or mutant CXCL17 precursors were subjected to in vitro refolding via a two-step procedure. The *S*-sulfonated precursor fraction eluted from Ni^2+^ column was first treated with 25 mM DTT at 37 °C for 1 h, and then 25-fold diluted into ice-cold refolding buffer (0.5 M L-arginine, pH 7.5) containing 2.0 mM oxidized glutathione. After overnight incubation on ice, the refolding mixture was acidified to pH3−4 by adding trifluoroacetic acid (TFA) and subjected to high performance liquid chromatography (HPLC). The refolded precursor was eluted from a C_18_ reverse-phase column (Zorbax 300SB-C8, 9.4 × 250 mm, Agilent Technologies, Santa Clara, CA, USA) by an acidic acetonitrile gradient and manually collected. After lyophilization, the refolded precursor fractions were dissolved in 1.0 mM aqueous hydrochloride (pH3.0) and quantified by ultra-violet absorbance at 280 nm using the extinction coefficient of ε_280 nm_ = 11000 M^-1^ cm^-1^.

To remove the N-terminal tag, the refolded CXCL17 precursor was diluted into the cleavage buffer (25 mM Tris-HCl, pH 8.0, 50 mM NaCl, 2 mM CaCl_2_) to the final concentration of ∼2 mg/mL. Thereafter, recombinant enterokinase (Yaxin Bio, Shanghai, China) was added to the final concentration of ∼4 U/mL to start cleavage. After incubation at 25 °C for ∼1.5 h, the reaction mixture was acidified to pH3□4 using TFA and applied to HPLC. The mature CXCL17 peptide was eluted from a C_18_ reverse-phase column (Zorbax 300SB-C8, 9.4 × 250 mm, Agilent Technologies) by an acidic acetonitrile gradient and manually collected. After lyophilization, mature WT or mutant CXCL17 fractions were dissolved in 1.0 mM aqueous hydrochloride (pH3.0), quantified by ultra-violet absorbance at 280 nm using the extinction coefficient of ε_280 nm_ = 11000 M^-1^ cm^-1^, and used for activity assay.

### Generation of the expression constructs for human GPR25

The coding region of human GPR25 was amplified by polymerase chain reaction (PCR) using human genomic DNA (extracted from HEK293T cells) as template and designed oligo primers listed in Table S5. After cleavage with restriction enzymes NheI and AgeI, the amplified DNA fragment was ligated into a pcDNA6 vector (Table S5), resulting in the expression construct pcDNA6/GPR25 that encodes an untagged human GPR25 (Fig. S4) whose nucleotide sequencing is identical to the reference sequence (NM_005298). Thereafter, the construct pcDNA6/GPR25 was used as template for PCR amplification using appropriate primers, and the amplified DNA fragments were then cloned into appropriate vectors via Gibson assembly or ligation by T4 DNA ligase (Table S5). The constructs pTRE3G-BI/GPR25-LgBiT:SmBiT-ARRB2 and pTRE3G-BI/GPR25-LgBiT:SmBiT-ARRB1 coexpress a C-terminally LgBiT-fused human GPR25 (GPR25-LgBiT) and an N-terminally SmBiT-fused human β-arrestin 2 (SmBiT-ARRB2) or human β-arrestin 1 (SmBiT-ARRB1) under a controllable manner via a Dox-response bidirectional promoter. The expression constructs for [W95A]GPR25 or [R178A]GPR25 were generated by PCR amplification and Gibson assembly as shown in Table S5. After cloning, the constructs were confirmed by DNA sequencing.

### **β**-arrestin recruitment assays

HEK293T cells (RRID: CVCL_0063) were from Shanghai Zhong Qiao Xin Zhou Biotechnology Co., Ltd. (Shanghai, China). The cell line was mycoplasma-free and authenticated using STR profiling within the last 3□years. The cells were cultured in complete medium (DMEM plus 10% fetal bovine serum and antibiotics) according to standard protocols. To conduct the β-arrestin recruitment assay, the coexpression construct pTRE3G-BI/GPR25-LgBiT:SmBiT-ARRB1, pTRE3G-BI/GPR25-LgBiT:SmBiT-ARRB2, or those encoding GPR25 mutants or other GPCRs was transiently transfected into HEK293T cells together with the expression control vector pCMV-TRE3G (Clontech, Mountain View, CA, USA) using the transfection reagent Lipo8000 (Beyotime Technology, Shanghai, China). Next day, the transfected cells were trypsinized, suspended in the induction medium (complete DMEM medium plus 1.0□10 ng/ml of Dox), seeded into white opaque 96-well plates, and cultured at 37 °C for ∼24 h. To start the assay, the medium was removed and pre-warmed activation solution (serum-free DMEM plus 1% BSA) containing NanoLuc substrate was added (40 μL/well, containing 1.0 μL of NanoLuc substrate stock from Promega, Madison, WI, USA), and bioluminescence data were immediately collected for ∼4 min on a SpectraMax iD3 plate reader (Molecular Devices, Sunnyvale, CA, USA) with an interval of 11 s. Subsequently, WT or mutant CXCL17 peptide (diluted in the activation solution) was added (10 μL/well), and bioluminescence data were continuously collected for ∼30 min or shorter time with an interval of 11 s. The measured absolute signals were corrected for inter well variability by forcing all curves after addition of NanoLuc substrate (without ligand) to same level and plotted using the SigmaPlot 10.0 software (SYSTAT software, Chicago, IL, USA). To obtain the dose curve for CXCL17 activating GPR25, average values of peak bioluminescence (715□935 s) after addition of CXCL17 were plotted with peptide concentrations using the SigmaPlot 10.0 software (SYSTAT software).

## Abbreviations

BSA: bovine serum albumin
CXCL17: C-X-C motif chemokine ligand 17
CRE: cAMP response element
DMEM: Dulbecco’s modified Eagle medium
Dox: doxycycline
GPCR: G protein-coupled receptor
GPR25: G protein-coupled receptor 25
HEK: human embryonic kidney
HPLC: high performance liquid chromatography
IPTG: isopropyl β-D-thiogalactoside
LgBiT: a large NanoLuc fragment for NanoBiT
NanoBiT: NanoLuc Binary Technology
NanoLuc: Nanoluciferase
PCR: polymerase chain reaction
SD: standard deviation
SDS/PAGE: sodium dodecyl sulfate/polyacrylamide gel electrophoresis
sLuc: secretory Nanoluciferase
SmBiT: a low-affinity complementation tag for NanoBiT
TFA: trifluoroacetic acid
TGFα: transforming growth factor α
TMD: transmembrane domain
WT: wild-type.

## Author Contributions

YFW, JY, JJW, RJS, YSZ, and TZ performed experiments; YLL and ZGX analyzed data; ZYG planned experiments and wrote the paper.

## Acknowledgments

We thank some former laboratory members, Ben-Jun Ji, Yu Liu, Ge Song, Lei Zhang, Yu-Qi Guo, Qing-Ping Wu, Li-Li Shou, and Ning Li, for generation of over 200 GPCR expression constructs. This work was supported by grants from the National Natural Science Foundation of China (31470767, 31971193).

## Data availability statement

All data generated or analyzed during this study are included in the published article.

## Supporting information

**Table S1.** Accession numbers of CXCL17 orthologs aligned in Fig. S1.

**Table S2.** Accession numbers of GPR25 orthologs aligned in Fig. S2.

**Table S3.** Possible binding of human CXCL17 with human GPCRs predicted by AlphaFold 3.

**Table S4.** AlphaFold 3-predicted possible binding of CXCL17□GPR25 pairs from different species.

**Table S5.** Primers and vectors used for generation of human GPR25 expression constructs.

**Fig. S1.** Amino acid sequence alignment of CXCL17 orthologs.

**Fig. S2.** Amino acid sequence alignment of GPR25 orthologs.

**Fig. S3.** The nucleotide and amino acid sequences of the designed CXCL17 precursor.

**Fig. S4.** The nucleotide and amino acid sequences of human GPR25 constructs.

**Fig. S5.** The nucleotide and amino acid sequences of the sLuc-TGFα reporter.

## References

1 Pisabarro MT, Leung B, Kwong M, Corpuz R, Frantz GD, Chiang N, Vandlen R, Diehl LJ, Skelton N, Kim HS, Eaton D & Schmidt KN (2006) Cutting edge: novel human dendritic cell- and monocyte-attracting chemokine-like protein identified by fold recognition methods. J Immunol 176, 2069□2073.

2 Weinstein EJ, Head R, Griggs DW, Sun D, Evans RJ, Swearingen ML, Westlin MM & Mazzarella R (2006) VCC-1, a novel chemokine, promotes tumor growth. Biochem Biophys Res Commun 350, 74□81.

3 Choreño-Parra JA, Thirunavukkarasu S, Zúñiga J & Khader SA (2020) The protective and pathogenic roles of CXCL17 in human health and disease: Potential in respiratory medicine. Cytokine Growth Factor Rev 53, 53□62.

4 Xiao S, Xie W & Zhou L (2021) Mucosal chemokine CXCL17: What is known and not known. Scand J Immunol 93, e12965.

5 Denisov SS (2021) CXCL17: The Black Sheep in the Chemokine Flock. Front Immunol 12, 712897.

6 Giblin SP & Pease JE (2023) What defines a chemokine? - The curious case of CXCL17. Cytokine 168, 156224.

7 Burkhardt AM, Tai KP, Flores-Guiterrez JP, Vilches-Cisneros N, Kamdar K, Barbosa-Quintana O, Valle-Rios R, Hevezi PA, Zuñiga J, Selman M, Ouellette AJ & Zlotnik A (2012) CXCL17 is a mucosal chemokine elevated in idiopathic pulmonary fibrosis that exhibits broad antimicrobial activity. J Immunol 188, 6399□6406.

8 Burkhardt AM, Maravillas-Montero JL, Carnevale CD, Vilches-Cisneros N, Flores JP, Hevezi PA & Zlotnik A (2014) CXCL17 is a major chemotactic factor for lung macrophages. J Immunol 193, 1468□1474.

9 Oka T, Sugaya M, Takahashi N, Takahashi T, Shibata S, Miyagaki T, Asano Y & Sato S (2017) CXCL17 Attenuates Imiquimod-Induced Psoriasis-like Skin Inflammation by Recruiting Myeloid-Derived Suppressor Cells and Regulatory T Cells. J Immunol 198, 3897□3908.

10 Srivastava R, Hernández-Ruiz M, Khan AA, Fouladi MA, Kim GJ, Ly VT, Yamada T, Lam C, Sarain SAB, Boldbaatar U, Zlotnik A, Bahraoui E & BenMohamed L (2018) CXCL17 Chemokine-Dependent Mobilization of CXCR8(+)CD8(+) Effector Memory and Tissue-Resident Memory T Cells in the Vaginal Mucosa Is Associated with Protection against Genital Herpes. J Immunol 200, 2915□2926.

11 Hernández-Ruiz M, Othy S, Herrera C, Nguyen HT, Arrevillaga-Boni G, Catalan-Dibene J, Cahalan MD & Zlotnik A (2019) Cxcl17(-/-) mice develop exacerbated disease in a T cell-dependent autoimmune model. J Leukoc Biol 105, 1027□1039.

12 Lowry E, Chellappa RC, Penaranda B, Sawant KV, Wakamiya M, Garofalo RP & Rajarathnam K (2024) CXCL17 is a proinflammatory chemokine and promotes neutrophil trafficking. J Leukoc Biol 115, 1177□1182.

13 Yin Y, Mu C, Wang J, Wang Y, Hu W, Zhu W, Yu X, Hao W, Zheng Y, Li Q & Han W (2023) CXCL17 Attenuates Diesel Exhaust Emissions Exposure-Induced Lung Damage by Regulating Macrophage Function. Toxics 11, 646.

14 Mu X, Chen Y, Wang S, Huang X, Pan H & Li M (2009) Overexpression of VCC-1 gene in human hepatocellular carcinoma cells promotes cell proliferation and invasion. Acta Biochim Biophys Sin (Shanghai) 41, 631□637.

15 Hiraoka N, Yamazaki-Itoh R, Ino Y, Mizuguchi Y, Yamada T, Hirohashi S & Kanai Y (2011) CXCL17 and ICAM2 are associated with a potential anti-tumor immune response in early intraepithelial stages of human pancreatic carcinogenesis. Gastroenterology 140, 310□121.

16 Zhou Z, Lu X, Zhu P, Zhu W, Mu X, Qu R & Li M (2012) VCC-1 over-expression inhibits cisplatin-induced apoptosis in HepG2 cells. Biochem Biophys Res Commun 420, 336□342.

17 Matsui A, Yokoo H, Negishi Y, Endo-Takahashi Y, Chun NA, Kadouchi I, Suzuki R, Maruyama K, Aramaki Y, Semba K, Kobayashi E, Takahashi M & Murakami T (2012) CXCL17 expression by tumor cells recruits CD11b+Gr1 high F4/80- cells and promotes tumor progression. PLoS One 7, e44080.

18 Lee WY, Wang CJ, Lin TY, Hsiao CL & Luo CW (2013) CXCL17, an orphan chemokine, acts as a novel angiogenic and anti-inflammatory factor. Am J Physiol Endocrinol Metab 304, E32□40.

19 Li L, Yan J, Xu J, Liu CQ, Zhen ZJ, Chen HW, Ji Y, Wu ZP, Hu JY, Zheng L & Lau WY (2014) CXCL17 expression predicts poor prognosis and correlates with adverse immune infiltration in hepatocellular carcinoma. PLoS One 9, e110064.

20 Koni E, Congur I & Tokcaer Keskin Z (2024) Overexpression of CXCL17 increases migration and invasion of A549 lung adenocarcinoma cells. Front Pharmacol 15, 1306273.

21 Maravillas-Montero JL, Burkhardt AM, Hevezi PA, Carnevale CD, Smit MJ & Zlotnik A (2015) Cutting edge: GPR35/CXCR8 is the receptor of the mucosal chemokine CXCL17. J Immunol 194, 29□33.

22 Park SJ, Lee SJ, Nam SY & Im DS (2018) GPR35 mediates lodoxamide-induced migration inhibitory response but not CXCL17-induced migration stimulatory response in THP-1 cells; is GPR35 a receptor for CXCL17? Br J Pharmacol 175, 154□161.

23. Binti Mohd Amir NAS, Mackenzie AE, Jenkins L, Boustani K, Hillier MC, Tsuchiya T, Milligan G & Pease JE (2018) Evidence for the Existence of a CXCL17 Receptor Distinct from GPR35. J Immunol 201, 714□724.

24 White CW, Platt S, Kilpatrick LE, Dale N, Abhayawardana RS, Dekkers S, Kindon ND, Kellam B, Stocks MJ, Pfleger KDG & Hill SJ (2024) CXCL17 is an allosteric inhibitor of CXCR4 through a mechanism of action involving glycosaminoglycans. Sci Signal 17, eabl3758.

25 Ding J, Hillig C, White CW, Fernandopulle NA, Anderton H, Kern JS, Menden MP & Mackay GA (2024) CXCL17 induces activation of human mast cells via MRGPRX2. Allergy 79, 1609□1612.

26 Giblin SP, Ranawana S, Hassibi S, Birchenough HL, Mincham KT, Snelgrove RJ, Tsuchiya T, Kanegasaki S, Dyer D & Pease JE (2023) CXCL17 binds efficaciously to glycosaminoglycans with the potential to modulate chemokine signaling. Front Immunol 14, 1254697.

27 Jung BP, Nguyen T, Kolakowski LF Jr, Lynch KR, Heng HH, George SR & O’Dowd BF (1997) Discovery of a novel human G protein-coupled receptor gene (GPR25) located on chromosome 1. Biochem Biophys Res Commun 230, 69□72.

28 Sherva R, Miller MB, Lynch AI, Devereux RB, Rao DC, Oberman A, Hopkins PN, Kitzman DW, Atwood LD & Arnett DK (2007) A whole genome scan for pulse pressure/stroke volume ratio in African Americans: the HyperGEN study. Am J Hypertens 20, 398□402.

29 Chen Z, Chen Q, Li S, Tu S, Chen Q & Wang A (2022) *IL-12RB1*: a novel immune prognostic biomarker for oral squamous cell carcinoma and linked to *PD-1/PD-L1* expression in the tumor immune microenvironment. Ann Transl Med 10, 144.

30 Abramson J, Adler J, Dunger J, Evans R, Green T, Pritzel A, Ronneberger O, Willmore L, Ballard AJ, Bambrick J, Bodenstein SW, Evans DA, Hung CC, O’Neill M, Reiman D, Tunyasuvunakool K, Wu Z, Žemgulytė A, Arvaniti E, Beattie C, Bertolli O, Bridgland A, Cherepanov A, Congreve M, Cowen-Rivers AI, Cowie A, Figurnov M, Fuchs FB, Gladman H, Jain R, Khan YA, Low CMR, Perlin K, Potapenko A, Savy P, Singh S, Stecula A, Thillaisundaram A, Tong C, Yakneen S, Zhong ED, Zielinski M, Žídek A, Bapst V, Kohli P, Jaderberg M, Hassabis D & Jumper JM (2024) Accurate structure prediction of biomolecular interactions with AlphaFold 3. Nature 630, 493□500.

31 Qi M, Chen TT, Li L, Gao PP, Li N, Zhang SH, Wei W & Sun WY (2024) Insight into the regulatory mechanism of β-arrestin2 and its emerging role in diseases. Br J Pharmacol 181, 3019□3038.

32 Saito A, Kise R & Inoue A (2024) Generation of Comprehensive GPCR-Transducer-Deficient Cell Lines to Dissect the Complexity of GPCR Signaling. Pharmacol Rev 76, 599□619.

33 Dixon AS, Schwinn MK, Hall MP, Zimmerman K, Otto P, Lubben TH, Butler BL, Binkowski BF, Machleidt T, Kirkland TA, Wood MG, Eggers CT, Encell LP & Wood KV (2016) NanoLuc Complementation Reporter Optimized for Accurate Measurement of Protein Interactions in Cells. ACS Chem Biol 11, 400□408.

34 Li HZ, Wang YF, Shao XX, Liu YL, Xu ZG, Wang SL & Guo ZY (2023) FAM237A, rather than peptide PEN and proCCK56-63, binds to and activates the orphan receptor GPR83. FEBS J 290, 3461□3479.

35 Li HZ, Wang YF, Hu WF, Liu YL, Xu ZG & Guo ZY. (2023) Nanomolar range of FAM237B can activate receptor GPR83. Amino Acids 55, 1557□1562.

36 Zheng YS, Liu YL, Xu ZG, He C, Guo ZY. (2024) Is myeloid-derived growth factor a ligand of the sphingosine-1-phosphate receptor 2? Biochem Biophys Res Commun 706, 149766.

37 Sallee NA, Lee E, Leffert A, Ramirez S, Brace AD, Halenbeck R, Kavanaugh WM & Sullivan KMC (2020) A Pilot Screen of a Novel Peptide Hormone Library Identified Candidate GPR83 Ligands. SLAS Discov 25, 1047□1063.

38. Al Hamwi G, Riedel YK, Clemens S, Namasivayam V, Thimm D & Müller CE (2022) MAS-related G protein-coupled receptors X (MRGPRX): Orphan GPCRs with potential as targets for future drugs. Pharmacol Ther 238, 108259.

39 Laschet C, Dupuis N & Hanson J (2018) The G protein-coupled receptors deorphanization landscape. Biochem Pharmacol 153, 62−74.

40 Davenport AP, Alexander SP, Sharman JL, Pawson AJ, Benson HE, Monaghan AE, Liew WC, Mpamhanga CP, Bonner TI, Neubig RR, Pin JP, Spedding M & Harmar AJ (2013) International Union of Basic and Clinical Pharmacology. LXXXVIII. G protein-coupled receptor list: recommendations for new pairings with cognate ligands. Pharmacol Rev 65, 967−986.

