## Supplementary Tables and Figures for "Identification of the orphan GPR25 as a receptor of the chemokine CXCL17"

### Contents:

**Table S1.** Accession numbers of CXCL17 orthologs aligned in Fig. S1.

**Table S2.** Accession numbers of GPR25 orthologs aligned in Fig. S2.

**Table S3.** Possible binding of human CXCL17 with human GPCRs predicted by AlphaFold 3.

**Table S4.** AlphaFold 3-predicted possible binding of CXCL17–GPR25 pairs from different species.

**Table S5.** Primers and vectors used for generation of human GPR25 expression constructs.

**Fig. S1.** Amino acid sequence alignment of CXCL17 orthologs.

**Fig. S2.** Amino acid sequence alignment of GPR25 orthologs.

**Fig. S3.** The nucleotide and amino acid sequences of the designed CXCL17 precursor.

**Fig. S4.** The nucleotide and amino acid sequences of human GPR25 constructs.

**Table S1.** Accession numbers of CXCL17 orthologs aligned in Fig. S1. These orthologs were manually downloaded from NCBI (https://ncbi.nlm.nih.gov/gene) and aligned via AlignX algorithm using the Vector NTI 11.5.1 software.

| Species | Gene ID | mRNA ID | Protein ID |
|---|---|---|---|
| *Homo sapiens* | 284340 | NM_198477 | NP_940879 |
| *Pan troglodytes* | 741429 | XM_001154726 | XP_001154726 |
| *Pan paniscus* | 100975966 | XM_003811758 | XP_003811806 |
| *Macaca mulatta* | 708108 | XM_001105835 | XP_001105835 |
| *Sapajus apella* | 116530866 | XM_032249326 | XP_032105217 |
| *Suricata suricatta* | 115281288 | XM_029926676 | XP_029782536 |
| *Galeopterus variegatus* | 103586108 | XM_008567252 | XP_008565474 |
| *Mus musculus* | 232983 | NM_153576 | NP_705804 |
| *Rattus norvegicus* | 308436 | NM_001107491 | NP_001100961 |
| *Mesocricetus auratus* | 101826068 | XM_021223580 | XP_021079239 |
| *Octodon degus* | 101573419 | XM_023704517 | XP_023560285 |
| *Arvicola amphibius* | 119821858 | XM_038340966 | XP_038196894 |
| *Arvicanthis niloticus* | 117720566 | XM_034519100 | XP_034374991 |
| *Mastomys coucha* | 116100415 | XM_031384229 | XP_031240089 |
| *Peromyscus leucopus* | 114685351 | XM_028859975 | XP_028715808 |
| *Grammomys surdaster* | 114632694 | XM_028781282 | XP_028637115 |
| *Marmota marmota* | 107151419 | XM_015496800 | XP_015352286 |
| *Fukomys damarensis* | 104861740 | XM_010623332 | XP_010621634 |
| *Chinchilla lanigera* | 102020166 | XM_005412354 | XP_005412411 |
| *Microtus ochrogaster* | 101982135 | XM_005361160 | XP_005361217 |
| *Heterocephalus glaber* | 101708913 | XM_004873008 | XP_004873065 |
| *Cavia porcellus* | 100734695 | XM_013146847 | XP_013002301 |
| *Felis catus* | 101086020 | XM_003997765 | XP_003997814 |
| *Canis lupus familiaris* | 111090090 | XM_038656810 | XP_038512738 |
| *Mustela putorius furo* | 101683082 | XM_004780427 | XP_004780484 |
| *Leopardus geoffroyi* | 123578900 | XM_045442260 | XP_045298216 |
| *Prionailurus bengalensis* | 122493991 | XM_043598820 | XP_043454755 |
| *Panthera leo* | 122207597 | XM_042917645 | XP_042773579 |
| *Puma yagouaroundi* | 121018500 | XM_040457200 | XP_040313134 |
| *Vulpes lagopus* | 121484348 | XM_041743498 | XP_041599432 |
| *Hyaena hyaena* | 120240684 | XM_039245502 | XP_039101433 |
| *Lontra canadensis* | 116857606 | XM_032841857 | XP_032697748 |
| *Mustela erminea* | 116580309 | XM_032326546 | XP_032182437 |
| *Ursus arctos* | 113243523 | XM_026482222 | XP_026338007 |
| *Vulpes vulpes* | 112931475 | XM_026014200 | XP_025869985 |
| *Puma concolor* | 112851386 | XM_025914921 | XP_025770706 |
| *Pteropus vampyrus* | 105307357 | XM_011382839 | XP_011381141 |
| *Panthera tigris* | 102961962 | XM_007086589 | XP_007086651 |
| *Pteropus alecto* | 102889606 | XM_006903850 | XP_006903912 |
| *Lutra lutra* | 125088637 | XM_047709872 | XP_047565828 |
| *Meles meles* | 123931218 | XM_045988111 | XP_045844067 |
| *Neogale vison* | 122912075 | XM_044257394 | XP_044113329 |

| | | | |
|---|---|---|---|
| *Talpa occidentalis* | 119249391 | XM_037516694 | XP_037372591 |
| *Sturnira hondurensis* | 119000497 | XM_037065531 | XP_036921426 |
| *Loxodonta africana* | 100668516 | XM_003406663 | XP_003406711 |
| *Bos taurus* | 788717 | NM_001083799 | NP_001077268 |
| *Camelus ferus* | 102511827 | XM_032485980 | XP_032341871 |
| *Odocoileus virginianus* | 110123322 | XM_020871140 | XP_020726799 |
| *Bison bison bison* | 104997641 | XM_010852534 | XP_010850836 |
| *Bubalus bubalis* | 102401266 | XM_006052397 | XP_006052459 |
| *Capra hircus* | 102181231 | XM_018062511 | XP_017918000 |
| *Ochotona princeps* | 101528914 | XM_012930594 | XP_012786048 |
| *Balaenoptera musculus* | 118885645 | XM_036834491 | XP_036690386 |
| *Halichoerus grypus* | 118542831 | XM_036103355 | XP_035959248 |
| *Mirounga leonina* | 117998055 | XM_034986582 | XP_034842473 |
| *Tursiops truncatus* | 117309080 | XM_033845069 | XP_033700960 |
| *Phocoena sinus* | 116744452 | XM_032614237 | XP_032470128 |
| *Phoca vitulina* | 116622190 | XM_032388455 | XP_032244346 |
| *Monodon monoceros* | 114903747 | XM_029236980 | XP_029092813 |
| *Eumetopias jubatus* | 114197075 | XM_028087908 | XP_027943709 |
| *Zalophus californianus* | 113935480 | XM_027617446 | XP_027473247 |
| *Lagenorhynchus obliquidens* | 113605953 | XM_027079714 | XP_026935515 |
| *Callorhinus ursinus* | 112807009 | XM_025849582 | XP_025705367 |
| *Physeter catodon* | 102983204 | XM_024132473 | XP_023988241 |
| *Trichechus manatus latirostris* | 101348725 | XM_004388765 | XP_004388822 |
| *Orcinus orca* | 101278327 | XM_004271217 | XP_004271265 |
| *Equus caballus* | 100629292 | XM_003362327 | XP_003362375 |
| *Choloepus didactylus* | 119521473 | XM_037819798 | XP_037675726 |
| *Tupaia chinensis* | 102492896 | XM_014582832 | XP_014438318 |
| *Erinaceus europaeus* | 103122352 | XM_007532948 | XP_007533010 |
| *Manis pentadactyla* | 118918052 | XM_036896470 | XP_036752365 |
| *Myotis myotis* | 118674385 | XM_036346119 | XP_036202012 |
| *Desmodus rotundus* | 112320178 | XM_024577629 | XP_024433397 |
| *Pipistrellus kuhlii* | 118702335 | XM_036408200 | XP_036264093 |
| *Molossus molossus* | 118639378 | XM_036275769 | XP_036131662 |
| *Rhinolophus ferrumequinum* | 117035523 | XM_033129434 | XP_032985325 |
| *Phyllostomus discolor* | 114511635 | XM_028530396 | XP_028386197 |
| *Miniopterus natalensis* | 107534226 | XM_016209250 | XP_016064736 |
| *Rousettus aegyptiacus* | 107502574 | XM_016129904 | XP_015985390 |
| *Phascolarctos cinereus* | 110219727 | XM_021003326 | XP_020858985 |
| *Trichosurus vulpecula* | 118835868 | XM_036743155 | XP_036599050 |
| *Sarcophilus harrisii* | 116422895 | XM_031963247 | XP_031819107 |
| *Dromiciops gliroides* | 122745891 | XM_043991340 | XP_043847275 |
| *Vombatus ursinus* | 114040160 | XM_027858170 | XP_027713971 |
| *Ornithorhynchus anatinus* | 103166888 | XM_029066640 | XP_028922473 |
| *Tachyglossus aculeatus* | 119946135 | XM_038767563 | XP_038623491 |

**Table S2.** Accession numbers of GPR25 orthologs aligned in Fig. S2. These orthologs were manually downloaded from NCBI database (https://ncbi.nlm.nih.gov/gene/) and aligned via AlignX algorithm using the Vector NTI 11.5.1 software.

| Class | Species | Gene ID | mRNA ID | Protein ID |
|---|---|---|---|---|
| Mammals | *Homo sapiens* | 2848 | NM_005298 | NP_005289 |
| | *Pan troglodytes* | 469633 | XM_016935344 | XP_016790833 |
| | *Macaca mulatta* | 709001 | XM_001109302 | XP_001109302 |
| | *Mus musculus* | 383563 | NM_001101516 | NP_001094986 |
| | *Rattus norvegicus* | 363993 | NM_001398594 | NP_001385523 |
| | *Urocitellus parryii* | 113188759 | XM_026397226 | XP_026253011 |
| | *Mesocricetus auratus* | 106021411 | XM_013115850 | XP_012971304 |
| | *Marmota monax* | 124107106 | XM_046465023 | XP_046320979 |
| | *Cavia porcellus* | 106028732 | XM_063239898 | XP_063095968 |
| | *Equus caballus* | 111771301 | XM_023632516 | XP_023488284 |
| | *Bos taurus* | 107133259 | XM_015475319 | XP_015330805 |
| | *Ovis aries* | 114117307 | XM_027976131 | XP_027831932 |
| | *Sus scrofa* | 110255658 | XM_021064502 | XP_020920161 |
| | *Orycteropus afer afer* | 103197286 | XM_007941182 | XP_007939373 |
| | *Felis catus* | 101098111 | XM_023247599 | XP_023103367 |
| | *Panthera tigris* | 122235733 | XM_042975503 | XP_042831437 |
| | *Panthera leo* | 122211437 | XM_042924579 | XP_042780513 |
| | *Lutra lutra* | 125086389 | XM_047705680 | XP_047561636 |
| | *Meles meles* | 123928346 | XM_045983463 | XP_045839419 |
| | *Neogale vison* | 122918176 | XM_044266324 | XP_044122259 |
| | *Choloepus didactylus* | 119516084 | XM_037812268 | XP_037668196 |
| | *Lontra canadensis* | 116879773 | XM_032878221 | XP_032734112 |
| | *Sapajus apella* | 116558669 | XM_032288671 | XP_032144562 |
| | *Lagenorhynchus obliquidens* | 113614478 | XM_027096014 | XP_026951815 |
| | *Physeter catodon* | 112066689 | XM_024130611 | XP_023986379 |
| | *Balaenoptera musculus* | 118879986 | XM_036823319 | XP_036679214 |
| | *Monodon monoceros* | 114898263 | XM_029227463 | XP_029083296 |
| | *Loxodonta africana* | 100672521 | XM_064273428 | XP_064129498 |
| | *Oryctolagus cuniculus* | 100357510 | XM_051821096 | XP_051677056 |
| | *Manis javanica* | 118969821 | XM_037004672 | XP_036860567 |
| | *Dasypus novemcinctus* | 101428608 | XM_004478694 | XP_004478751 |
| | *Hyaena hyaena* | 120228895 | XM_039227546 | XP_039083477 |
| | *Erinaceus europaeus* | 132534721 | XM_060179177 | XP_060035160 |
| | *Mustela lutreola* | 131815237 | XM_059146936 | XP_059002919 |
| | *Nycticebus coucang* | 128596150 | XM_053605626 | XP_053461601 |
| | *Myotis daubentonii* | 132222403 | XM_059677331 | XP_059533314 |
| | *Phyllostomus hastatus* | 123805005 | XM_045817451 | XP_045673407 |
| | *Gracilinanus agilis* | 123245948 | XM_044674916 | XP_044530851 |
| | *Trichosurus vulpecula* | 118845788 | XM_036753817 | XP_036609712 |
| | *Vombatus ursinus* | 114029117 | XM_027843464 | XP_027699265 |
| | *Dromiciops gliroides* | 122754650 | XM_044003143 | XP_043859078 |
| | *Phascolarctos cinereus* | 110219523 | XM_021002910 | XP_020858569 |

| | | | | |
|---|---|---|---|---|
| | *Sarcophilus harrisii* | 100926064 | XM_003770662 | XP_003770710 |
| | *Monodelphis domestica* | 100027263 | XM_007507853 | XP_007507915 |
| | *Tachyglossus aculeatus* | 119930418 | XM_038748745 | XP_038604673 |
| | *Ornithorhynchus anatinus* | 100093467 | XM_029069673 | XP_028925506 |
| Birds | *Gallus gallus* | 770582 | XM_015275387 | XP_015130873 |
| | *Tyto alba* | 116961340 | XM_032992033 | XP_032847924 |
| | *Cygnus olor* | 121059087 | XM_040535442 | XP_040391376 |
| | *Aythya fuligula* | 116498693 | XM_032203033 | XP_032058924 |
| | *Dromaius novaehollandiae* | 135323481 | XM_064498340 | XP_064354410 |
| | *Harpia harpyja* | 128154745 | XM_052815786 | XP_052671746 |
| | *Chaetura pelagica* | 104391103 | XM_010000329 | XP_009998631 |
| | *Dryobates pubescens* | 104298804 | XM_009898901 | XP_009897203 |
| | *Falco biarmicus* | 130159778 | XM_056361775 | XP_056217750 |
| | *Cuculus canorus* | 104062316 | XM_054087959 | XP_053943934 |
| Reptiles | *Chelonia mydas* | 102942126 | XM_027834774 | XP_027690575 |
| | *Gavialis gangeticus* | 109295412 | XM_019514272 | XP_019369817 |
| | *Alligator sinensis* | 102371715 | XM_006038971 | XP_006039033 |
| | *Anolis carolinensis* | 100553771 | XM_062978417 | XP_062834487 |
| | *Python bivittatus* | 112541279 | XM_025170036 | XP_025025804 |
| | *Emys orbicularis* | 135876995 | XM_065402312 | XP_065258384 |
| | *Candoia aspera* | 134494859 | XM_063300105 | XP_063156175 |
| | *Rhineura floridana* | 133386853 | XM_061630622 | XP_061486606 |
| | *Chelonoidis abingdonii* | 116832686 | XM_032793590 | XP_032649481 |
| | *Podarcis muralis* | 114599189 | XM_028733912 | XP_028589745 |
| | *Varanus komodoensis* | 123034042 | XM_044451090 | XP_044307025 |
| | *Gekko japonicus* | 107110959 | XM_015411818 | XP_015267304 |
| | *Terrapene triunguis* | 112121048 | XM_024220160 | XP_024075928 |
| | *Zootoca vivipara* | 118088240 | XM_035121611 | XP_034977502 |
| Amphibians | *Microcaecilia unicolor* | 115482117 | XM_030221686 | XP_030077546 |
| | *Rhinatrema bivittatum* | 115074284 | XM_029573602 | XP_029429462 |
| | *Bombina bombina* | 128652406 | XM_053705343 | XP_053561318 |
| | *Geotrypetes seraphini* | 117347567 | XM_033918662 | XP_033774553 |
| | *Nanorana parkeri* | 108799369 | XM_018571232 | XP_018426734 |
| | *Xenopus tropicalis* | 100486541 | XM_002941165 | XP_002941211 |
| Fishes | *Danio rerio* | 795188 | XM_021479994 | XP_021335669 |
| | *Astyanax mexicanus* | 103045076 | XM_007234744 | XP_007234806 |
| | *Amphiprion ocellaris* | 111579763 | XM_023287176 | XP_023142944 |
| | *Carcharodon carcharias* | 121282297 | XM_041195951 | XP_041051885 |
| | *Leucoraja erinacea* | 129708680 | XM_055654601 | XP_055510576 |
| | *Xiphias gladius* | 120783363 | XM_040116319 | XP_039972253 |
| | *Callorhinchus milii* | 103179173 | XM_007894305 | XP_007892496 |
| | *Latimeria chalumnae* | 102365624 | XM_005988473 | XP_005988535 |

**Table S3.** Possible binding of human CXCL17 with human GPCRs predicted by AlphaFold 3.

| Name | Gene ID | mRNA ID | Protein ID | Amino acid sequence of mature protein (without signal peptide) | ipTM | pTM | Note |
|---|---|---|---|---|---|---|---|
| CXCL17 | 284340 | NM_198477 | NP_940879 | SSLNPGVARGHRDRGQASRRWLQEGGQECECKDWFLRAPRRKFMTVSGLPKKQCPCDHFKGNVKKTRHQRHHRKPNKHSRACQQFLKQCQLRSFALPL | | | |
| ACKR1a | 2532 | NM_001122951 | NP_001116423 | MASSGYVLQAELSPSTENSSQLDFEDVWNSSYGVNDSFPDGDYGANLEAAAPCHSCNLLDDSALPFFILTSVLGILASSTVLFMLFRPLFRWQLCPGWPVLAQLAVGSALFSIVVPVLAPGLGSTRSSALCSLGYCVWYGSAFAQALLLGCHASLGHRLGAGQVPGLTLGLTVGIWGVAALLTLPVTLASGASGGLCTLIYSTELKALQATHTVACLAIFVLLPLGLFGAKGLKKALGMGPGPWMNILWAWFIFWWPHGVVLGLDFLVRSKLLLLSTCLAQQALDLLLNLAEALAILHCVATPLLLALFCHQATRTLLPSLPLPEGWSSHLDTLGSKS | 0.27 | 0.62 | |
| ACKR1b | 2532 | NM_002036 | NP_002027 | MGNCLHRAELSPSTENSSQLDFEDVWNSSYGVNDSFPDGDYGANLEAAAPCHSCNLLDDSALPFFILTSVLGILASSTVLFMLFRPLFRWQLCPGWPVLAQLAVGSALFSIVVPVLAPGLGSTRSSALCSLGYCVWYGSAFAQALLLGCHASLGHRLGAGQVPGLTLGLTVGIWGVAALLTLPVTLASGASGGLCTLIYSTELKALQATHTVACLAIFVLLPLGLFGAKGLKKALGMGPGPWMNILWAWFIFWWPHGVVLGLDFLVRSKLLLLSTCLAQQALDLLLNLAEALAILHCVATPLLLALFCHQATRTLLPSLPLPEGWSSHLDTLGSKS | 0.24 | 0.62 | |
| ACKR2 | 1238 | NM_001296 | NP_001287 | MAATASPQPLATEDADSENSSFYYYDYLDEVAFMLCRKDAVVSFGKVFLPVFYSLIFVLGLSGNLLLLMVLLRYVPRRRMVEIYLLNLAISNLLFLVTLPFWGISVAWHWVFGSFLCKMVSTLYTINFYSGIFFISCMSLDKYLEIVHAQPYHRLRTRAKSLLLATIVWAVSLAVSIPDMVFVQTHENPKGVWNCHADFGGHGTIWKLFLRFQQNLLGFLLPLLAMIFFYSRIGCVLVRLRPAGQGRALKIAAALVVAFFVLWFPYNLTLFLHTLLDLQVFGNCEVSQHLDYALQVTESIAFLHCCFSPILYAFSSHRFRQYLKAFLAAVLGWHLAPGTAQASLSSCSESSILTAQEEMTGMNDLGERQSENYPNKEDVGNKSA | 0.3 | 0.65 | |
| ACKR3 | 57007 | NM_020311 | NP_064707 | MDLHLFDYSEPGNFSDISWPCNSSDCIVVDTVMCPNMPNKSVLLYTLSFIYIFIFVIGMIANSVVVWVNIQAKTTGYDTHCYILNLAIADLWVVLTIPVWVVSLVQHNQWPMGELTCKVTHLIFSINLFGSIFFLTCMSVDRYLSITYFTNTPSSRKKMVRRVVCILVWLLAFCVSLPDTYYLKTVTSASNNETYCRSFYPEHSIKEWLIGMELVSVVLGFAVPFSIIAVFYFLLARAISASSDQEKHSSRKIIFSYVVVFLVCWLPYHVAVLLDIFSILHYIPFTCRLEHALFTALHVTQCLSLVHCCVNPVLYSFINRNYRYELMKAFIFKYSAKTGLTKLIDASRVSETEYSALEQSTK | 0.26 | 0.66 | |
| ACKR4 | 51554 | NM_016557 | NP_057641 | MALEQNQSTDYYYEENEMNGTYDYSQYELICIKEDVREFAKVFLPVFLTIVFVIGLAGNSMVVAIYAYYKKQRTKTDVYILNLAVADLLLLFTLPFWAVNAVHGWVLGKIMCKITSALYTLNFVSGMQFLACISIDRYVAVTKVPSQSGVGKPCWIICFCVWMAAILLSIPQLVFYTVNDNARCIPIFPRYLGTSMKALIQMLEICIGFVVPFLIMGVCYFITARTLMKMPNIKISRPLKVLLTVVIVFIVTQLPYNIVKFCRAIDIIYSLITSCNMSKRMDIAIQVTESIALFHSCLNPILYVFMGASFKNYVMKVAKKYGSWRRQRQSVEEFPFDSEGPTEPTSTFSI | 0.18 | 0.62 | |
| CCR1 | 1230 | NM_001295 | NP_001286 | METPNTTEDYDTTTEFDYGDATPCQKVNERAFGAQLLPPLYSLVFVIGLVGNILVVLVLVQYKRLKNMTSIYLLNLAISDLLFLFTLPFWIDYKLKDDWVFGDAMCKILSGFYYTGLYSEIFFIILLTIDRYLAIVHAVFALRARTVTFGVITSIIIWALAILASMPGLYFSKTQWEFTHHTCSLHFPHESLREWKLFQALKLNLFGLVLPLLVMIICYTGIIKILLRRPNEKKSKAVRLIFVIMIIFFLFWTPYNLTILISVFQDFLFTHECEQSRHLDLAVQVTEVIAYTHCCVNPVIYAFVGERFRKYLRQLFHRRVAVHLVKWLPFLSVDRLERVSSTSPSTGEHELSAGF | 0.16 | 0.66 | |
| CCR2a | 729230 | NM_001123041 | NP_001116513 | MLSTSRSRFIRNTNESGEEVTTFFDYDYGAPCHKFDVKQIGAQLLPPLYSLVFIFGFVGNMLVVLILINCKKLKCLTDIYLLNLAISDLLFLITLPLWAHSAANEWVFGNAMCKLFTGLYHIGYFGGIFFIILLTIDRYLAIVHAVFALKARTVTFGVVTSVITWLVAVFASVPGIIFTKCQKEDSVYVCGPYFPRGWNNFHTIMRNILGLVLPLLIMVICYSGILKTLLRCRNEKKRHRAVRVIFTIMIVYFLFWTPYNIVILLNTFQEFFGLSNCESTSQLDQATQVTETLGMTHCCINPIIYAFVGEKFRSLFHIALGCRIAPLQKPVCGGPGVRPGKNVKVTTQGLLDGRGKGKSIGRAPEASLQDKEGA | 0.33 | 0.64 | |
| CCR2b | 729230 | NM_001123396 | NP_001116868 | MLSTSRSRFIRNTNESGEEVTTFFDYDYGAPCHKFDVKQIGAQLLPPLYSLVFIFGFVGNMLVVLILINCKKLKCLTDIYLLNLAISDLLFLITLPLWAHSAANEWVFGNAMCKLFTGLYHIGYFGGIFFIILLTIDRYLAIVHAVFALKARTVTFGVVTSVITWLVAVFASVPGIIFTKCQKEDSVYVCGPYFPRGWNNFHTIMRNILGLVLPLLIMVICYSGILKTLLRCRNEKKRHRAVRVIFTIMIVYFLFWTPYNIVILLNTFQEFFGLSNCESTSQLDQATQVTETLGMTHCCINPIIYAFVGEKFRRYLSVFFRKHITKRFCKQCPVFYRETVDGVTSTNTPSTGEQEVSAGL | 0.22 | 0.65 | |

| | | | | | | |
|---|---|---|---|---|---|---|
| CCR3-1 | 1232 | NM_001837 | NP_001828 | MTTSLDTVETFGTTSYYDDVGLLCEKADTRALMAQFVPPLYSLVFTVGLLGNVVVVMILIKYRRLRIMTNIYLLNLAISDLLFLVTLPFWIHYVRGHNWVFGHGMCKLLSGFYHTGLYSEIFFIILLTIDRYLAIVHAVFALRARTVTFGVITSIVTWGLAVLAALPEFIFYETEELFEETLCSALYPEDTVYSWRHFHTLRMTIFCLVLPLLVMAICYTGIIKTLLRCPSKKKYKAIRLIFVIMAVFFIFWTPYNVAILLSSYQSILFGNDCERSKHLDLVMLVTEVIAYSHCCMNPVIYAFVGERFRKYLRHFFHRHLLMHLGRYIPFLPSEKLERTSSVSPSTAEPELSIVF | 0.2 | 0.67 |
| CCR3-2 | 1232 | NM_178328 | NP_847898 | MPFGIRMLLRAHKPGSSRRSEMTTSLDTVETFGTTSYYDDVGLLCEKADTRALMAQFVPPLYSLVFTVGLLGNVVVVMILIKYRRLRIMTNIYLLNLAISDLLFLVTLPFWIHYVRGHNWVFGHGMCKLLSGFYHTGLYSEIFFIILLTIDRYLAIVHAVFALRARTVTFGVITSIVTWGLAVLAALPEFIFYETEELFEETLCSALYPEDTVYSWRHFHTLRMTIFCLVLPLLVMAICYTGIIKTLLRCPSKKKYKAIRLIFVIMAVFFIFWTPYNVAILLSSYQSILFGNDCERSKHLDLVMLVTEVIAYSHCCMNPVIYAFVGERFRKYLRHFFHRHLLMHLGRYIPFLPSEKLERTSSVSPSTAEPELSIVF | 0.15 | 0.65 |
| CCR3-3 | 1232 | NM_001164680 | NP_001158152 | MPFGIRMLLRAHKPGRSEMTTSLDTVETFGTTSYYDDVGLLCEKADTRALMAQFVPPLYSLVFTVGLLGNVVVVMILIKYRRLRIMTNIYLLNLAISDLLFLVTLPFWIHYVRGHNWVFGHGMCKLLSGFYHTGLYSEIFFIILLTIDRYLAIVHAVFALRARTVTFGVITSIVTWGLAVLAALPEFIFYETEELFEETLCSALYPEDTVYSWRHFHTLRMTIFCLVLPLLVMAICYTGIIKTLLRCPSKKKYKAIRLIFVIMAVFFIFWTPYNVAILLSSYQSILFGNDCERSKHLDLVMLVTEVIAYSHCCMNPVIYAFVGERFRKYLRHFFHRHLLMHLGRYIPFLPSEKLERTSSVSPSTAEPELSIVF | 0.22 | 0.67 |
| CCR4 | 1233 | NM_005508 | NP_005499 | MNPTDIADTTLDESIYSNYYLYESIPKPCTKEGIKAFGELFLPPLYSLVFVFGLLGNSVVVLVLFKYKRLRSMTDVYLLNLAISDLLFVFSLPFWGYYAADQWVFGLGLCKMISWMYLVGFYSGIFFVMLMSIDRYLAIVHAVFSLRARTLTYGVITSLATWSVAVFASLPGFLFSTCYTERNHTYCKTKYSLNSTTWKVLSSLEINILGLVIPLGIMLFCYSMIIRTLQHCKNEKKNKAVKMIFAVVVLFLGFWTPYNIVLFLETLVELEVLQDCTFERYLDYAIQATETLAFVHCCLNPIIYFFLGEKFRKYILQLFKTCRGLFVLCQYCGLLQIYSADTPSSSYTQSTMDHDLHDAL | 0.4 | 0.68 |
| CCR5 | 1234 | NM_000579 | NP_000570 | MDYQVSSPIYDINYYTSEPCQKINVKQIAARLLPPLYSLVFIFGFVGNMLVILILINCKRLKSMTDIYLLNLAISDLFFLLTVPFWAHYAAAQWDFGNTMCQLLTGLYFIGFFSGIFFIILLTIDRYLAVVHAVFALKARTVTFGVVTSVITWVVAVFASLPGIIFTRSQKEGLHYTCSSHFPYSQYQFWKNFQTLKIVILGLVLPLLVMVICYSGILKTLLRCRNEKKRHRAVRLIFTIMIVYFLFWAPYNIVLLLNTFQEFFGLNNCSSSNRLDQAMQVTETLGMTHCCINPIIYAFVGEKFRNYLLVFFQKHIAKRFCKCCSIFQQEAPERASSVYTRSTGEQEISVGL | 0.23 | 0.67 |
| CCR6 | 1235 | NM_001394582 | NP_001381511 | MSGESMNFSDVFDSSEDYFVSVNTSYYSVDSEMLLCSLQEVRQFSRLFVPIAYSLICVFGLLGNILVVITFAFYKKARSMTDVYLLNMAIADILFVLTLPFWAVSHATGAWVFSNATCKLLKGIYAINFNCGMLLLTCISMDRYIAIVQATKSFRLRSRTLPRSKIICLVVWGLSVIISSSTFVFNQKYNTQGSDVCEPKYQTVSEPIRWKLLMLGLELLFGFFIPLMFMIFCYTFIVKTLVQAQNSKRHKAIRVIIAVVLVFLACQIPHNMVLLVTAANLGKMNRSCQSEKLIGYTKTVTEVLAFLHCCLNPVLYAFIGQKFRNYFLKILKDLWCVRRKYKSSGFSCAGRYSENISRQTSETADNDNASSFTM | 0.23 | 0.64 |
| CCR7a | 1236 | NM_001838 | NP_001829 | MDLGKPMKSVLVVALLVIFQVCLCQDEVTDDYIGDNTTVDYTLFESLCSKKDVRNFKAWFLPIMYSIICFVGLLGNGLVVLTYIYFKRLKTMTDTYLLNLAVADILFLLTLPFWAYSAAKSWVFGVHFCKLIFAIYKMSFFSGMLLLLCISIDRYVAIVQAVSAHRHRARVLLISKLSCVGIWILATVLSIPELLYSDLQRSSSEQAMRCSLITEHVEAFITIQVAQMVIGFLVPLLAMSFCYLVIIRTLLQARNFERNKAIKVIIAVVVVFIVFQLPYNGVVLAQTVANFNITSSTCELSKQLNIAYDVTYSLACVRCCVNPFLYAFIGVKFRNDLFKLFKDLGCLSQEQLRQWSSCRHIRRSSMSVEAETTTTFSP | 0.17 | 0.6 |
| CCR7b | 1236 | NM_001301714 | NP_001288643 | MYSIICFVGLLGNGLVVLTYIYFKRLKTMTDTYLLNLAVADILFLLTLPFWAYSAAKSWVFGVHFCKLIFAIYKMSFFSGMLLLLCISIDRYVAIVQAVSAHRHRARVLLISKLSCVGIWILATVLSIPELLYSDLQRSSSEQAMRCSLITEHVEAFITIQVAQMVIGFLVPLLAMSFCYLVIIRTLLQARNFERNKAIKVIIAVVVVFIVFQLPYNGVVLAQTVANFNITSSTCELSKQLNIAYDVTYSLACVRCCVNPFLYAFIGVKFRNDLFKLFKDLGCLSQEQLRQWSSCRHIRRSSMSVEAETTTTFSP | 0.14 | 0.53 |
| CCR7c | 1236 | NM_001301716 | NP_001288645 | MKSVLVVALLVIFQVCLCQDEVTDDYIGDNTTVDYTLFESLCSKKDVRNFKAWFLPIMYSIICFVGLLGNGLVVLTYIYFKRLKTMTDTYLLNLAVADILFLLTLPFWAYSAAKSWVFGVHFCKLIFAIYKMSFFSGMLLLLCISIDRYVAIVQAVSAHRHRARVLLISKLSCVGIWILATVLSIPELLYSDLQRSSSEQAMRCSLITEHVEAFITIQVAQMVIGFLVPLLAMSFCYLVIIRTLLQARNFERNKAIKVIIAVVVVFIVFQLPYNGVVLAQTVANFNITSSTCELSKQLNIAYDVTYSLACVRCCVNPFLYAFIGVKFRNDLFKLFKDLGCLSQEQLRQWSSCRH | 0.13 | 0.6 |

| | | | | | | |
|---|---|---|---|---|---|---|
| | | | | IRRSSMSVEAETTTTFSP | | |
| CCR8 | 1237 | NM_005201 | NP_005192 | MDYTLDLSVTTVTDYYYPDIFSSPCDAELIQTNGKLLLAVFYCLLFVFSLLGNSLVILV<br>LVVCKKLRSITDVYLLNLALSDLLFVFSFPFQTYYLLDQWVFGTVMCKVVSGFYYIGFY<br>SSMFFITLMSVDRYLAVVHAVYALKVRTIRMGTTLCLAVWLTAIMATIPLLVFYQVASE<br>DGVLQCYSFYNQQTLKWKIFTNFKMNILGLLIPFTIFMFCYIKILHQLKRCQNHNKTKA<br>IRLVLIVVIASLLFWVPFNVVLFLTSLHSMHILDGCSISQQLTYATHVTEIISFTHCCV<br>NPVIYAFVGEKFKKHLSEIFQKSCSQIFNYLGRQMPRESCEKSSSCQQHSSRSSSVDYI<br>L | 0.15 | 0.64 |
| CCR9a | 10803 | NM_001386447 | NP_001373376 | MTPTDFTSPIPNMADDYGSESTSSMEDYVNFNFTDFYCEKNNVRQFASHFLPPLYWLVF<br>IVGALGNSLVILVYWYCTRVKTMTDMFLLNLAIADLLFLVTLPFWAIAAADQWKFQTFM<br>CKVVNSMYKMNFYSCVLLIMCISVDRYIAIAQAMRAHTWREKRLLYSKMVCFTIWVLAA<br>ALCIPEILYSQIKEESGIAICTMVYPSDESTKLKSAVLTLKVILGFFLPFVVMACCYTI<br>IIHTLIQAKKSSKHKALKVTITVLTVFVLSQFPYNCILLVQTIDAYAMFISNCAVSTNI<br>DICFQVTQTIAFFHSCLNPVLYVFVGERFRRDLVKTLKNLGCISQAQWVSFTRREGSLK<br>LSSMLLETTSGALSL | 0.29 | 0.65 |
| CCR9b | 10803 | NM_001256369 | NP_001243298 | MADDYGSESTSSMEDYVNFNFTDFYCEKNNVRQFASHFLPPLYWLVFIVGALGNSLVIL<br>VYWYCTRVKTMTDMFLLNLAIADLLFLVTLPFWAIAAADQWKFQTFMCKVVNSMYKMNF<br>YSCVLLIMCISVDRYIAIAQAMRAHTWREKRLLYSKMVCFTIWVLAAALCIPEILYSQI<br>KEESGIAICTMVYPSDESTKLKSAVLTLKVILGFFLPFVVMACCYTIIIHTLIQAKKSS<br>KHKALKVTITVLTVFVLSQFPYNCILLVQTIDAYAMFISNCAVSTNIDICFQVTQTIAF<br>FHSCLNPVLYVFVGERFRRDLVKTLKNLGCISQAQWVSFTRREGSLKLSSMLLETTSGA<br>LSL | 0.25 | 0.63 |
| CCR10 | 2826 | NM_016602 | NP_057686 | MGTEATEQVSWGHYSGDEEDAYSAEPLPELCYKADVQAFSRAFQPSVSLTVAALGLAGN<br>GLVLATHLAARRAARSPTSAHLLQLALADLLLALTLPFAAAGALQGWSLGSATCRTISG<br>LYSASFHAGFLFLACISADRYVAIARALPAGPRPSTPGRAHLVSVIVWLLSLLLALPAL<br>LFSQDGQREGQRRCRLIFPEGLTQTVKGASAVAQVALGFALPLGVMVACYALLGRTLLA<br>ARGPERRRALRVVVALVAAFVVLQLPYSLALLLDTADLLAARERSCPASKRKDVALLVT<br>SGLALARCGLNPVLYAFLGLRFRQDLRRLLRGGSCPSGPQPRRGCPRRPRLSSCSAPTE<br>THSLSWDN | 0.32 | 0.63 |
| CXCR1 | 3577 | NM_000634 | NP_000625 | MSNITDPQMWDFDDLNFTGMPPADEDYSPCMLETETLNKYVVIIAYALVFLLSLLGNSL<br>VMLVILYSRVGRSVTDVYLLNLALADLLFALTLPIWAASKVNGWIFGTFLCKVVSLLKE<br>VNFYSGILLLACISVDRYLAIVHATRTLTQKRHLVKFVCLGCWGLSMNLSLPFFLFRQA<br>YHPNNSSPVCYEVLGNDTAKWRMVLRILPHTFGFIVPLFVMLFCYGFTLRTLFKAHMGQ<br>KHRAMRVIFAVVLIFLLCWLPYNLVLLADTLMRTQVIQESCERRNNIGRALDATEILGF<br>LHSCLNPIIYAFIGQNFRHGFLKILAMHGLVSKEFLARHRVTSYTSSSVNVSSNL | 0.31 | 0.66 |
| CXCR2 | 3579 | NM_001557 | NP_001548 | MEDFNMESDSFEDFWKGEDLSNYSYSSTLPPFLLDAAPCEPESLEINKYFVVIIYALVF<br>LLSLLGNSLVMLVILYSRVGRSVTDVYLLNLALADLLFALTLPIWAASKVNGWIFGTFL<br>CKVVSLLKEVNFYSGILLLACISVDRYLAIVHATRTLTQKRYLVKFICLSIWGLSLLLA<br>LPVLLFRRTVYSSNVSPACYEDMGNNTANWRMLLRILPQSFGFIVPLLIMLFCYGFTLR<br>TLFKAHMGQKHRAMRVIFAVVLIFLLCWLPYNLVLLADTLMRTQVIQETCERRNHIDRA<br>LDATEILGILHSCLNPLIYAFIGQKFRHGLLKILAIHGLISKDSLPKDSRPSFVGSSSG<br>HTSTTL | 0.21 | 0.6 |
| CXCR3a | 2833 | NM_001504 | NP_001495 | MVLEVSDHQVLNDAEVAALLENFSSSYDYGENESDSCCTSPPCPQDFSLNFDRAFLPAL<br>YSLLFLLGLLGNGAVAAVLLSRRTALSSTDTFLLHLAVADTLLVLTLPLWAVDAAVQWV<br>FGSGLCKVAGALFNINFYAGALLLACISFDRYLNIVHATQLYRRGPPARVTLTCLAVWG<br>LCLLFALPDFIFLSAHHDERLNATHCQYNFPQVGRTALRVLQLVAGFLLPLLVMAYCYA<br>HILAVLLVSRGQRRLRAMRLVVVVVVAFALCWTPYHLVVLVDILMDLGALARNCGRESR<br>VDVAKSVTSGLGYMHCCLNPLLYAFVGVKFRERMWMLLLRLGCPNQRGLQRQPSSSRRD<br>SSWSETSEASYSGL | 0.18 | 0.58 |
| CXCR3b | 2833 | NM_001142797 | NP_001136269 | MELRKYGPGRLAGTVIGGAAQSKSQTKSDSITKEFLPGLYTAPSSPFPPSQVSDHQVLN<br>DAEVAALLENFSSSYDYGENESDSCCTSPPCPQDFSLNFDRAFLPALYSLLFLLGLLGN<br>GAVAAVLLSRRTALSSTDTFLLHLAVADTLLVLTLPLWAVDAAVQWVFGSGLCKVAGAL<br>FNINFYAGALLLACISFDRYLNIVHATQLYRRGPPARVTLTCLAVWGLCLLFALPDFIF<br>LSAHHDERLNATHCQYNFPQVGRTALRVLQLVAGFLLPLLVMAYCYAHILAVLLVSRGQ<br>RRLRAMRLVVVVVVAFALCWTPYHLVVLVDILMDLGALARNCGRESRVDVAKSVTSGLG<br>YMHCCLNPLLYAFVGVKFRERMWMLLLRLGCPNQRGLQRQPSSSRRDSSWSETSEASYS<br>GL | 0.17 | 0.55 |
| CXCR4a | 7852 | NM_001008540 | NP_001008540 | MSIPLPLLQIYTSDNYTEEMGSGDYDSMKEPCFREENANFNKIFLPTIYSIIFLTGIVG<br>NGLVILVMGYQKKLRSMTDKYRLHLSVADLLFVITLPFWAVDAVANWYFGNFLCKAVHV<br>IYTVNLYSSVLILAFISLDRYLAIVHATNSQRPRKLLAEKVVYVGVWIPALLLTIPDFI | 0.23<br>021<br>0.2 | 0.65<br>0.64<br>0.64 |

| | | | | | | |
|---|---|---|---|---|---|---|
| | | | | FANVSEADDRYICDRFYPNDLWVVVFQFQHIMVGLILPGIVILSCYCIIISKLSHSKGHQKRKALKTTVILILAFFACWLPYYIGISIDSFILLEIIKQGCEFENTVHKWISITEALAFFHCCLNPILYAFLGAKFKTSAQHALTSVSRGSSLKILSKGKRGGHSSVSTESESSSFHSS | | |
| CXCR4b | 7852 | NM_003467 | NP_003458 | MEGISIYTSDNYTEEMGSGDYDSMKEPCFREENANFNKIFLPTIYSIIFLTGIVGNGLVILVMGYQKKLRSMTDKYRLHLSVADLLFVITLPFWAVDAVANWYFGNFLCKAVHVIYTVNLYSSVLILAFISLDRYLAIVHATNSQRPRKLLAEKVVYVGVWIPALLLTIPDFIFANVSEADDRYICDRFYPNDLWVVVFQFQHIMVGLILPGIVILSCYCIIISKLSHSKGHQKRKALKTTVILILAFFACWLPYYIGISIDSFILLEIIKQGCEFENTVHKWISITEALAFFHCCLNPILYAFLGAKFKTSAQHALTSVSRGSSLKILSKGKRGGHSSVSTESESSSFHSS | 0.23<br>0.21<br>021 | 0.64<br>0.64<br>0.64 |
| CXCR4c | 7852 | NM_001348056 | NP_001334985 | MEGISENAPLPNVPNAPSDKHEDGKRPTHRRSARLGEEVPFVHFLTLPPNIPQAPKGLRFKTAFSLPTTSCLKPRMIYTSDNYTEEMGSGDYDSMKEPCFREENANFNKIFLPTIYSIIFLTGIVGNGLVILVMGYQKKLRSMTDKYRLHLSVADLLFVITLPFWAVDAVANWYFGNFLCKAVHVIYTVNLYSSVLILAFISLDRYLAIVHATNSQRPRKLLAEKVVYVGVWIPALLLTIPDFIFANVSEADDRYICDRFYPNDLWVVVFQFQHIMVGLILPGIVILSCYCIIISKLSHSKGHQKRKALKTTVILILAFFACWLPYYIGISIDSFILLEIIKQGCEFENTVHKWISITEALAFFHCCLNPILYAFLGAKFKTSAQHALTSVSRGSSLKILSKGKRGGHSSVSTESESSSFHSS | 0.25<br>0.21<br>0.21 | 0.6<br>0.58<br>0.57 |
| CXCR4d | 7852 | NM_001348059 | NP_001334988 | MEGISENAPLPNVPNAPSDKHEDGKRPTHRRSARLGEEIYTSDNYTEEMGSGDYDSMKEPCFREENANFNKIFLPTIYSIIFLTGIVGNGLVILVMGYQKKLRSMTDKYRLHLSVADLLFVITLPFWAVDAVANWYFGNFLCKAVHVIYTVNLYSSVLILAFISLDRYLAIVHATNSQRPRKLLAEKVVYVGVWIPALLLTIPDFIFANVSEADDRYICDRFYPNDLWVVVFQFQHIMVGLILPGIVILSCYCIIISKLSHSKGHQKRKALKTTVILILAFFACWLPYYIGISIDSFILLEIIKQGCEFENTVHKWISITEALAFFHCCLNPILYAFLGAKFKTSAQHALTSVSRGSSLKILSKGKRGGHSSVSTESESSSFHSS | 0.2<br>0.22<br>0.23 | 0.6<br>0.61<br>0.6 |
| CXCR4e | 7852 | NM_001348060 | NP_001334989 | MGSGDYDSMKEPCFREENANFNKIFLPTIYSIIFLTGIVGNGLVILVMGYQKKLRSMTDKYRLHLSVADLLFVITLPFWAVDAVANWYFGNFLCKAVHVIYTVNLYSSVLILAFISLDRYLAIVHATNSQRPRKLLAEKVVYVGVWIPALLLTIPDFIFANVSEADDRYICDRFYPNDLWVVVFQFQHIMVGLILPGIVILSCYCIIISKLSHSKGHQKRKALKTTVILILAFFACWLPYYIGISIDSFILLEIIKQGCEFENTVHKWISITEALAFFHCCLNPILYAFLGAKFKTSAQHALTSVSRGSSLKILSKGKRGGHSSVSTESESSSFHSS | 0.16<br>0.21<br>0.22 | 0.61<br>0.66<br>0.66 |
| CXCR5-1 | 643 | NM_001716 | NP_001707 | MNYPLTLEMDLENLEDLFWELDRLDNYNDTSLVENHLCPATEGPLMASFKAVFVPVAYSLIFLLGVIGNVLVLVILERHRQTRSSTETFLFHLAVADLLLVFILPFAVAEGSVGWVLGTFLCKTVIALHKVNFYCSSLLLACIAVDRYLAIVHAVHAYRHRRLLSIHITCGTIWLVGFLLALPEILFAKVSQGHHNNSLPRCTFSQENQAETHAWFTSRFLYHVAGFLLPMLVMGWCYVGVVHRLRQAQRRPQRQKAVRVAILVTSIFFLCWSPYHIVIFLDTLARLKAVDNTCKLNGSLPVAITMCEFLGLAHCCLNPMLYTFAGVKFRSDLSRLLTKLGCTGPASLCQLFPSWRRSSLSESENATSLTTF | 0.19 | 0.62 |
| CXCR5-2 | 643 | NM_032966 | NP_116743 | MASFKAVFVPVAYSLIFLLGVIGNVLVLVILERHRQTRSSTETFLFHLAVADLLLVFILPFAVAEGSVGWVLGTFLCKTVIALHKVNFYCSSLLLACIAVDRYLAIVHAVHAYRHRRLLSIHITCGTIWLVGFLLALPEILFAKVSQGHHNNSLPRCTFSQENQAETHAWFTSRFLYHVAGFLLPMLVMGWCYVGVVHRLRQAQRRPQRQKAVRVAILVTSIFFLCWSPYHIVIFLDTLARLKAVDNTCKLNGSLPVAITMCEFLGLAHCCLNPMLYTFAGVKFRSDLSRLLTKLGCTGPASLCQLFPSWRRSSLSESENATSLTTF | 0.14 | 0.62 |
| CXCR6 | 10663 | NM_001386435 | NP_001373364 | MAEHDYHEDYGFSSFNDSSQEEHQDFLQFSKVFLPCMYLVVFVCGLVGNSLVLVISIFYHKLQSLTDVFLVNLPLADLVFVCTLPFWAYAGIHEWVFGQVMCKSLLGIYTINFYTSMLILTCITVDRFIVVVKATKAYNQQAKRMTWGKVTSLLIWVISLLVSLPQIIYGNVFNLDKLICGYHDEAISTVVLATQMTLGFFLPLLTMIVCYSVIIKTLLHAGGFQKHRSLKIIFLVMAVFLLTQMPFNLMKFIRSTHWEYYAMTSFHYTIMVTEAIAYLRACLNPVLYAFVSLKFRKNFWKLVKDIGCLPYLGVSHQWKSSEDNSKTFSASHNVEATSMFQL | 0.39 | 0.67 |
| CX3CR1a | 1524 | NM_001171174 | NP_001164645 | MREPLEAFKLADLDFRKSSLASGWRMASGAFTMDQFPESVTENFEYDDLAEACYIGDIVVFGTVFLSIFYSVIFAIGLVGNLLVVFALTNSKKPKSVTDIYLLNLALSDLLFVATLPFWTHYLINEKGLHNAMCKFTTAFFFIGFFGSIFFITVISIDRYLAIVLAANSMNNRTVQHGVTISLGVWAAAILVAAPQFMFTKQKENECLGDYPEVLQEIWPVLRNVETNFLGFLLPLLIMSYCYFRIIQTLFSCKNHKKAKAIKLILLVVIVFFLFWTPYNVMIFLETLKLYDFFPSCDMRKDLRLALSVTETVAFSHCCLNPLIYAFAGEKFRRYLYHLYGKCLAVLCGRSVHVDFSSSESQRSRHGSVLSSNFTYHTSDGDALLLL | 0.16 | 0.61 |
| CX3CR1b | 1524 | NM_001337 | NP_001328 | MDQFPESVTENFEYDDLAEACYIGDIVVFGTVFLSIFYSVIFAIGLVGNLLVVFALTNSKKPKSVTDIYLLNLALSDLLFVATLPFWTHYLINEKGLHNAMCKFTTAFFFIGFFGSIFFITVISIDRYLAIVLAANSMNNRTVQHGVTISLGVWAAAILVAAPQFMFTKQKENECLGDYPEVLQEIWPVLRNVETNFLGFLLPLLIMSYCYFRIIQTLFSCKNHKKAKAIKLILLVVIVFFLFWTPYNVMIFLETLKLYDFFPSCDMRKDLRLALSVTETVAFSHCCLNPLIYAFAGEKFRRYLYHLYGKCLAVLCGRSVHVDFSSSESQRSRHGSVLSSNFTYHTSDGDALLLL | 0.17 | 0.63 |
| XCR1 | 2829 | NM_001024644 | NP_001019815 | MESSGNPESTTFFYYDLQSQPCENQAWVFATLATTVLYCLVFLLSLVGNSLVLWVLVKYESLESLTNIFILNLCLSDLVFACLLPVWISPYHWGWVLGDFLCKLLNMIFSISLYSSIFFLTIMTIHRYLSVVSPLSTLRVPTLRCRVLVTMAVWVASILSSILDTIFHKVLSSGCDYSELTWYLTSVYQHNLFFLLSLGIILFCYVEILRTLFRSRSKRRHRTVKLIFAIVVAYFL | 0.34 | 0.69 |

| | | | | | | | |
|---|---|---|---|---|---|---|---|
| | | | | SWGPYNFTLFLQTLFRTQIIRSCEAKQQLEYALLICRNLAFSHCCFNPVLYVFVGVKFRTHLKHVLRQFWFCRLQAPSPASIPHSPGAFAYEGASFY | | | |
| CCRL2-1 | 9034 | NM_003965 | NP_003956 | MANYTLAPEDEYDVLIEGELESDEAEQCDKYDAQALSAQLVPSLCSAVFVIGVLDNLLVVLILVKYKGLKRVENIYLLNLAVSNLCFLLTLPFWAHAGGDPMCKILIGLYFVGLYSETFFNCLLTVQRYLVFLHKGNFFSARRRVPCGIITSVLAWVTAILATLPEFVVYKPQMEDQKYKCAFSRTPFLPADETFWKHFLTLKMNISVLVLPLFIFTFLYVQMRKTLRFREQRYSLFKLVFAIMVVFLLMWAPYNIAFFLSTFKEHFSLSDCKSSYNLDKSVHITKLIATTHCCINPLLYAFLDGTFSKYLCRCFHLRSNTPLQPRGQSAQGTSREEPDHSTEV | 0.1 | 0.61 | |
| CCRL2-2 | 9034 | NM_001130910 | NP_001124382 | MIYTRFLKGSLKMANYTLAPEDEYDVLIEGELESDEAEQCDKYDAQALSAQLVPSLCSAVFVIGVLDNLLVVLILVKYKGLKRVENIYLLNLAVSNLCFLLTLPFWAHAGGDPMCKILIGLYFVGLYSETFFNCLLTVQRYLVFLHKGNFFSARRRVPCGIITSVLAWVTAILATLPEFVVYKPQMEDQKYKCAFSRTPFLPADETFWKHFLTLKMNISVLVLPLFIFTFLYVQMRKTLRFREQRYSLFKLVFAIMVVFLLMWAPYNIAFFLSTFKEHFSLSDCKSSYNLDKSVHITKLIATTHCCINPLLYAFLDGTFSKYLCRCFHLRSNTPLQPRGQSAQGTSREEPDHSTEV | 0.1 | 0.6 | |
| GPR3 | 2827 | NM_005281 | NP_005272 | MMWGAGSPLAWLSAGSGNVNVSSVGPAEGPTGPAAPLPSPKAWDVVLCISGTLVSCENALVVAIIVGTPAFRAPMFLLVGSLAVADLLAGLGLVLHFAAVFCIGSAEMSLVLVGVLAMAFTASIGSLLAITVDRYLSLYNALTYYSETTVTRTYVMLALVWGGALGLGLLPVLAWNCLDGLTTCGVVYPLSKNHLVVLAIAFFMVFGIMLQLYAQICRIVCRHAQQIALQRHLLPASHYVATRKGIATLAVVLGAFAACWLPFTVYCLLGDAHSPPLYTYLTLLPATYNSMINPIIYAFRNQDVQKVLWAVCCCCSSSKIPFRSRSPSDV | 0.36 | 0.7 | |
| GPR4 | 2828 | NM_005282 | NP_005273 | MGNHTWEGCHVDSRVDHLFPPSLYIFVIGVGLPTNCLALWAAYRQVQQRNELGVYLMNLSIADLLYICTLPLWVDYFLHHDNWIHGPGSCKLFGFIFYTNIYISIAFLCCISVDRYLAVAHPLRFARLRRVKTAVAVSSVVWATELGANSAPLFHDELFRDRYNHTFCFEKFPMEGWVAWMNLYRVFVGFLFPWALMLLSYRGILRAVRGSVSTERQEKAKIKRLALSLIAIVLVCFAPYHVLLLSRSAIYLGRPWDCGFEERVFSAYHSSLAFTSLNCVADPILYCLVNEGARSDVAKALHNLLRFLASDKPQEMANASLTLETPLTSKRNSTAKAMTGSWAATPPSQGDQVQLKMLPPAQ | 0.53 | 0.7 | Intra-cellular |
| GPR6a | 2830 | NM_001286099 | NP_001273028 | MTLLAWCTRGANPAAMNASAASLNDSQVVVVAAEGAAAAATAAGGPDTGEWGPPAAAALGAGGGANGSLELSSQLSAGPPGLLLPAVNPWDVLLCVSGTVIAGENALVVALIASTPALRTPMFVLVGSLATADLLAGCGLILHFVFQYLVPSETVSLLTVGFLVASFAASVSSLLAITVDRYLSLYNALTYYSRRTLLGVHLLLAATWTVSLGLGLLPVLGWNCLAERAACSVVRPLARSHVALLSAAFFMVFGIMLHLYVRICQVVWRHAHQIALQQHCLAPPHLAATRKGVGTLAVVLGTFGASWLPFAIYCVVGSHEDPAVYTYATLLPATYNSMINPIIYAFRNQEIQRALWLLLCGCFQSKVPFRSRSPSEV | 0.39 | 0.63 | |
| GPR6b | 2830 | NM_005284 | NP_005275 | MNASAASLNDSQVVVVAAEGAAAAATAAGGPDTGEWGPPAAAALGAGGGANGSLELSSQLSAGPPGLLLPAVNPWDVLLCVSGTVIAGENALVVALIASTPALRTPMFVLVGSLATADLLAGCGLILHFVFQYLVPSETVSLLTVGFLVASFAASVSSLLAITVDRYLSLYNALTYYSRRTLLGVHLLLAATWTVSLGLGLLPVLGWNCLAERAACSVVRPLARSHVALLSAAFFMVFGIMLHLYVRICQVVWRHAHQIALQQHCLAPPHLAATRKGVGTLAVVLGTFGASWLPFAIYCVVGSHEDPAVYTYATLLPATYNSMINPIIYAFRNQEIQRALWLLLCGCFQSKVPFRSRSPSEV | 0.32 | 0.63 | |
| GPR12 | 2835 | NM_005288 | NP_005279 | MNEDLKVNLSGLPRDYLDAAAAENISAAVSSRVPAVEPEPELVVNPWDIVLCTSGTLISCENAIVVLIIFHNPSLRAPMFLLIGSLALADLLAGIGLITNFVFAYLLQSEATKLVTIGLIVASFSASVCSLLAITVDRYLSLYYALTYHSERTVTFTYVMLVMLWGTSICLGLLPVMGWNCLRDESTCSVVRPLTKNNAAILSVSFLFMFALMLQLYIQICKIVMRHAHQIALQHHFLATSHYVTTRKGVSTLAIILGTFAACWMPFTLYSLIADYTYPSIYTYATLLPATYNSIINPVIYAFRNQEIQKALCLICCGCIPSSLAQRARSPSDV | 0.38 | 0.68 | |
| GPR15 | 2838 | NM_005290 | NP_005281 | MDPEETSVYLDYYYATSPNSDIRETHSHVPYTSVFLPVFYTAVFLTGVLGNLVLMGALHFKPGSRRLIDIFIINLAASDFIFLVTLPLWVDKEASLGLWRTGSFLCKGSSYMISVNMHCSVLLLTCMSVDRYLAIVWPVVSRKFRRTDCAYVVCASIWFISCLLGLPTLLSRELTLIDDKPYCAEKKATPIKLIWSLVALIFTFFVPLLSIVTCYCCIARKLCAHYQQSGKHNKKLKKSIKIIFIVVAAFLVSWLPFNTFKFLAIVSGLRQEHYLPSAILQLGMEVSGPLAFANSCVNPFIYYIFDSYIRRAIVHCLCPCLKNYDFGSSTETSDSHLTKALSTFIHAEDFARRRKRSVSL | 0.28 | 0.67 | |
| GPR17a | 2840 | NM_005291 | NP_005282 | MSKRSWWAGSRKPPREMLKLSGSDSSQSMNGLEVAPPGLITNFSLATAEQCGQETPLENMLFASFYLLDFILALVGNTLALWLFIRDHKSGTPANVFLMHLAVADLSCVLVLPTRLVYHFSGNHWPFGEIACRLTGFLFYLNMYASIYFLTCISADRFLAIVHPVKSLKLRRPLYAHLACAFLWVVVAVAMAPLLVSPQTVQTNHTVVCLQLYREKASHHALVSLAVAFTFPFITTVTCYLLIIRSLRQGLRVEKRLKTKAVRMIAIVLAIFLVCFVPYHVNRSVYVLHYRSHGASCATQRILALANRITSCLTSLNGALDPIMYFFVAEKFRHALCNLLCGKRLKGPPPSFEGKTNESSLSAKSEL | 0.49 | 0.69 | |
| GPR17b | 2840 | NM_001161416 | NP_001154888 | MNGLEVAPPGLITNFSLATAEQCGQETPLENMLFASFYLLDFILALVGNTLALWLFIRD | 0.53 | 0.72 | Intra- |

| | | | | | | | |
|---|---|---|---|---|---|---|---|
| | | | | HKSGTPANVFLMHLAVADLSCVLVLPTRLVYHFSGNHWPFGEIACRLTGFLFYLNMYAS<br>IYFLTCISADRFLAIVHPVKSLKLRRPLYAHLACAFLWVVVAVAMAPLLVSPQTVQTNH<br>TVVCLQLYREKASHHALVSLAVAFTFPFITTVTCYLLIIRSLRQGLRVEKRLKTKAVRM<br>IAIVLAIFLVCFVPYHVNRSVYVLHYRSHGASCATQRILALANRITSCLTSLNGALDPI<br>MYFFVAEKFRHALCNLLCGKRLKGPPPSFEGKTNESSLSAKSEL | | | cellular |
| GPR19 | 2842 | NM_006143 | NP_006134 | MVFAHRMDNSKPHLIIPTLLVPLQNRSCTETATPLPSQYLMELSEEHSWMSNQTDLHYV<br>LKPGEVATASIFFGILWLFSIFGNSLVCLVIHRSRRTQSTTNYFVVSMACADLLISVAS<br>TPFVLLQFTTGRWTLGSATCKVVRYFQYLTPGVQIYVLLSICIDRFYTIVYPLSFKVSR<br>EKAKKMIAASWVFDAGFVTPVLFFYGSNWDSHCNYFLPSSWEGTAYTVIHFLVGFVIPS<br>VLIILFYQKVIKYIWRIGTDGRTVRRTMNIVPRTKVKTIKMFLILNLLFLLSWLPFHVA<br>QLWHPHEQDYKKSSLVFTAITWISFSSSASKPTLYSIYNANFRRGMKETFCMSSMKCYR<br>SNAYTITTSSRMAKKNYVGISEIPSMAKTITKDSIYDSFDREAKEKKLAWPINSNPPNT<br>FV | 0.11 | 0.56 | |
| GPR20 | 2843 | NM_005293 | NP_005284 | MPSVSPAGPSAGAVPNATAVTTVRTNASGLEVPLFHLFARLDEELHGTFPGLWLALMAV<br>HGAIFLAGLVLNGLALYVFCCRTRAKTPSVIYTINLVVTDLLVGLSLPTRFAVYYGARG<br>CLRCAFPHVLGYFLNMHCSILFLTCICVDRYLAIVRPEGSRRCRQPACARAVCAFVWLA<br>AGAVTLSVLGVTGSRPCCRVFALTVLEFLLPLLVISVFTGRIMCALSRPGLLHQGRQRR<br>VRAMQLLLTVLIIFLVCFTPFHARQVAVALWPDMPHHTSLVVYHVAVTLSSLNSCMDPI<br>VYCFVTSGFQATVRGLFGQHGEREPSSGDVVSMHRSSKGSGRHHILSAGPHALTQALAN<br>GPEA | 0.42 | 0.65 | |
| GPR21 | 2844 | NM_005294 | NP_005285 | MNSTLDGNQSSHPFCLLAFGYLETVNFCLLEVLIIVFLTVLIISGNIIVIFVFHCAPLL<br>NHHTTSYFIQTMAYADLFVGVSCVVPSLSLLHHPLPVEESLTCQIFGFVVSVLKSVSMA<br>SLACISIDRYIAITKPLTYNTLVTPWRLRLCIFLIWLYSTLVFLPSFFHWGKPGYHGDV<br>FQWCAESWHTDSYFTLFIVMMLYAPAALIVCFTYFNIFRICQQHTKDISERQARFSSQS<br>GETGEVQACPDKRYAMVLFRITSVFYILWLPYIIYFLLESSTGHSNRFASFLTTWLAIS<br>NSFCNCVIYSLSNSVFQRGLKRLSGAMCTSCASQTTANDPYTVRSKGPLNGCHI | 0.34 | 0.69 | |
| GPR22 | 2845 | NM_005295 | NP_005286 | MCFSPILEINMQSESNITVRDDIDDINTNMYQPLSYPLSFQVSLTGFLMLEIVLGLGSN<br>LTVLVLYCMKSNLINSVSNIITMNLHVLDVIICVGCIPLTIVILLLSLESNTALICCFH<br>EACVSFASVSTAINVFAITLDRYDISVKPANRILTMGRAVMLMISIWIFSFFSFLIPFI<br>EVNFFSLQSGNTWENKTLLCVSTNEYYTELGMYYHLLVQIPIFFFTVVVMLITYTKILQ<br>ALNIRIGTRFSTGQKKKARKKKTISLTTQHEATDMSQSSGGRNVVFGVRTSVSVIIALR<br>RAVKRHRERRERQKRVFRMSLLIISTFLLCWTPISVLNTTILCLGPSDLLVKLRLCFLV<br>MAYGTTIFHPLLYAFTRQKFQKVLKSKMKKRVVSIVEADPLPNNAVIHNSWIDPKRNKK<br>ITFEDSEIREKCLVPQVVTD | 0.22 | 0.57 | |
| GPR25 | 2848 | NM_005298 | NP_005289 | MAPTEPWSPSPGSAPWDYSGLDGLEELELCPAGDLPYGYVYIPALYLAAFAVGLLGNAF<br>VVWLLAGRRGPRRLVDTFVLHLAAADLGFVLTLPLWAAAAALGGRWPFGDGLCKLSSFA<br>LAGTRCAGALLLAGMSVDRYLAVVKLLEARPLRTPRCALASCCGVWAVALLAGLPSLVY<br>RGLQPLPGGQDSQCGEEPSHAFQGLSLLLLLLTFVLPLVVTLFCYCRISRRLRRPPHVG<br>RARRNSLRIIFAIESTFVGSWLPFSALRAVFHLARLGALPLPCPLLLALRWGLTIATCL<br>AFVNSCANPLIYLLLDRSFRARALDGACGRTGRLARRISSASSLSRDDSSVFRCRAQAA<br>NTASASW | 0.68<br>0.68<br>0.67<br>0.68<br>0.66<br>0.68 | 0.72<br>0.73<br>0.73<br>0.71<br>0.71<br>0.72 | |
| GPR26 | 2849 | NM_153442 | NP_703143 | MNSWDAGLAGLLVGTMGVSLLSNALVLLCLLHSADIRRQAPALFTLNLTCGNLLCTVVN<br>MPLTLAGVVAQRQPAGDRLCRLAAFLDTFLAANSMLSMAALSIDRWVAVVFPLSYRAKM<br>RLRDAALMVAYTWLHALTFPAAALALSWLGFHQLYASCTLCSRRPDERLRFAVFTGAFH<br>ALSFLLSFVVLCCTYLKVLKVARFHCKRIDVITMQTLVLLVDLHPSVRERCLEEQKRRR<br>QRATKKISTFIGTFLVCFAPYVITRLVELFSTVPIGSHWGVLSKCLAYSKAASDPFVYS<br>LLRHQYRKSCKEILNRLLHRRSIHSSGLTGDSHSQNILPVSE | 0.41 | 0.71 | |
| GPR27 | 2850 | NM_018971 | NP_061844 | MANASEPGGSGGGEAAALGLKLATLSLLLCVSLAGNVLFALLIVRERSLHRAPYYLLLD<br>LCLADGLRALACLPAVMLAARRAAAAAGAPPGALGCKLLAFLAALFCFHAAFLLLGVGV<br>TRYLAIAHHRFYAERLAGWPCAAMLVCAAWALALAAAFPPVLDGGGDDEDAPCALEQRP<br>DGAPGALGFLLLLAVVVGATHLVYLRLLFFIHDRRKMRPARLVPAVSHDWTFHGPGATG<br>QAAANWTAGFGRGPTPPALVGIRPAGPGRGARRLLVLEEFKTEKRLCKMFYAVTLLFLL<br>LWGPYVVASYLRVLVRPGAVPQAYLTASVWLTFAQAGINPVVCFLFNRELRDCFRAQFP<br>CCQSPRTTQATHPCDLKGIGL | 0.23 | 0.59 | |
| GPR31 | 2853 | NM_005299 | NP_005290 | MPFPNCSAPSTVVATAVGVLLGLECGLGLLGNAVALWTFLFRVRVWKPYAVYLLNLALA<br>DLLLAACLPFLAAFYLSLQAWHLGRVGCWALHFLLDLSRSVGMAFLAAVALDRYLRVVH<br>PRLKVNLLSPQAALGVSGLVWLLMVALTCPGLLISEAAQNSTRCHSFYSRADGSFSIIW<br>QEALSCLQFVLPFGLIVFCNAGIIRALQKRLREPEKQPKLQRAQALVTLVVVLFALCFL<br>PCFLARVLMHIFQNLGSCRALCAVAHTSDVTGSLTYLHSVLNPVVYCFSSPTFRSSYRR<br>VFHTLRGKGQAAEPPDFNPRDSYS | 0.34 | 0.7 | |

| | | | | | | |
|---|---|---|---|---|---|---|
| GPR32 | 2854 | NM_001506 | NP_001497 | MNGVSEGTRGCSDRQPGVLTRDRSCSRKMNSSGCLSEEVGSLRPLTVVILSASIVVGVLGNGLVLWMTVFRMARTVSTVCFFHLALADFMLSLSLPIAMYYIVSRQWLLGEWACKLYITFVFLSYFASNCLLVFISVDRCISVLYPVWALNHRTVQRASWLAFGVWLLAAALCSAHLKFRTTRKWNGCTHCYLAFNSDNETAQIWIEGVVEGHIIGTIGHFLLGFLGPLAIIGTCAHLIRAKLLREGWVHANRPKRLLLVLVSAFFIFWSPFNVVLLVHLWRRVMLKEIYHPRMLLILQASFALGCVNSSLNPFLYVFVGRDFQEKFFQSLTSALARAFGEEEFLSSCPRGNAPRE | 0.31 | 0.68 |
| GPR33 | 2856 | NM_001197184 | NP_001184113 | MDLINSTDYLINASTLVRNSTQFLAPASKMIIALSLYISSIIGTITNGLYLWVLRFKMKQTVNTLLFFHLILSYFISTMILPFMATSQLQDNHWNFGTALCKVFNGTLSLGMFTSVFFLSAIGLDRYLLTLHPVWSQQHRTPRWASSIVLGVWISAAALSIPYLIFRETHHDRKGKVTCQNNYAVSTNWESKEMQASRQWIHVACFISRFLLGFLLPFFIIIFCYERVASKVKERSLFKSSKPFKVMMTAIISFFVCWMPYHIHQGLLLTTNQSLLLELTLILTVLTTSFNTIFSPTLYLFVGENFKKVFKKSILALFESTFSEDSSVERTQT | 0.25 | 0.69 |
| GPR34 | 2857 | NM_005300 | NP_005291 | MRSHTITMTTTSVSSWPYSSHRMRFITNHSDQPPQNFSATPNVTTCPMDEKLLSTVLTTSYSVIFIVGLVGNIIALYVFLGIHRKRNSIQIYLLNVAIADLLLIFCLPFRIMYHINQNKWTLGVILCKVVGTLFYMNMYISIILLGFISLDRYIKINRSIQQRKAITTKQSIYVCCIVWMLALGGFLTMIILTLKKGGHNSTMCFHYRDKHNAKGEAIFNFILVVMFWLIFLLIILSYIKIGKNLLRISKRRSKFPNSGKYATTARNSFIVLIIFTICFVPYHAFRFIYISSQLNVSSCYWKEIVHKTNEIMLVLSSFNSCLDPVMYFLMSSNIRKIMCQLLFRRFQGEPSRSESTSEFKPGYSLHDTSVAVKIQSSSKST | 0.22 | 0.61 |
| GPR35a | 2859 | NM_005301 | NP_005292 | MNGTYNTCGSSDLTWPPAIKLGFYAYLGVLLVLGLLLNSLALWVFCCRMQQWTETRIYMTNLAVADLCLLCTLPFVLHSLRDTSDTPLCQLSQGIYLTNRYMSISLVTAIAVDRYVAVRHPLRARGLRSPRQAAAVCAVLWVLVIGSLVARWLLGIQEGGFCFRSTRHNFNSMAFPLLGFYLPLAVVVFCSLKVVTALAQRPPTDVGQAEATRKAARMVWANLLVFVVCFLPLHVGLTVRLAVGWNACALLETIRRALYITSKLSDANCCLDAICYYYMAKEFQEASALAVAPSAKAHKSQDSLCVTLA | 0.35<br>0.34<br>0.36 | 0.69<br>0.67<br>0.66 |
| GPR35b | 2859 | NM_001195381 | NP_001182310 | MLSGSRAVPTPHRGSEELLKYMLHSPCVSLTMNGTYNTCGSSDLTWPPAIKLGFYAYLGVLLVLGLLLNSLALWVFCCRMQQWTETRIYMTNLAVADLCLLCTLPFVLHSLRDTSDTPLCQLSQGIYLTNRYMSISLVTAIAVDRYVAVRHPLRARGLRSPRQAAAVCAVLWVLVIGSLVARWLLGIQEGGFCFRSTRHNFNSMAFPLLGFYLPLAVVVFCSLKVVTALAQRPPTDVGQAEATRKAARMVWANLLVFVVCFLPLHVGLTVRLAVGWNACALLETIRRALYITSKLSDANCCLDAICYYYMAKEFQEASALAVAPSAKAHKSQDSLCVTLA | 0.31<br>0.33<br>0.3 | 0.64<br>0.65<br>0.64 |
| GPR37 | 2861 | NM_005302 | NP_005293 | ALGVAPASRNETCLGESCAPTVIQRRGRDAWGPGNSARDVLRARAPREEQGAAFLAGPSWDLPAAPGRDPAAGRGAEASAAGPPGPPTRPPGPWRWKGARGQEPSETLGRGNPTALQLFLQISEEEEKGPRGAGISGRSQEQSVKTVPGASDLFYWPRRAGKLQGSHHKPLSKTANGLAGHEGWTIALPGRALAQNGSLGEGIHEPGGPRRGNSTNRRVRLKNPFYPLTQESYGAYAVMCLSVVIFGTGIIGNLAVMCIVCHNYYMRSISNSLLANLAFWDFLIIFFCLPLVIFHELTKKWLLEDFSCKIVPYIEVASLGVTTFTLCALCIDRFRAATNVQMYYEMIENCSSTTAKLAVIWVGALLLALPEVVLRQLSKEDLGFSGRAPAERCIIKISPDLPDTIYVLALTYDSARLWWYFGCYFCLPTLFTITCSLVTARKIRKAEKACTRGNKRQIQLESQMNCTVVALTILYGFCIIPENICNIVTAYMATGVSQQTMDLLNIISQFLLFFKSCVTPVLLFCLCKPFSRAFMECCCCCCEECIQKSSTVTSDDNDNEYTTELELSPFSTIRREMSTFASVGTHC | 0.14 | 0.5 |
| GPR37-s | 2861 | NM_005302 | NP_005293 | RLKNPFYPLTQESYGAYAVMCLSVVIFGTGIIGNLAVMCIVCHNYYMRSISNSLLANLAFWDFLIIFFCLPLVIFHELTKKWLLEDFSCKIVPYIEVASLGVTTFTLCALCIDRFRAATNVQMYYEMIENCSSTTAKLAVIWVGALLLALPEVVLRQLSKEDLGFSGRAPAERCIIKISPDLPDTIYVLALTYDSARLWWYFGCYFCLPTLFTITCSLVTARKIRKAEKACTRGNKRQIQLESQMNCTVVALTILYGFCIIPENICNIVTAYMATGVSQQTMDLLNIISQFLLFFKSCVTPVLLFCLCKPFSRAFMECCCCCCEECIQKSSTVTSDDNDNEYTTELELSPFSTIRREMSTFASVGTHC | 0.39 | 0.69 |
| GPR37L1 | 9283 | NM_004767 | NP_004758 | APLHLGRHRAETQEQQSRSKRGTEDEEAKGVQQYVPEEWAEYPRPIHPAGLQPTKPLVATSPNPGKDGGTPDSGQELRGNLTGAPGQRLQIQNPLYPVTESSYSAYAIMLLALVVFAVGIVGNLSVMCIVWHSYYLKSAWNSILASLALWDFLVLFFCLPIVIFNEITKQRLLGDVSCRAVPFMEVSSLGVTTFSLCALGIDRFHVATSTLPKVRPIERCQSILAKLAVIWVGSMTLAVPELLLWQLAQEPAPTMGTLDSCIMKPSASLPESLYSLVMTYQNARMWWYFGCYFCLPILFTVTCQLVTWRVRGPPGRKSECRASKHEQCESQLNSTVVGLTVVYAFCTLPENVCNIVVAYLSTELTRQTLDLLGLINQFSTFFKGAITPVLLLCICRPLGQAFLDCCCCCCCEECGGASEASAANGSDNKLKTEVSSSIYFHKPRESPPLLPLGTPC | 0.11 | 0.55 |
| GPR37L1-s | 9283 | NM_004767 | NP_004758 | NPLYPVTESSYSAYAIMLLALVVFAVGIVGNLSVMCIVWHSYYLKSAWNSILASLALWDFLVLFFCLPIVIFNEITKQRLLGDVSCRAVPFMEVSSLGVTTFSLCALGIDRFHVATSTLPKVRPIERCQSILAKLAVIWVGSMTLAVPELLLWQLAQEPAPTMGTLDSCIMKPSASL | 0.13 | 0.62 |

| | | | | | | |
|---|---|---|---|---|---|---|
| | | | | PESLYSLVMTYQNARMWWYFGCYFCLPILFTVTCQLVTWRVRGPPGRKSECRASKHEQCESQLNSTVVGLTVVYAFCTLPENVCNIVVAYLSTELTRQTLDLLGLINQFSTFFKGAITPVLLLCICRPLGQAFLDCCCCCCCEECGGASEASAANGSDNKLKTEVSSSIYFHKPRESPPLLPLGTPC | | |
| GPR39 | 2863 | NM_001508 | NP_001499 | MASPSLPGSDCSQIIDHSHVPEFEVATWIKITLILVYLIIFVMGLLGNSATIRVTQVLQKKGYLQKEVTDHMVSLACSDILVFLIGMPMEFYSIIWNPLTTSSYTLSCKLHTFLFEACSYATLLHVLTLSFERYIAICHPFRYKAVSGPCQVKLLIGFVWVTSALVALPLLFAMGTEYPLVNVPSHRGLTCNRSSTRHHEQPETSNMSICTNLSSRWTVFQSSIFGAFVVYLVVLLSVAFMCWNMMQVLMKSQKGSLAGGTRPPQLRKSESEESRTARRQTIIFLRLIVVTLAVCWMPNQIRRIMAAAKPKHDWTRSYFRAYMILLPFSETFFYLSSVINPLLYTVSSQQFRRVFVQVLCCRLSLQHANHEKRLRVHAHSTTDSARFVQRPLLFASRRQSSARRTEKIFLSTFQSEAEPQSKSQSLSLESLEPNSGAKPANSAAENGFQEHEV | 0.4 | 0.66 |
| GPR42 | 2866 | NM_001348195 | NP_001335124 | MDTGPDQSYFSGNHWFVFSVYLLTFLVGLPLNLLALVVFVGKLRCRPVAVDVLLLNLTASDLLLLLFLPFRMVEAANGMHWPLPFILCPLSGFIFFTTIYLTALFLAAVSIERFLSVAHPLWYKTRPRLGQAGLVSVACWLLASAHCSVVYVIEFSGDISHSQGTNGTCYLEFWKDQLAILLPVRLEMAVVLFVVPLIITSYCYSRLVWILGRGGSHRRQRRVAGLVAATLLNFLVCFGPYNVSHVVGYICGESPVWRIYVTLLSTLNSCVDPFVYYFSSSGFQADFHELLRRLCGLWGQWQQESSMELKEQKGGEEQRADRPAERKTSEHSQGCGTGGQVACAEN | 0.25 | 0.67 |
| GPR45 | 11250 | NM_007227 | NP_009158 | MACNSTSLEAYTYLLLNTSNASDSGSTQLPAPLRISLAIVMLLMTVVGFLGNTVVCIIVYQRPAMRSAINLLLATLAFSDIMLSLCCMPFTAVTLITVRWHFGDHFCRLSATLYWFFVLEGVAILLIISVDRFLIIVQRQDKLNPRRAKVIIAVSWVLSFCIAGPSLTGWTLVEVPARAPQCVLGYTELPADRAYVVTLVVAVFFAPFGVMLCAYMCILNTVRKNAVRVHNQSDSLDLRQLTRAGLRRLQRQQQVSVDLSFKTKAFTTILILFVGFSLCWLPHSVYSLLSVFSQRFYCGSSFYATSTCVLWLSYLKSVFNPIVYCWRIKKFREACIELLPQTFQILPKVPERIRRRIQPSTVYVCNENQSAV | 0.23 | 0.64 |
| GPR50 | 9248 | NM_004224 | NP_004215 | MGPTLAVPTPYGCIGCKLPQPEYPPALIIFMFCAMVITIVVDLIGNSMVILAVTKNKKLRNSGNIFVVSLSVADMLVAIYPYPLMLHAMSIGGWDLSQLQCQMVGFITGLSVVGSIFNIVAIAINRYCYICHSLQYERIFSVRNTCIYLVITWIMTVLAVLPNMYIGTIEYDPRTYTCIFNYLNNPVFTVTIVCIHFVLPLLIVGFCYVRIWTKVLAARDPAGQNPDNQLAEVRNFLTMFVIFLLFAVCWCPINVLTVLVAVSPKEMAGKIPNWLYLAAYFIAYFNSCLNAVIYGLLNENFRREYWTIFHAMRHPIIFFSGLISDIREMQEARTLAR | 0.2 | 0.68 |
| GPR52 | 9293 | NM_005684 | NP_005675 | MNESRWTEWRILNMSSGIVNVSERHSCPLGFGHYSVVDVCIFETVVIVLLTFLIIAGNLTVIFVFHCAPLLHHYTTSYFIQTMAYADLFVGVSCLVPTLSLLHYSTGVHESLTCQVFGYIISVLKSVSMACLACISVDRYLAITKPLSYNQLVTPCRLRICIILIWIYSCLIFLPSFFGWGKPGYHGDIFEWCATSWLTSAYFTGFIVCLLYAPAAFVVCFTYFHIFKICRQHTKEINDRRARFPSHEVDSSRETGHSPDRRYAMVLFRITSVFYMLWLPYIIYFLLESSRVLDNPTLSFLTTWLAISNSFCNCVIYSLSNSVFRLGLRRLSETMCTSCMCVKDQEAQEPKPRKRANSCSI | 0.36 | 0.69 |
| GPR61 | 83873 | NM_031936 | NP_114142 | MESSPIPQSSGNSSTLGRVPQTPGPSTASGVPEVGLRDVASESVALFFMLLLDLTAVAGNAAVMAVIAKTPALRKFVFVFHLCLVDLLAALTLMPLAMLSSSALFDHALFGEVACRLYLFLSVCFVSLAILSVSAINVERYYYVVHPMRYEVRMTLGLVASVLVGVWVKALAMASVPVLGRVSWEEGAPSVPPGCSLQWSHSAYCQLFVVVFAVLYFLLPLLLILVVYCSMFRVARVAAMQHGPLPTWMETPRQRSESLSSRSTMVTSSGAPQTTPHRTFGGGKAAVVLLAVGGQFLLCWLPYFSFHLYVALSAQPISTGQVESVVTWIGYFCFTSNPFFYGCLNRQIRGELSKQFVCFFKPAPEEELRLPSREGSIEENFLQFLQGTGCPSESWVSRPLPSPKQEPPAVDFRIPGQIAEETSEFLEQQLTSDIIMSDSYLRPAASPRLES | 0.45 | 0.6 |
| GPR62 | 118442 | NM_080865 | NP_543141 | MANSTGLNASEVAGSLGLILAAVVEVGALLGNGALLVVVLRTPGLRDALYLAHLCVVDLLAAASIMPLGLLAAPPPGLGRVRLGPAPCRAARFLSAALLPACTLGVAALGLARYRLIVHPLRPGSRPPPVLVLTAVWAAAGLLGALSLLGTPPAPPPAPARCSVLAGGLGPFRPLWALLAFALPALLLLGAYGGIFVVARRAALRPPRPARGSRLHSDSLDSRLSILPPLRPRLPGGKAALAPALAVGQFAACWLPYGCACLAPAARAAEAEAAVTWVAYSAFAAHPFLYGLLQRPVRLALGRLSRRALPGPVRACTPQAWHPRALLQCLQRPPEGPAVGPSEAPEQTPELAGGRSPAYQGPPESSLS | 0.47 | 0.65 |
| GPR63 | 81491 | NM_030784 | NP_110411 | MVFSAVLTAFHTGTSNTTFVVYENTYMNITLPPPFQHPDLSPLLRYSFETMAPTGLSSLTVNSTAVPTTPAAFKSLNLPLQITLSAIMIFILFVSFLGNLVVCLMVYQKAAMRSAINILLASLAFADMLLAVLNMPFALVTILTTRWIFGKFFCRVSAMFFWLFVIEGVAILLIISIDRFLIIVQRQDKLNPYRAKVLIAVSWATSFCVAFPLAVGNPDLQIPSRAPQCVFGYTTNPGYQAYVILISLISFFIPFLVILYSFMGILNTLRHNALRIHSYPEGICLSQASKLGLMSLQRPFQMSIDMGFKTRAFTTILILFAVFIVCWAPFTTYSLVATFSKHFYYQHNFFEISTWLLWLCYLKSALNPLIYYWRIKKFHDACLDMMPKSFKFLPQLPGHTKRRIRPSAVYVCG | 0.23 | 0.58 |

| | | | | | | |
|---|---|---|---|---|---|---|
| | | | | EHRTVV | | |
| GPR65 | 8477 | NM_003608 | NP_003599 | MNSTCIEEQHDLDHYLFPIVYIFVIIVSIPANIGSLCVSFLQAKKESELGIYLFSLSLS<br>DLLYALTLPLWIDYTWNKDNWTFSPALCKGSAFLMYMNFYSSTAFLTCIAVDRYLAVVY<br>PLKFFFLRTRRFALMVSLSIWILETIFNAVMLWEDETVVEYCDAEKSNFTLCYDKYPLE<br>KWQINLNLFRTCTGYAIPLVTILICNRKVYQAVRHNKATENKEKKRIIKLLVSITVTFV<br>LCFTPFHVMLLIRCILEHAVNFEDHSNSGKRTYTMYRITVALTSLNCVADPILYCFVTE<br>TGRYDMWNILKFCTGRCNTSQRQRKRILSVSTKDTMELEVLE | 0.34 | 0.71 |
| GPR68 | 8111 | NM_003485 | NP_003476 | MGNITADNSSMSCTIDHTIHQTLAPVVYVTVLVVGFPANCLSLYFGYLQIKARNELGVY<br>LCNLTVADLFYICSLPFWLQYVLQHDNWSHGDLSCQVCGILLYENIYISVGFLCCISVD<br>RYLAVAHPFRFHQFRTLKAAVGVSVVIWAKELLTSIYFLMHEEVIEDENQHRVCFEHYP<br>IQAWQRAINYYRFLVGFLFPICLLLASYQGILRAVRRSHGTQKSRKDQIQRLVLSTVVI<br>FLACFLPYHVLLLVRSVWEASCDFAKGVFNAYHFSLLLTSFNCVADPVLYCFVSETTHR<br>DLARLRGACLAFLTCSRTGRAREAYPLGAPEASGKSGAQGEEPELLTKLHPAFQTPNSP<br>GSGGFPTGRLA | 0.49 | 0.68 |
| GPR75 | 10936 | NM_006794 | NP_006785 | MNSTGHLQDAPNATSLHVPHSQEGNSTSLQEGLQDLIHTATLVTCTFLLAVIFCLGSYG<br>NFIVFLSFFDPAFRKFRTNFDFMILNLSFCDLFICGVTAPMFTFVLFFSSASSIPDAFC<br>FTFHLTSSGFIIMSLKTVAVIALHRLRMVLGKQPNRTASFPCTVLLTLLLWATSFTLAT<br>LATLKTSKSHLCLPMSSLIAGKGKAILSLYVVDFTFCVAVVSVSYIMIAQTLRKNAQVR<br>KCPPVITVDASRPQPFMGVPVQGGGDPIQCAMPALYRNQNYNKLQHVQTRGYTKSPNQL<br>VTPAASRLQLVSAINLSTAKDSKAVVTCVIIVLSVLVCCLPLGISLVQVVLSSNGSFIL<br>YQFELFGFTLIFFKSGLNPFIYSRNSAGLRRKVLWCLQYIGLGFF | 0.25 | 0.63 |
| GPR78 | 27201 | NM_080819 | NP_543009 | MGPGEALLAGLLVMVLAVALLSNALVLLCCAYSAELRTRASGVLLVNLSLGHLLLAALD<br>MPFTLLGVMRGRTPSAPGACQVIGFLDTFLASNAALSVAALSADQWLAVGFPLRYAGRL<br>RPRYAGLLLGCAWGQSLAFSGAALGCSWLGYSSAFASCSLRLPPEPERPRFAAFTATLH<br>AVGFVLPLAVLCLTSLQVHRVARRHCQRMDTVTMKALALLADLHPSVRQRCLIQQKRRR<br>HRATRKIGIAIATFLICFAPYVMTRLAELVPFVTVNAQWGILSKCLTYSKAVADPFTYS<br>LLRRPFRQVLAGMVHRLLKRTPRPASTHDSSLDVAGMVHQLLKRTPRPASTHNGSVDTE<br>NDSCLQQTH | 0.5 | 0.72 |
| GPR82 | 27197 | NM_080817 | NP_543007 | MNNNTTCIQPSMISSMALPIIYILLCIVGVFGNTLSQWIFLTKIGKKTSTHIYLSHLVT<br>ANLLVCSAMPFMSIYFLKGFQWEYQSAQCRVVNFLGTLSMHASMFVSLLILSWIAISRY<br>ATLMQKDSSQETTSCYEKIFYGHLLKKFRQPNFARKLCIYIWGVVLGIIIPVTVYYSVI<br>EATEGEESLCYNRQMELGAMISQIAGLIGTTFIGFSFLVVLTSYYSFVSHLRKIRTCTS<br>IMEKDLTYSSVKRHLLVIQILLIVCFLPYSIFKPIFYVLHQRDNCQQLNYLIETKNILT<br>CLASARSSTDPIIFLLLDKTFKKTLYNLFTKSNSAHMQSYG | 0.28 | 0.67 |
| GPR83 | 10888 | NM_016540 | NP_057624 | TEPHEGRADEQSAEAALAVPNASHFFSWNNYTFSDWQNFVGRRRYGAESQNPTVKALLI<br>VAYSFIIVFSLFGNVLVCHVIFKNQRMHSATSLFIVNLAVADIMITLLNTPFTLVRFVN<br>STWIFGKGMCHVSRFAQYCSLHVSALTLTAIAVDRHQVIMHPLKPRISITKGVIYIAVI<br>WTMATFFSLPHAICQKLFTFKYSEDIVRSLCLPDFPEPADLFWKYLDLATFILLYILPL<br>LIISVAYARVAKKLWLCNMIGDVTTEQYFALRRKKKKTIKMLMLVVVLFALCWFPLNCY<br>VLLLSSKVIRTNNALYFAFHWFAMSSTCYNPFIYCWLNENFRIELKALLSMCQRPPKPQ<br>EDRPPSPVPSFRVAWTEKNDGQRAPLANNLLPTSQLQSGKTDLSSVEPIVTMS | 0.28 | 0.63 |
| GPR84 | 53831 | NM_020370 | NP_065103 | MWNSSDANFSCYHESVLGYRYVAVSWGVVVAVTGTVGNVLTLLALAIQPKLRTRFNLLI<br>ANLTLADLLYCTLLQPFSVDTYLHLHWRTGATFCRVFGLLLFASNSVSILTLCLIALGR<br>YLLIAHPKLFPQVFSAKGIVLALVSTWVVGVASFAPLWPIYILVPVVCTCSFDRIRGRP<br>YTTILMGIYFVLGLSSVGIFYCLIHRQVKRAAQALDQYKLRQASIHSNHVARTDEAMPG<br>RFQELDSRLASGGPSEGISSEPVSAATTQTLEGDSSEVGDQINSKRAKQMAEKSPPEAS<br>AKAQPIKGARRAPDSSSEFGKVTRMCFAVFLCFALSYIPFLLLNILDARVQAPRVVHML<br>AANLTWLNGCINPVLYAAMNRQFRQAYGSILKRGPRSFHRLH | 0.24 | 0.63 |
| GPR85 | 54329 | NM_018970 | NP_061843 | MANYSHAADNILQNLSPLTAFLKLTSLGFIIGVSVVGNLLISILLVKDKTLHRAPYYFL<br>LDLCCSDILRSAICFPFVFNSVKNGSTWTYGTLTCKVIAFLGVLSCFHTAFMLFCISVT<br>RYLAIAHHRFYTKRLTFWTCLAVICMVWTLSVAMAFPPVLDVGTYSFIREEDQCTFQHR<br>SFRANDSLGFMLLLALILLATQLVYLKLIFFVHDRRKMKPVQFVAAVSQNWTFHGPGAS<br>GQAAANWLAGFGRGPTPPTLLGIRQNANTTGRRRLLVLDEFKMEKRISRMFYIMTFLFL<br>TLWGPYLVACYWRVFARGPVVPGGFLTAAVWMSFAQAGINPFVCIFSNRELRRCFSTTL<br>LYCRKSRLPREPYCVI | 0.09 | 0.6 |
| GPR87 | 53836 | NM_023915 | NP_076404 | MGFNLTLAKLPNNELHGQESHNSGNRSDGPGKNTTLHNEFDTIVLPVLYLIIFVASILL<br>NGLAVWIFFHIRNKTSFIFYLKNIVVADLIMTLTFPFRIVHDAGFGPWYFKFILCRYTS<br>VLFYANMYTSIVFLGLISIDRYLKVVKPFGDSRMYSITFTKVLSVCVWVIMAVLSLPNI<br>ILTNGQPTEDNIHDCSKLKSPLGVKWHTAVTYVNSCLFVAVLVILIGCYIAISRYIHKS<br>SRQFISQSSRKRKHNQSIRVVVAVFFTCFLPYHLCRIPFTFSHLDRLLDESAQKILYYC | 0.25 | 0.65 |

| | | | | | | | |
|---|---|---|---|---|---|---|---|
| | | | | KEITLFLSACNVCLDPIIYFFMCRSFSRRLFKKSNIRTRSESIRSLQSVRRSEVRIYYDYTDV | | | |
| GPR88 | 54112 | NM_022049 | NP_071332 | MTNSSSTSTSSTTGGSLLLLCEEEESWAGRRIPVSLLYSGLAIGGTLANGMVIYLVSSFRKLQTTSNAFIVNGCAADLSVCALWMPQEAVLGLLPTGSAEPPADWDGAGGSYRLLRGGLLGLGLTVSLLSHCLVALNRYLLITRAPATYQALYQRRHTAGMLALSWALALGLVLLLPPWAPRPGAAPPRVHYPALLAAAALLAQTALLLHCYLGIVRRVRVSVKRVSVLNFHLLHQLPGCAAAAAAFPGAQHAPGPGGAAHPAQAQPLPPALHPRRAQRRLSGLSVLLLCCVFLLATQPLVWVSLASGFSLPVPWGVQAASWLLCCALSALNPLLYTWRNEEFRRSVRSVLPGVGDAAAAAVAATAVPAVSQAQLGTRAAGQHW | 0.2 | 0.59 | |
| GPR101 | 83550 | NM_054021 | NP_473362 | MTSTCTNSTRESNSSHTCMPLSKMPISLAHGIIRSTVLVIFLAASFVGNIVLALVLQRKPQLLQVTNRFIFNLLVTDLLQISLVAPWVVATSVPLFWPLNSHFCTALVSLTHLFAFASVNTIVVVSVDRYLSIIHPLSYPSKMTQRRGYLLLYGTWIVAILQSTPPLYGWGQAAFDERNALCSMIWGASPSYTILSVVSFIVIPLIVMIACYSVVFCAARRQHALLYNVKRHSLEVRVKDCVENEDEEGAEKKEEFQDESEFRRQHEGEVKAKEGRMEAKDGSLKAKEGSTGTSESSVEARGSEEVRESSTVASDGSMEGKEGSTKVEENSMKADKGRTEVNQCSIDLGEDDMEFGEDDINFSEDDVEAVNIPESLPPSRRNSNSNPPLPRCYQCKAAKVIFIIIFSYVLSLGPYCFLAVLAVWVDVETQVPQWVITIIIWLFFLQCCIHPYVYGYMHKTIKKEIQDMLKKFFCKEKPPKEDSHPDLPGTEGGTEGKIVPSYDSATFP | 0.51 | 0.58 | Intra-cellular |
| GPR132a | 29933 | NM_001278694 | NP_001265623 | MCPMLLKNGYNGNATPVTTTAPWASLGLSAKTCNNVSFEESRIVLVVVYSAVCTLGVPANCLTAWLALLQVLQGNVLAVYLLCLALCELLYTGTLPLWVIYIRNQHRWTLGLLACKVTAYIFFCNIYVSILFLCCISCDRFVAVVYALESRGRRRRRTAILISACIFILVGIVHYPVFQTEDKETCFDMLQMDSRIAGYYYARFTVGFAIPLSIIAFTNHRIFRSIKQSMGLSAAQKAKVKHSAIAVVVIFLVCFAPYHLVLLVKAAAFSYYRGDRNAMCGLEERLYTASVVFLCLSTVNGVADPIIYVLATDHSRQEVSRIHKGWKEWSMKTDVTRLTHSRDTEELQSPVALADHYTFSRPVHPPGSPCPAKRLIEESC | 0.4 | 0.65 | |
| GPR132b | 29933 | NM_001278695 | NP_001265624 | MPGNATPVTTTAPWASLGLSAKTCNNVSFEESRIVLVVVYSAVCTLGVPANCLTAWLALLQVLQGNVLAVYLLCLALCELLYTGTLPLWVIYIRNQHRWTLGLLACKVTAYIFFCNIYVSILFLCCISCDRFVAVVYALESRGRRRRRTAILISACIFILVGIVHYPVFQTEDKETCFDMLQMDSRIAGYYYARFTVGFAIPLSIIAFTNHRIFRSIKQSMGLSAAQKAKVKHSAIAVVVIFLVCFAPYHLVLLVKAAAFSYYRGDRNAMCGLEERLYTASVVFLCLSTVNGVADPIIYVLATDHSRQEVSRIHKGWKEWSMKTDVTRLTHSRDTEELQSPVALADHYTFSRPVHPPGSPCPAKRLIEESC | 0.45 | 0.67 | |
| GPR135 | 64582 | NM_022571 | NP_072093 | MEEPQPPRPPASMALLGSQHSGAPSAAGPPGGTSSAATAAVLSFSTVATAALGNLSDASGGGTAAAPGGGGLGGSGAAREAGAAVRRPLGPEAAPLLSHGAAVAAQALVLLLIFLLSSLGNCAVMGVIVKHRQLRTVTNAFILSLSLSDLLTALLCLPAAFLDLFTPPGGSAPAAAAGPWRGFCAASRFFSSCFGIVSTLSVALISLDRYCAIVRPPREKIGRRRALQLLAGAWLTALGFSLPWELLGAPRELAAAQSFHGCLYRTSPDPAQLGAAFSVGLVVACYLLPFLLMCFCHYHICKTVRLSDVRVRPVNTYARVLRFFSEVRTATTVLIMIVFVICCWGPYCFLVLLAAARQAQTMQAPSLLSVVAVWLTWANGAINPVIYAIRNPNISMLLGRNREEGYRTRNVDAFLPSQGPGLQARSRSRLRNRYANRLGACNRMSSSNPASGVAGDVAMWARKNPVVLFCREGPPEPVTAVTKQPKSEAGDTSL | 0.38 | 0.57 | |
| GPR139 | 124274 | NM_001002911 | NP_001002911 | MEHTHAHLAANSSLSWWSPGSACGLGFVPVVYYSLLLCLGLPANILTVIILSQLVARRQKSSYNYLLALAAADILVLFFIVFVDFLLEDFILNMQMPQVPDKIIEVLEFSSIHTSIWITVPLTIDRYIAVCHPLKYHTVSYPARTRKVIVSVYITCFLTSIPYYWWPNIWTEDYISTSVHHVLIWIHCFTVYLVPCSIFFILNSIIVYKLRRKSNFRLRGYSTGKTTAILFTITSIFATLWAPRIIMILYHLYGAPIQNRWLVHIMSDIANMLALLNTAINFFLYCFISKRFRTMAAATLKAFFKCQKQPVQFYTNHNFSITSSPWISPANSHCIKMLVYQYDKNGKPIKVSP | 0.29 | 0.63 | |
| GPR141 | 353345 | NM_181791 | NP_861456 | MPGHNTSRNSSCDPIVTPHLISLYFIVLIGGLVGVISILFLLVKMNTRSVTTMAVINLVVVHSVFLLTVPFRLTYLIKKTWMFGLPFCKFVSAMLHIHMYLTFLFYVVILVTRYLIFFKCKDKVEFYRKLHAVAASAGMWTLVIVIVVPLVVSRYGIHEEYNEEHCFKFHKELAYTYVKIINYMIVIFVIAVAVILLVFQVFIIMLMVQKLRHSLLSHQEFWAQLKNLFFIGVILVCFLPYQFFRIYYLNVVTHSNACNSKVAFYNEIFLSVTAISCYDLLLFVFGGSHWFKQKIIGLWNCVLCR | 0.19 | 0.65 | |
| GPR142a | 350383 | NM_181790 | NP_861455 | MSIMMLPMEQKIQWVPTSLQDITAVLGTEAYTEEDKSMVSHAQKSQHSCLSHSRWLRSPQVTGGSWDLRIRPSKDSSSFRQAQCLRKDPGANNHLESQGVRGTAGDADRELRGPSEKATAGQPRVTLLPTPHVSGLSQEFESHWPEIAERSPCVAGVIPVIYYSVLLGLGLPVSLLTAVALARLATRTRRPSYYYLLALTASDIIIQVVIVFAGFLLQGAVLARQVPQAVVRTANILEFAANHASVWIAILLTVDRYTALCHPLHHRAASSPGRTRRAIAAVLSAALLTGIPFYWWLDMWRDTDSPRTLDEVLKWAHCLTVYFIPCGVFLVTNSAIIHRLRRRGRSGLQPRVGKSTAILLGITTLFTLLWAPRVFVMLYHMYVAPVHRDWRVHLALDVANMVAMLHTAANFGL | 0.19 | 0.49 | |

| | | | | | | | |
|---|---|---|---|---|---|---|---|
| | | | | YCFVSKTFRATVRQVIHDAYLPCTLASQPEGMAAKPVMEPPGLPTGAEV | | | |
| GPR142b | 350383 | NM_001331076 | NP_001318005 | MLTGSCGDPQKKPQVTQDSGPQSMGLEGRETAGQPRVTLLPTPHVSGLSQEFESHWPEI<br>AERSPCVAGVIPVIYYSVLLGLGLPVSLLTAVALARLATRTRRPSYYYLLALTASDIII<br>QVVIVFAGFLLQGAVLARQVPQAVVRTANILEFAANHASVWIAILLTVDRYTALCHPLH<br>HRAASSPGRTRRAIAAVLSAALLTGIPFYWWLDMWRDTDSPRTLDEVLKWAHCLTVYFI<br>PCGVFLVTNSAIIHRLRRRGRSGLQPRVGKSTAILLGITTLFTLLWAPRVFVMLYHMYV<br>APVHRDWRVHLALDVANMVAMLHTAANFGLYCFVSKTFRATVRQVIHDAYLPCTLASQP<br>EGMAAKPVMEPPGLPTGAEV | 0.25 | 0.57 | |
| GPR146 | 115330 | NM_001303474 | NP_001290403 | MWSCSWFNGTGLVEELPACQDLQLGLSLLSLLGLVVGVPVGLCYNALLVLANLHSKASM<br>TMPDVYFVNMAVAGLVLSALAPVHLLGPPSSRWALWSVGGEVHVALQIPFNVSSLVAMY<br>STALLSLDHYIERALPRTYMASVYNTRHVCGFVWGGALLTSFSSLLFYICSHVSTRALE<br>CAKMQNAEAADATLVFIGYVVPALATLYALVLLSRVRREDTPLDRDTGRLEPSAHRLLV<br>ATVCTQFGLWTPHYLILLGHTVIISRGKPVDAHYLGLLHFVKDFSKLLAFSSSFVTPLL<br>YRYMNQSFPSKLQRLMKKLPCGDRHCSPDHMGVQQVLA | 0.16 | 0.62 | |
| GPR148 | 344561 | NM_207364 | NP_997247 | MGDELAPCPVGTTAWPALIQLISKTPCMPQAASNTSLGLGDLRVPSSMLYWLFLPSSLL<br>AAATLAVSPLLLVTILRNQRLRQEPHYLLPANILLSDLAYILLHMLISSSSLGGWELGR<br>MACGILTDAVFAACTSTILSFTAIVLHTYLAVIHPLRYLSFMSHGAAWKAVALIWLVAC<br>CFPTFLIWLSKWQDAQLEEQGASYILPPSMGTQPGCGLLVIVTYTSILCVLFLCTALIA<br>NCFWRIYAEAKTSGIWGQGYSRARGTLLIHSVLITLYVSTGVVFSLDMVLTRYHHIDSG<br>THTWLLAANSEVLMMLPRAMLTYLYLLRYRQLLGMVRGHLPSRRHQAIFTIS | 0.52 | 0.69 | Intra-cellular |
| GPR149 | 344758 | NM_001038705 | NP_001033794 | MSLFLSNLSTNDSSLWKENHNSTDLLNPPGTLNIYLFCLTCLMTFAALVGSIYSLISLL<br>KMQNRTVVSMLVASWSVDDLMSVLSVTIFMFLQWPNEVPGYFQFLCTTSALMYLCQGLS<br>SNLKATLLVSYNFYTMHRGVGSQTASRRSGQVLGVVLTVWAASLLLSALPLCGWGAFVR<br>TPWGCLVDCSSSYVLFLSIVYALAFGLLVGLSVPLTHRLLCSEEPPRLHSNYQEISRGA<br>SIPGTPPTAGRVVSLSPEDAPGPSLRRSGGCSPSSDTVFGPGAPAAAGAEACRRENRGT<br>LYGTRSFTVSVAQKRFALILALTKVVLWLPMMMHMVVQNVVGFQSLPLETFSFLLTLLA<br>TTVTPVFVLSKRWTHLPCGCIINCRQNAYAVASDGKKIKRKGFEFNLSFQKSYGIYKIA<br>HEDYYDDDENSIFYHNLMNSECETTKDPQRDNRNIFNAIKVEISTTPSLDSSTQRGINK<br>CTNTDITEAKQDSNNKKDAFSDKTGGDINYEETTFSEGPERRLSHEESQKPDLSDWEWC<br>RSKSERTPRQRSGYALAIPLCAFQGTVSLHAPTGKTLSLSTYEVSAEGQKITPASKKIE<br>VYRSKSVGHEPNSEDSSSTFVDTSVKIHLEVLEICDNEEALDTVSIISNISQSSTQVRS<br>PSLRYSRKENRFVSCDLGETASYSLFLPTSNPDGDINISIPDTVEAHRQNSKRQHQERD<br>GYQEEIQLLNKAYRKREEESKGS | 0.25 | 0.42 | |
| GPR150 | 285601 | NM_199243 | NP_954713 | MEDLFSPSILPPAPNISVPILLGWGLNLTLGQGAPASGPPSRRVRLVFLGVILVVAVAG<br>NTTVLCRLCGGGGPWAGPKRRKMDFLLVQLALADLYACGGTALSQLAWELLGEPRAATG<br>DLACRFLQLLQASGRGASAHLVVLIALERRRAVRLPHGRPLPARALAALGWLLALLLAL<br>PPAFVVRGDSPSPLPPPPPPTSLQPGAPPAARAWPGERRCHGIFAPLPRWHLQVYAFYE<br>AVAGFVAPVTVLGVACGHLLSVWWRHRPQAPAAAAPWSASPGRAPAPSALPRAKVQSLK<br>MSLLLALLFVGCELPYFAARLAAAWSSGPAGDWEGEGLSAALRVVAMANSALNPFVYLF<br>FQAGDCRLRRQLRKRLGSLCCAPQGGAEDEEGPRGHQALYRQRWPHPHYHHARREPLDE<br>GGLRPPPPRPRPLPCSCESAF | 0.15 | 0.55 | |
| GPR151 | 134391 | NM_194251 | NP_919227 | MLAAAFADSNSSSMNVSFAHLHFAGGYLPSDSQDWRTIIPALLVAVCLVGFVGNLCVIG<br>ILLHNAWKGKPSMIHSLILNLSLADLSLLLFSAPIRATAYSKSVWDLGWFVCKSSDWFI<br>HTCMAAKSLTIVVVAKVCFMYASDPAKQVSIHNYTIWSVLVAIWTVASLLPLPEWFFST<br>IRHHEGVEMCLVDVPAVAEEFMSMFGKLYPLLAFGLPLFFASFYFWRAYDQCKKRGTKT<br>QNLRNQIRSKQVTVMLLSIAIISALLWLPEWVAWLWVWHLKAAGPAPPQGFIALSQVLM<br>FSISSANPLIFLVMSEEFREGLKGVWKWMITKKPPTVSESQETPAGNSEGLPDKVPSPE<br>SPASIPEKEKPSSPSSGKGKTEKAEIPILPDVEQFWHERDTVPSVQDNDPIPWEHEDQE<br>TGEGVK | 0.12 | 0.58 | |
| GPR152 | 390212 | NM_206997 | NP_996880 | MDTTMEADLGATGHRPRTELDDEDSYPQGGWDTVFLVALLLLGLPANGLMAWLAGSQAR<br>HGAGTRLALLLLSLALSDFLFLAAAAFQILEIRHGGHWPLGTAACRFYYFLWGVSYSSG<br>LFLLAALSLDRCLLALCPHWYPGHRPVRLPLWVCAGVWVLATLFSVPWLVFPEAAVWWY<br>DLVICLDFWDSEELSLRMLEVLGGFLPFLLLLVCHVLTQATACRTCHRQQQPAACRGFA<br>RVARTILSAYVVLRLPYQLAQLLYLAFLWDVYSGYLLWEALVYSDYLILLNSCLSPFLC<br>LMASADLRTLLRSVLSSFAAALCEERPGSFTPTEPQTQLDSEGPTLPEPMAEAQSQMDP<br>VAQPQVNPTLQPRSDPTAQPQLNPTAQPQSDPTAQPQLNLMAQPQSDSVAQPQADTNVQ<br>TPAPAASSVPSPCDEASPTPSSHPTPGALEDPATPPASEGESPSSTPPEAAPGAGPT | 0.23 | 0.54 | |
| GPR153 | 387509 | NM_207370 | NP_997253 | MSDERRLPGSAVGWLVCGGLSLLANAWGILSVGAKQKKWKPLEFLLCTLAATHMLNVAV<br>PIATYSVVQLRRQRPDFEWNEGLCKVFVSTFYTLTLATCFSVTSLSYHRMWMVCWPVNY<br>RLSNAKKQAVHTVMGIWMVSFILSALPAVGWHDTSERFYTHGCRFIVAEIGLGFGVCFL | 0.13 | 0.46 | |

| | | | | | | |
|---|---|---|---|---|---|---|
| | | | | LLVGGSVAMGVICTAIALFQTLAVQVGRQADRRAFTVPTIVVEDAQGKRRSSIDGSEPAKTSLQTTGLVTTIVFIYDCLMGFPVLVVSFSSLRADASAPWMALCVLWCSVAQALLLPVFLWACDRYRADLKAVREKCMALMANDEESDDETSLEGGISPDLVLERSLDYGYGGDFVALDRMAKYEISALEGGLPQLYPLRPLQEDKMQYLQVPPTRRFSHDDADVWAAVPLPAFLPRWGSGEDLAALAHLVLPAGPERRRASLLAFAEDAPPSRARRSAESLLSLRPSALDSGPRGARDSPPGSPRRRPGPGPRSASASLLPDAFALTAFECEPQALRRPPGPFPAAPAAPDGADPGEAPTPPSSAQRSPGPRPSAHSHAGSLRPGLSASWGEPGGLRAAGGGGSTSSFLSSPSESSGYATLHSDSLGSAS | | |
| GPR160 | 26996 | NM_014373 | NP_055188 | MTALSSENCSFQYQLRQTNQPLDVNYLLFLIILGKILLNILTLGMRRKNTCQNFMEYFCISLAFVDLLLLVNISIILYFRDFVLLSIRFTKYHICLFTQIISFTYGFLHYPVFLTACIDYCLNFSKTTKLSFKCQKLFYFFTVILIWISVLAYVLGDPAIYQSLKAQNAYSRHCPFYVSIQSYWLSFFMVMILFVAFITCWEEVTTLVQAIRITSYMNETILYFPFSSHSSYTVRSKKIFLSKLIVCFLSTWLPFVLLQVIIVLLKVQIPAYIEMNIPWLYFVNSFLIATVYWFNCHKLNLKDIGLPLDPFVNWKCCFIPLTIPNLEQIEKPISIMIC | 0.28 | 0.65 |
| GPR161a | 23432 | NM_153832 | NP_722561 | MSLNSSLSCRKELSNLTEEEGGEGGVIITQFIAIIVITIFVCLGNLVIVVTLYKKSYLLTLSNKFVFSLTLSNFLLSVLVLPFVVTSSIRREWIFGVVWCNFSALLYLLISSASMLTLGVIAIDRYYAVLYPMVYPMKITGNRAVMALVYIWLHSLIGCLPPLFGWSSVEFDEFKWMCVAAWHREPGYTAFWQIWCALFPFLVMLVCYGFIFRVARVKARKVHCGTVVIVEEDAQRTGRKNSSTSTSSSGSRRNAFQGVVYSANQCKALITILVVLGAFMVTWGPYMVVIASEALWGKSSVSPSLETWATWLSFASAVCHPLIYGLWNKTVRKELLGMCFGDRYYREPFVQRQRTSRLFSISNRITDLGLSPHLTALMAGGQPLGHSSSTGDTGFSCSQDSGTDMMLLEDYTSDDNPPSHCTCPPKRRSSVTFEDEVEQIKEAAKNSILHVKAEVHKSLDSYAASLAKAIEAEAKINLFGEEALPGVLVTARTVPGGGFGGRRGSRTLVSQRLQLQSIEEGDVLAAEQR | 0.47 | 0.55 |
| GPR161b | 23432 | NM_001267609 | NP_001254538 | MSARGVVQHALPTPRRGALTMSLNSSLSCRKELSNLTEEEGGEGGVIITQFIAIIVITIFVCLGNLVIVVTLYKKSYLLTLSNKFVFSLTLSNFLLSVLVLPFVVTSSIRREWIFGVVWCNFSALLYLLISSASMLTLGVIAIDRYYAVLYPMVYPMKITGNRAVMALVYIWLHSLIGCLPPLFGWSSVEFDEFKWMCVAAWHREPGYTAFWQIWCALFPFLVMLVCYGFIFRVARVKARKVHCGTVVIVEEDAQRTGRKNSSTSTSSSGSRRNAFQGVVYSANQCKALITILVVLGAFMVTWGPYMVVIASEALWGKSSVSPSLETWATWLSFASAVCHPLIYGLWNKTVRKELLGMCFGDRYYREPFVQRQRTSRLFSISNRITDLGLSPHLTALMAGGQPLGHSSSTGDTGFSCSQDSGTDMMLLEDYTSDDNPPSHCTCPPKRRSSVTFEDEVEQIKEAAKNSILHVKAEVHKSLDSYAASLAKAIEAEAKINLFGEEALPGVLVTARTVPGGGFGGRRGSRTLVSQRLQLQSIEEGDVLAAEQR | 0.5 | 0.53 |
| GPR161c | 23432 | NM_001267611 | NP_001254540 | MKVVQHALPTPRRGALTMSLNSSLSCRKELSNLTEEEGGEGGVIITQFIAIIVITIFVCLGNLVIVVTLYKKSYLLTLSNKFVFSLTLSNFLLSVLVLPFVVTSSIRREWIFGVVWCNFSALLYLLISSASMLTLGVIAIDRYYAVLYPMVYPMKITGNRAVMALVYIWLHSLIGCLPPLFGWSSVEFDEFKWMCVAAWHREPGYTAFWQIWCALFPFLVMLVCYGFIFRVARVKARKVHCGTVVIVEEDAQRTGRKNSSTSTSSSGSRRNAFQGVVYSANQCKALITILVVLGAFMVTWGPYMVVIASEALWGKSSVSPSLETWATWLSFASAVCHPLIYGLWNKTVRKELLGMCFGDRYYREPFVQRQRTSRLFSISNRITDLGLSPHLTALMAGGQPLGHSSSTGDTGFSCSQDSGTDMMLLEDYTSDDNPPSHCTCPPKRRSSVTFEDEVEQIKEAAKNSILHVKAEVHKSLDSYAASLAKAIEAEAKINLFGEEALPGVLVTARTVPGGGFGGRRGSRTLVSQRLQLQSIEEGDVLAAEQR | 0.33 | 0.51 |
| GPR162 | 27239 | NM_019858 | NP_062832 | MARGGAGAEEASLRSNALSWLACGLLALLANAWIILSISAKQQKHKPLELLLCFLAGTHILMAAVPLTTFAVVQLRRQASSDYDWNESICKVFVSTYYTLALATCFTVASLSYHRMWMVRWPVNYRLSNAKKQALHAVMGIWMVSFILSTLPSIGWHNNGERYYARGCQFIVSKIGLGFGVCFSLLLLGGIVMGLVCVAITFYQTLWARPRRARQARRVGGGGGTKAGGPGALGTRPAFEVPAIVVEDARGKRRSSLDGSESAKTSLQVTNLVSAIVFLYDSLTGVPILVVSFFSLKSDSAPPWMVLAVLWCSMAQTLLLPSFIWSCERYRADVRTVWEQCVAIMSEEDGDDDGGCDDYAEGRVCKVRFDANGATGPGSRDPAQVKLLPGRHMLFPPLERVHYLQVPLSRRLSHDETNIFSTPREPGSFLHKWSSSDDIRVLPAQSRALGGPPEYLGQRHRLEDEDEEEAEGGGLASLRQFLESGVLGSGGGPPRGPGFFREEITTFIDETPLPSPTASPGHSPRRPRPLGLSPRRLSLGSPESRAVGLPLGLSAGRRCSLTGGEESARAWGGSWGPGNPIFPQLTL | 0.18 | 0.48 |
| GPR171 | 29909 | NM_013308 | NP_037440 | MTNSSFFCPVYKDLEPFTYFFYLVFLVGIIGSCFATWAFIQKNTNHRCVSIYLINLLTADFLLTLALPVKIVVDLGVAPWKLKIFHCQVTACLIYINMYLSIIFLAFVSIDRCLQLTHSCKIYRIQEPGFAKMISTVVWLMVLLIMVPNMMIPIKDIKEKSNVGCMEFKKEFGRNWHLLTNFICVAIFLNFSAIILISNCLVIRQLYRNKDNENYPNVKKALINILLVTTGYIICFVPYHIVRIPYTLSQTEVITDCSTRISLFKAKEATLLLAVSNLCFDPILYYHLSKAFRSKVTETFASPKETKAQKEKLRCENNA | 0.28 | 0.7 |
| GPR173 | 54328 | NM_018969 | NP_061842 | MANTTGEPEEVSGALSPPSASAYVKLVLLGLIMCVSLAGNAILSLLVLKERALHKAPYY | 0.08 | 0.57 |

| | | | | | | |
|---|---|---|---|---|---|---|
| | | | | FLLDLCLADGIRSAVCFPFVLASVRHGSSWTFSALSCKIVAFMAVLFCFHAAFMLFCIS<br>VTRYMAIAHHRFYAKRMTLWTCAAVICMAWTLSVAMAFPPVFDVGTYKFIREEDQCIFE<br>HRYFKANDTLGFMLMLAVLMAATHAVYGKLLLFEYRHRKMKPVQMVPAISQNWTFHGPG<br>ATGQAAANWIAGFGRGPMPPTLLGIRQNGHAASRRLLGMDEVKGEKQLGRMFYAITLLF<br>LLLWSPYIVACYWRVFVKACAVPHRYLATAVWMSFAQAAVNPIVCFLLNKDLKKCLRTH<br>APCWGTGGAPAPREPYCVM | | |
| GPR174 | 84636 | NM_032553 | NP_115942 | MPANYTCTRPDGDNTDFRYFIYAVTYTVILVPGLIGNILALWVFYGYMKETKRAVIFMI<br>NLAIADLLQVLSLPLRIFYYLNHDWPFGPGLCMFCFYLKYVNMYASIYFLVCISVRRFW<br>FLMYPFRFHDCKQKYDLYISIAGWLIICLACVLFPLLRTSDDTSGNRTKCFVDLPTRNV<br>NLAQSVVMMTIGELIGFVTPLLIVLYCTWKTVLSLQDKYPMAQDLGEKQKALKMILTCA<br>GVFLICFAPYHFSFPLDFLVKSNEIKSCLARRVILIFHSVALCLASLNSCLDPVIYYFS<br>TNEFRRRLSRQDLHDSIQLHAKSFVSNHTASTMTPELC | 0.42 | 0.7 |
| GPR176a | 11245 | NM_007223 | NP_009154 | MGHNGSWISPNASEPHNASGAEAAGVNRSALGEFGEAQLYRQFTTTVQVVIFIGSLLGN<br>FMVLWSTCRTTVFKSVTNRFIKNLACSGICASLVCVPFDIILSTSPHCCWWIYTMLFCK<br>VVKFLHKVFCSVTILSFPAIALDRYYSVLYPLERKISDAKSRELVMYIWAHAVVASVPV<br>FAVTNVADIYATSTCTEVWSNSLGHLVYVLVYNITTVIVPVVVVFLFLILIRRALSASQ<br>KKKVIIAALRTPQNTISIPYASQREAELHATLLSMVMVFILCSVPYATLVVYQTVLNVP<br>DTSVFLLLTAVWLPKVSLLANPVLFLTVNKSVRKCLIGTLVQLHHRYSRRNVVSTGSGM<br>AEASLEPSIRSGSQLLEMFHIGQQQIFKPTEDEEESEAKYIGSADFQAKEIFSTCLEGE<br>QGPQFAPSAPPLSTVDSVSQVAPAAPVEPETFPDKYSLQFGFGPFELPPQWLSETRNSK<br>KRLLPPLGNTPEELIQTKVPKVGRVERKMSRNNKVSIFPKVDS | 0.32 | 0.53 |
| GPR176b | 11245 | NM_001271854 | NP_001258783 | MGHNGSWISPNASEPHNASGAEAAGVNRSALGEFGEAQLYRQFTTTVQVVIFIGSLLEF<br>GNMEVTRKLDKSRLPGIKFIKNLACSGICASLVCVPFDIILSTSPHCCWWIYTMLFCKV<br>VKFLHKVFCSVTILSFPAIALDRYYSVLYPLERKISDAKSRELVMYIWAHAVVASVPVF<br>AVTNVADIYATSTCTEVWSNSLGHLVYVLVYNITTVIVPVVVVFLFLILIRRALSASQK<br>KKVIIAALRTPQNTISIPYASQREAELHATLLSMVMVFILCSVPYATLVVYQTVLNVPD<br>TSVFLLLTAVWLPKVSLLANPVLFLTVNKSVRKCLIGTLVQLHHRYSRRNVVSTGSGMA<br>EASLEPSIRSGSQLLEMFHIGQQQIFKPTEDEEESEAKYIGSADFQAKEIFSTCLEGEQ<br>GPQFAPSAPPLSTVDSVSQVAPAAPVEPETFPDKYSLQFGFGPFELPPQWLSETRNSKK<br>RLLPPLGNTPEELIQTKVPKVGRVERKMSRNNKVSIFPKVDS | 0.27 | 0.49 |
| GPR182 | 11318 | NM_007264 | NP_009195 | MSVKPSWGPGPSEGVTAVPTSDLGEIHNWTELLDLFNHTLSECHVELSQSTKRVVLFAL<br>YLAMFVVGLVENLLVICVNWRGSGRAGLMNLYILNMAIADLGIVLSLPVWMLEVTLDYT<br>WLWGSFSCRFTHYFYFVNMYSSIFFLVCLSVDRYVTLTSASPSWQRYQHRVRRAMCAGI<br>WVLSAIIPLPEVVHIQLVEGPEPMCLFMAPFETYSTWALAVALSTTILGFLLPFPLITV<br>FNVLTACRLRQPGQPKSRRHCLLLCAYVAVFVMCWLPYHVTLLLLTLHGTHISLHCHLV<br>HLLYFFYDVIDCFSMLHCVINPILYNFLSPHFRGRLLNAVVHYLPKDQTKAGTCASSSS<br>CSTQHSIIITKGDSQPAAAAPHPEPSLSFQAHHLLPNTSPISPTQPLTPS | 0.21 | 0.61 |
| GPR183 | 1880 | NM_004951 | NP_004942 | MDIQMANNFTPPSATPQGNDCDLYAHHSTARIVMPLHYSLVFIIGLVGNLLALVVIVQN<br>RKKINSTTLYSTNLVISDILFTTALPTRIAYYAMGFDWRIGDALCRITALVFYINTYAG<br>VNFMTCLSIDRFIAVVHPLRYNKIKRIEHAKGVCIFVWILVFAQTLPLLINPMSKQEAE<br>RITCMEYPNFEETKSLPWILLGACFIGYVLPLIIILICYSQICCKLFRTAKQNPLTEKS<br>GVNKKALNTIILIIVVFVLCFTPYHVAIIQHMIKKLRFSNFLECSQRHSFQISLHFTVC<br>LMNFNCCMDPFIYFFACKGYKRKVMRMLKRQVSVSISSAVKSAPEENSREMTETQMMIH<br>SKSSNGK | 0.24 | 0.67 |
| MAS1 | 4142 | NM_002377 | NP_002368 | MDGSNVTSFVVEEPTNISTGRNASVGNAHRQIPIVHWVIMSISPVGFVENGILLWFLCFRMRRNPFTV<br>YITHLSIADISLLFCIFILSIDYALDYELSSGHYYTIVTLSVTFLFGYNTGLYLLTAISVERCLSVLY<br>PIWYRCHRPKYQSALVCALLWALSCLVTTMEYVMCIDREEESHSRNDCRAVIIFIAILSFLVFTPLML<br>VSSTILVVKIRKNTWASHSSKLYIVIMVTIIIFLIFAMPMRLLYLLYYEYWSTFGNLHHISLLFSTIN<br>SSANPFIYFFVGSSKKKRFKESLKVVLTRAFKDEMQPRRQKDNCNTVTVETVV | 0.26 | 0.67 |
| MAS1L | 116511 | NM_052967 | NP_443199 | MVWGKICWFSQRAGWTVFAESQISLSCSLCLHSGDQEAQNPNLVSQLCGVFLQNETNETIHMQMSMAV<br>GQQALPLNIIAPKAVLVSLCGVLLNGTVFWLLCCGATNPYMVYILHLVAADVIYLCCSAVGFLQVTLL<br>TYHGVVFFIPDFLAILSPFSFEVCLCLLVAISTERCVCVLFPIWYRCHRPKYTSNVVCTLIWGLPFCI<br>NIVKSLFLTYWKHVKACVIFLKLSGLFHAILSLVMCVSSLTLLIRFLCCSQQQKATRVYAVVQISAPM<br>FLLWALPLSVAPLITDFKMFVTTSYLISLFLIINSSANPIIYFFVGSLRKKRLKESLRVILQRALADK<br>PEVGRNKKAAGIDPMEQPHSTQHVENLLPREHRVDVET | 0.23 | 0.6 |
| MRGPRD | 116512 | NM_198923 | NP_944605 | MNQTLNSSGTVESALNYSRGSTVHTAYLVLSSLAMFTCLCGMAGNSMVIWLLGFRMHRNPFCIYILNL<br>AAADLLFLFSMASTLSLETQPLVNTTDKVHELMKRLMYFAYTVGLSLLTAISTQRCLSVLFPIWFKCH<br>RPRHLSAWVCGLLWTLCLLMNGLTSSFCSKFLKFNEDRCFRVDMVQAALIMGVLTPVMTLSSLTLFVW<br>VRRSSQQWRRQPTRLFVVVLASVLVFLICSLPLSIYWFVLYWLSLPPEMQVLCFSLSRLSSSVSSSAN<br>PVIYFLVGSRRSHRLPTRSLGTVLQQALREEPELEGGETPTVGTNEMGA | 0.59 | 0.74 |
| MRGPRE | 116534 | NM_001039165 | NP_001034254 | MMEPREAGQHVGAANGAQEDVAFNLIILSLTEGLGLGGLLGNGAVLWLLSSNVYRNPFAIYLLDVACA<br>DLIFLGCHMVAIVPDLLQGRLDFPGFVQTSLATLRFFCYIVGLSLLAAVSVEQCLAALFPAWYSCRRP<br>RHLTTCVCALTWALCLLLHLLLSGACTQFFGEPSRHLCRTLWLVAAVLLALLCCTMCGASLMLLLRVE<br>RGPQRPPPRGFPGLILLTVLLFLFCGLPFGIYWLSRNLLWYIPHYFYHFSFLMAAVHCAAKPVVYFCL<br>GSAQGRRLPLRLVLQRALGDEAELGAVRETSRRGLVDIAA | 0.19 | 0.66 |
| MRGPRF | 116535 | NM_145015 | NP_659452 | MAGNCSWEAHPGNRNKMCPGLSEAPELYSRGFLTIEQIAMLPPPAVMNYIFLLLCLCGLVGNGLVLWF<br>FGFSIKRNPFSIYFLHLASADVGYLFSKAVFSILNTGGFLGTFADYIRSVCRVLGLCMFLTGVSLLPA | 0.1 | 0.61 |

| | | | | | | |
|---|---|---|---|---|---|---|
| | | | | VSAERCASVIFPAWYWRRRPKRLSAVVCALLWVLSLLVTCLHNYFCVFLGRGAPGAACRHMDIFLGILLFLLCCPLMVLPCLALILHVECRARRRQRSAKLNHVILAMVSVFLVSSIYLGIDWFLFWVFQIPAPFPEYVTDLCICINSSAKPIVYFLAGRDKSQRLWEPLRVVFQRALRDGAELGEAGGSTPNTVTMEMQCPPGNAS | | | |
| MRGPRG | 386746 | NM_001164377 | NP_001157849 | MFGLFGLWRTFDSVVFYLTLIVGLGGPVGNGLVLWNLGFRIKKGPFSIYLLHLAAADFLFLSCRVGFSVAQAALGAQDTLYFVLTFLWFAVGLWLLAAFSVERCLSDLFPACYQGCRPRHASAVLCALVWTPTLPAVPLPANACGLLRNSACPLVCPRYHVASVTWFLVLARVAWTAGVVLFVWVTCCSTRPRPRLYGIVLGALLLLFFCGLPSVFYWSLQPLLNFLLPVFSPLATLLACVNSSSKPLIYSGLGRQPGKREPLRSVLRRALGEGAELGARGQSLPMGLL | 0.18 | 0.66 | |
| MRGPRX1 | 259249 | NM_147199 | NP_671732 | MDPTISTLDTELTPINGTEETLCYKQTLSLTVLTCIVSLVGLTGNAVVLWLLGCRMRRNAFSIYILNLAAADFLFLSGRLIYSLLSFISIPHTISKILYPVMMFSYFAGLSFLSAVSTERCLSVLWPIWYRCHRPTHLSAVVCVLLWALSLLRSILEWMLCGFLFSGADSAWCQTSDFITVAWLIFLCVVLCGSSLVLLIRILCGSRKIPLTRLYVTILLTVLVFLLCGLPFGIQFFLFLWIHVDREVLFCHVHLVSIFLSALNSSANPIIYFFVGSFRQRQNRQNLKLVLQRALQDASEVDEGGGQLPEEILELSGSRLEQ | 0.42 | 0.71 | |
| MRGPRX2 | 117194 | NM_054030 | NP_473371 | MDPTTPAWGTESTTVNGNDQALLLLCGKETLIPVFLILFIALVGLVGNGFVLWLLGFRMRRNAFSVYVLSLAGADFLFLCFQIINCLVYLSNFFCSISINFPSFFTTVMTCAYLAGLSMLSTVSTERCLSVLWPIWYRCRRPRHLSAVVCVLLWALSLLLSILEGKFCGFLFSDGDSGWCQTFDFITAAWLIFLFMVLCGSSLALLVRILCGSRGLPLTRLYLTILLTVLVFLLCGLPFGIQWFLILWIWKDSDVLFCHIHPVSVVLSSLNSSANPIIYFFVGSFRKQWRLQQPILKLALQRALQDIAEVDHSEGCFRQGTPEMSRSSLV | 0.28<br>0.3<br>0.48 | 0.68<br>0.65<br>0.72 | |
| MRGPRX3 | 117195 | NM_054031 | NP_473372 | MDSTIPVLGTELTPINGREETPCYKQTLSFTGLTCIVSLVALTGNAVVLWLLGCRMRRNAVSIYILNLVAADFLFLSGHIICSPLRLINIRHPISKILSPVMTFPYFIGLSMLSAISTERCLSILWPIWYHCRRPRYLSSVMCVLLWALSLLRSILEWMFCDFLFSGANSVWCETSDFITIAWLVFLCVVLCGSSLVLLVRILCGSRKMPLTRLYVTILLTVLVFLLCGLPFGIQWALFSRIHLDWKVLFCHVHLVSIFLSALNSSANPIIYFFVGSFRQRQNRQNLKLVLQRALQDTPEVDEGGGWLPQETLELSGSRLEQ | 0.27 | 0.68 | |
| MRGPRX4 | 117196 | NM_054032 | NP_473373 | MDPTVPVFGTKLTPINGREETPCYNQTLSFTVLTCIISLVGLTGNAVVLWLLGYRMRRNAVSIYILNLAAADFLFLSFQIIRLPLRLINISHLIRKILVSVMTFPYFTGLSMLSAISTERCLSVLWPIWYRCRRPTHLSAVVCVLLWGLSLLFSMLEWRFCDFLFSGADSSWCETSDFIPVAWLIFLCVVLCVSSLVLLVRILCGSRKMPLTRLYVTILLTVLVFLLCGLPFGILGALIYRMHLNLEVLYCHVYLVCMSLSSLNSSANPIIYFFVGSFRQRQNRQNLKLVLQRALQDKPEVDKGEGQLPEESLELSGSRLGP | 0.31 | 0.69 | |
| P2RY8 | 286530 | NM_178129 | NP_835230 | MQVPNSTGPDNATLQMLRNPAIAVALPVVYSLVAAVSIPGNLFSLWVLCRRMGPRSPSVIFMINLSVTDLMLASVLPFQIYYHCNRHHWVFGVLLCNVVTVAFYANMYSSILTMTCISVERFLGVLYPLSSKRWRRRRYAVAACAGTWLLLLTALSPLARTDLTYPVHALGIITCFDVLKWTMLPSVAMWAVFLFTIFILLFLIPFVITVACYTATILKLLRTEEAHGREQRRRAVGLAAVVLLAFVTCFAPNNFVLLAHIVSRLFYGKSYYHVYKLTLCLSCLNNCLDPFVYYFASREFQLRLREYLGCRRVPRDTLDTRRESLFSARTTSVRSEAGAHPEGMEGATRPGLQRQESVF | 0.57 | 0.7 | Intra-cellular |
| P2RY10 | 27334 | NM_014499 | NP_055314 | MANLDKYTETFKMGSNSTSTAEIYCNVTNVKFQYSLYATTYILIFIPGLLANSAALWVLCRFISKKNKAIIFMINLSVADLAHVLSLPLRIYYYISHHWPFQRALCLLCFYLKYLNMYASICFLTCISLQRCFFLLKPFRARDWKRRYDVGISAAIWIVVGTACLPFPILRSTDLNNNKSCFADLGYKQMNAVALVGMITVAELAGFVIPVIIIAWCTWKTTISLRQPPMAFQGISERQKALRMVFMCAAVFFICFTPYHINFIFYTMVKETIISSCPVVRIALYFHPFCLCLASLCCLLDPILYYFMASEFRDQLSRHGSSVTRSRLMSKESGSSMIG | 0.48 | 0.72 | Intra-cellular |
| TAAR2 | 9287 | NM_014626 | NP_055441 | MYSFMAGSIFITIFGNLAMIISISYFKQLHTPTNFLILSMAITDFLLGFTIMPYSMIRSVENCWYFGLTFCKIYYSFDLMLSITSIFHLCSVAIDRFYAICYPLLYSTKITIPVIKRLLLLCWSVPGAFAFGVVFSEAYADGIEGYDILVACSSSCPVMFNKLWGTTLFMAGFFTPGSMMVGIYGKIFAVSRKHAHAINNLRENQNNQVKKDKKAAKTLGIVIGVFLLCWFPCFFTILLDPFLNFSTPVVLFDALTWFGYFNSTCNPLIYGFFYPWFRRALKYILLGKIFSSCFHNTILCMQKESE | 0.28 | 0.71 | |
| TAAR5 | 9038 | NM_003967 | NP_003958 | MRAVFIQGAEEHPAAFCYQVNGSCPRTVHTLGIQLVIYLACAAGMLIIVLGNVFVAFAVSYFKALHTPTNFLLLSLALADMFLGLLVLPLSTIRSVESCWFFGDFLCRLHTYLDTLFCLTSIFHLCFISIDRHCAICDPLLYPSKFTVRVALRYILAGWGVPAAYTSLFLYTDVVETRLSQWLEEMPCVGSCQLLLNKFWGWLNFPLFFVPCLIMISLYVKIFVVATRQAQQITTLSKSLAGAAKHERKAAKTLGIAVGIYLLCWLPFTIDTMVDSLLHFITPPLVFDIFIWFAYFNSACNPIIYVFSYQWFRKALKLTLSQKVFSPQTRTVDLYQE | 0.19 | 0.69 | |
| TAAR6 | 319100 | NM_175067 | NP_778237 | MSSNSSLLVAVQLCYANVNGSCVKIPFSPGSRVILYIVFGFGAVLAVFGNLLVMISILHFKQLHSPTNFLVASLACADFLVGVTVMPFSMVRTVESCWYFGRSFCTFHTCCDVAFCYSSLFHLCFISIDRYIAVTDPLVYPTKFTVSVSGICISVSWILPLMYSGAVFYTGVYDDGLEELSDALNCIGGCQTVVNQNWVLTDFLSFFIPTFIMIILYGNIFLVARRQAKKIENTGSKTESSSESYKARVARRERKAAKTLGVTVVAFMISWLPYSIDSLIDAFMGFITPACIYEICCWCAYYNSAMNPLIYALFYPWFRKAIKVIVTGQVLKNSSATMNLFSEHI | 0.22 | 0.71 | |
| TAAR8 | 83551 | NM_053278 | NP_444508 | MTSNFSQPVVQLCYEDVNGSCIETPYSPGSRVILYTAFSFGSLLAVFGNLLVMTSVLHFKQLHSPTNFLIASLACADFLVGVTVMLFSMVRTVESCWYFGAKFCTLHSCCDVAFCYSSVLHLCFICIDRYIVVTDPLVYATKFTVSVSGICISVSWILPLTYSGAVFYTGVNDDGLEELVSALNCVGGCQIIVSQGWVLIDFLLFFIPTLVMIILYSKIFLIAKQQAIKIETTSSKVESSSESYKIRVAKRERKAAKTLGVTVLAFVISWLPYTVDILIDAFMGFLTPAYIYEICCWSAYYNSAMNPLIYALFYPWFRKAIKLILSGDVLKASSSTISLFLE | 0.35 | 0.76 | |
| TAAR9 | 134860 | NM_175057 | NP_778227 | MVNNFSQAEAVELCYKNVNESCIKTPYSPGPRSILYAVLGFGAVLAAFGNLLVMIAILHFKQLHTPTNFLIASLACADFLVGVTVMPFSTVRSVESCWYFGDSYCKFHTCFDTSFCFASLFHLCCISVDRYIAVTDPLTYPTKFTVSVSGICIVLSWFFSVTYSFSIFYTGANEEGIEELVVALTCVGGCQAPLNQNWVLLCFLLFFIPNVAMVFIYSKIFLVAKHQARKIESTASQAQSSSESYKERVAKRERKAAKTLGIAMAAFLVSWLPYLVDAVIDAYMNFITPPYVYEILVWCVYYNSAMNPLIYAFFYQWFGKAIKLIVSGKVLRTDSSTTNLFSEEVETD | 0.27 | 0.73 | |
| GPR156 | 165829 | NM_153002 | NP_694547 | MEPEINCSELCDSFPGQELDRRPLHDLCKTTITSSHHSSKTISSLSPVLLGIVWTFLSCGLLLILFFLAFTIHCRKNRIVKMSSPNLNIVTLLGSCLTYSSAYLFGIQDVLVGSSMETLIQTRLSMLCIGTSLVFGPILGKSWRLYKVFTQRVPDKRVIIKDLQLLGLVAALLMADVILLMTWVLTDPIQCLQILSVSMTVTGKDVSCTSTSTHFCASRYSDVWIALIWGCKGLLLLYGAYLAGLTGHVSSPPVNQSLTIMVGVNLLVLAAGLLFVVTRYLHSWPNLVFGLTSGGIFVCTTTINCFIFIPQLKQWKAFEEENQTIRRMAKYFSTPNKSFHTQYGEEENCHPRGEKSSMERLLTEKNAVIESLQEQVNNAKEKIVRLMSAECTYDLPEGAAPPASSPNKDVQAVASVHTLAAAQGPSGHLSDFQNDPGMAARDSQCTSGPSSYAQSLEGPGKDSSFSPGKEEKISDSKDFSDHLDSGCSQKPWTEQSLGPERGDQVPMNPSQSLLPERGGSDPQRQRHLENSEEPPERRSRVSSVIREKLQEVLQDLGLGPEASLSTAPSCHQQTWKNSAAFSPQKMPLSKELGFSPYMVRRRRAAQRARSHFPGSAPSSVGHRANRTVPGAHSRLHVQNGDSPSLAPQTTDSRVRRPSSRKPSLPSDPQDRPGTLEGSKQSQTEPEGARGSKAAFLRQPSGSGRAPSPAAPCLSKASPDLPEQWQLWPPVPSGCASLSSQHSYFDTESSSSDEFFCRCHRPYCEICFQSSSDSSDSGTSDTDPEPTGGLASWEKLWARSKPIVNFKDDLKPTLV | 0.32 | 0.41 | |
| GPR158 | 57512 | NM_020752 | NP_065803 | ASRDPQGRPDSPRERTPKGKPHAQQPGRASASDSSAPWSRSTDGTILAQKLAEEVPMDVASYLYTGDSHQLKRANCSGRYELAGLPGKWPALASAHPSLHRALDTLTHATNFLNVMLQSNKSREQNLQDDLDWYQALVWSLLEGEPSISRAAITFSTDSLSAPAPQVFLQATREESRILLQDLSSSAPHLANATLETEWFHGLRRKWRPHLHRRGPNQGPRGLGHSWRRKDGLGGDKSHFKWSPPYLECENGSYKPGWLVTLSSAIYGLQPNLVPEFRGVMKVDINLQKVDIDQCSSDGWFSGTHKCHLNNSECMPIKGLGFVLGAYECICKAGFYHPGV | 0.17 | 0.38 | |

| | | | | | | |
|---|---|---|---|---|---|---|
| | | | | LPVNNFRRRGPDQHISGSTKDVSEEAYVCLPCREGCPFCADDSPCFVQEDKYLRLAIISFQALCMLLD<br>FVSMLVVYHFRKAKSIRASGLILLETILFGSLLLYFPVVILYFEPSTFRCILLRWARLLGFATVYGTV<br>TLKLHRVLKVFLSRTAQRIPYMTGGRVMRMLAVILLVVFWFLIGWTSSVCQNLEKQISLIGQGKTSDH<br>LIFNMCLIDRWDYMTAVAEFLFLLWGVYLCYAVRTVPSAFHEPRYMAVAVHNELIISAIFHTIRFVLA<br>SRLQSDWMLMLYFAHTHLTVTVTIGLLLIPKFSHSSNNPRDDIATEAYEDELDMGRSGSYLNSSINSA<br>WSEHSLDPEDIRDELKKLYAQLEIYKRKKMITNNPHLQKKRCSKKGLGRSIMRRITEIPETVSRQCSK<br>EDKEGADHGTAKGTALIRKNPPESSGNTGKSKEETLKNRVFSLKKSHSTYDHVRDQTEESSSLPTESQ<br>EEETTENSTLESLSGKKLTQKLKEDSEAESTESVPLVCKSASAHNLSSEKKTGHPRTSMLQKSLSVIA<br>SAKEKTLGLAGKTQTAGVEERTKSQKPLPKDKETNRNHSNSDNTETKDPAPQNSNPAEEPRKPQKSGI<br>MKQQRVNPTTANSDLNPGTTQMKDNFDIGEVCPWEVYDLTPGPVPSESKVQKHVSIVASEMEKNPTFS<br>LKEKSHHKPKAAEVCQQSNQKRIDKAEVCLWESQGQSILEDEKLLISKTPVLPERAKEENGGQPRAAN<br>VCAGQSEELPPKAVASKTENENLNQIGHQEKKTSSSEENVRGSYNSSNNFQQPLTSRAEVCPWEFETP<br>AQPNAGRSVALPASSALSANKIAGPRKEEIWDSFKV | | | |
| GPR179 | 440435 | NM_001004334 | NP_001004334 | LGGPRPIRSLPPLSSQVKPGSVPMQVPLEGAEAALAYLYSGDAQQLSQVNCSERYEARGAGAMPGLPP<br>SLQGAAGTLAQAANFLNMLLQANDIRESSVEEDVEWYQALVRSVAEGDPRVYRALLTFNPPPGASHLQ<br>LALQATRTGEETILQDLSGNWVQEENPPGDLDTPALKKRVLTNDLGSLGSPKWPQADGYVGDTQQVRL<br>SPPFLECQEGRLRPGWLITLSATFYGLKPDLSPEVRGQVQMDVDLQSVDINQCASGPGWYSNTHLCDL<br>NSTQCVPLESQGFVLGRYLCRCRPGFYGASPSGGLEESDFQTTGQFGFPEGRSGRLLQCLPCPEGCTS<br>CMDATPCLVEEAAVLRAAVLACQACCMLAIFLSMLVSYRCRRNKRIWASGVVLLETVLFGFLLLYFPV<br>FILYFKPSVFRCIALRWVRLLGFAIVYGTIILKLYRVLQLFLSRTAQRSALLSSGRLLRHLGLLLLPV<br>LGFLAVWTVGALERGIQHAPLVIRGHTPSGRHFYLCHHDRWDYIMVVAELLLLCWGSFLCYATRAVLS<br>AFHEPRYMGIALHNELLLSAAFHTARFVLVPSLHPDWTLLLFFFHTHSTVTTTLALIFIPKFWKLGAP<br>PREEMVDEVCEDELDLQHSGSYLGSSIASAWSEHSLDPGDIRDELKKLYAQLEVHKTKEMAANNPHLP<br>KKRGSSCQGLGRSFMRYLAEFPEALARQHSRDSGSPGHGSLPGSSRRRLLSSSLQEPEGTPALHKSRS<br>TYDQRREQDPPLLDSLLRRKLAKKASRTESRESVEGPPALGFRSASAHNLTVGERLPRARPASLQKSL<br>SVASSREKALLMASQAYLEETYRQAKEREERKKAKAAMASLVRRPSARRLERPRGAPLSAPPSPAKSS<br>SVDSSHTSGRLHEEARRRLPHPPIRHQVSTPILALSGGLGEPRMLSPTSTLAPALLPALAPTPAPALA<br>PVPVSPQSPNLLTYICPWENAELPAKQENVPQEGPSGPERGHHSPAPARARLWRALSVAVEKSRAGEN<br>EMDAEDAHHQREANDVDEDRPKIFPKSHSLKAPVQQGSMRSLGLAIKALTRSRSTYREKESVEESPEG<br>QNSGTAGESMGAPSRSPRLGRPKAVSKQAALIPSDDKESLQNQQNAHTSRMLQVCQREGSREQEDRGR<br>RMTQGLGERKAERAGKTGLAMLRQVSRDKNIKQSKETPVGWQELPKAGLQSLGSADHRVAEVCPWEVT<br>ESETRQPDSGNKAEICPWETSEGAPESRALRQDPGDSQKKRGEARGKSEPIDVVPMMRKKPERLVREQ<br>EAVCPWESADRGGLSPGSAPQDPGRIRDKSEAGDSVEARKVEKPGWEAAGPEAHTPDITKAEPCPWEA<br>SEGGEDGKPAQEAVKDLPQEKQKTRKATFWKEQKPGGDLESLCPWESTDFRGPSAVSIQAPGSSECSG<br>SLGSGIAEVCLWEAGDAPAIQKAEICPWELDDNVMGQEMLSLGTGRESLQEKEKASRKGSFGEMGEQT<br>VKAVQKLSQQQESVCPRESTVPGHSSPCLDNSSSKAGSQFLCNGGSRATQVCPQEDLRPEAQEATPAK<br>TEICPWEVNERTREEWTSAQVPRGGESQKDKEKMPGKSEIEDVTAWEKPEGQIQKQEAVGPWESVDPG<br>SFSPQPRPQDTERPQTLLQMSGSVGSKAADICPLDVEENLTAGKAEICPWEVGAGAGEERALGAEAIR<br>KSPNDTGKVSADLGPRERAVTAPEKPQKPTPEWEVACPWGSVGPGACSQHPGTLDADGPKAGFQELDH<br>MGCRPGEVCPWEAQEAATSEKAKICPWEVSEGTTGKGLDQKAGSESAEQREKALEKGRLTSLGEDVSK<br>GMAKLCQQQETICIWENKDLRESPAQAPKISDLPSSMSSEVAEGHSLEATEKGDLRQDPKTGSFPEHI<br>TQEKAPAADTEEFTTEDGEKTSHELQSVCPWETTAPADSVSHLDRQRPDQPKASSQRLVSTGGRAADV<br>CPWDVPDAGVYKSDSSAKAETCPWEVTERIPVKGVSRQDGKGDSQEEKGRAPEKSEPKGVPVQKKPEM<br>ADFRQQEAVCPWESQDGKGLSPQPAPDASDRSRGSSEAAGSVETRVAEVCLWEVVEAPSAKKAEICPW<br>EAGGGAAEEGEQERESQGQGEMFLQKAGPGGTEEHFSKAAAKPREQEAVCPGEGTGSGGLLPQSGALD<br>PELKVSPKEAGSMGSRMAELCQWEITDPEGNKIKGTMADICPGEETGVPSEESGLLALTATRREFFPT<br>APEKPLCLLVHGPLDHFFPESKIPCPKVSRPASTFTLEGVRELQGPSGLEPRTSLAPEPSLQEAESQS<br>SSLTEDSGQVAFEAQYEEFTPPTVYPWDWE | 0.58 | 0.34 | Intra-cellular |
| GPRC5A | 9052 | NM_003979 | NP_003970 | MATTVPDGCRNGLKSKYYRLCDKAEAWGIVLETVATAGVVTSVAFMLTLPILVCKVQDSNRRKMLPTQ<br>FLFLLGVLGIFGLTFAFIIGLDGSTGPTRFFLFGILFSICFSCLLAHAVSLTKLVRGRKPLSLLVILG<br>LAVGFSLVQDVIAIEYIVLTMNRTNVNVFSELSAPRRNEDFVLLLTYVLFLMALTFLMSSFTFCGSFT<br>GWKRHGAHIYLTMLLSIAIWVAWITLLMLPDFDRRWDDTILSSALAANGWVFLLAYVSPEFWLLTKQR<br>NPMDYPVEDAFCKPQLVKKSYGVENRAYSQEEITQGFEETGDTLYAPYSTHFQLQNQPPQKEFSIPRA<br>HAWPSPYKDYEVKKEGS | 0.1 | 0.56 | |
| GPRC5B | 51704 | NM_016235 | NP_057319 | ENASTSRGCGLDLLPQYVSLCDLDAIWGIVVEAVAGAGALITLLLMLILLVRLPFIKEKEKKSPVGLH<br>FLFLLGTLGLFGLTFAFIIQEDETICSVRRFLWGVLFALCFSCLLSQAWRVRRLVRHGTGPAGWQLVG<br>LALCLMLVQVIIAVEWLVLTVLRDTRPACAYEPMDFVMALIYDMVLLVVTLGLALFTLCGKFKRWKLN<br>GAFLLITAFLSVLIWVAWMTMYLFGNVKLQQGDAWNDPTLAITLAASGWVFVIFHAIPEIHCTLLPAL<br>QENTPNYFDTSQPRMRETAFEEDVQLPRAYMENKAFSMDEHNAALRTAGFPNGSLGKRPSGSLGKRPS<br>APFRSNVYQPTEMAVVLNGGTIPTAPPSHTGRHLW | 0.1 | 0.55 | |
| GPRC5C | 55890 | NM_018653 | NP_061123 | QGHVPPGCSQGLNPLYYNLCDRSGAWGIVLEAVAGAGIVTTFVLTIILVASLPFVQDTKKRSLLGTQV<br>FFLLGTLGLFCLVFACVVKPDFSTCASRRFLFGVLFAICFSCLAAHVFALNFLARKNHGPRGWVIFTV<br>ALLLTLVEVIINTEWLIITLVRGSGEGGPQGNSSAGWAVASPCAIANMDFVMALIYVMLLLLGAFLGA<br>WPALCGRYKRWRKHGVFVLLTTATSVAIWVVWIVMYTYGNKQHNSPTWDDPTLAIALAANAWAFVLFY<br>VIPEVSQVTKSSPEQSYQGDMYPTRGVGYETILKEQKGQSMFVENKAFSMDEPVAAKRPVSPYSGYNG<br>QLLTSVYQPTEMALMHKVPSEGAYDIILPRATANSQVMGSANSTLRAEDMYSAQSHQAATPPKDGKNS<br>QVFRNPYVWD | 0.11 | 0.55 | |
| GPRC5D | 55507 | NM_018654 | NP_061124 | MYKDCIESTGDYFLLCDAEGPWGIILESLAILGIVVTILLLLAFLFLMRKIQDCSQWNVLPTQLLFLL<br>SVLGLFGLAFAFIIELNQQTAPVRYFLFGVLFALCFSCLLAHASNLVKLVRGCVSFSWTTILCIAIGC<br>SLLQIIIATEYVTLIMTRGMMFVNMTPCQLNVDFVVLLVYVLFLMALTFFVSKATFCGPCENWKQHGR<br>LIFITVLFSIIIWVVWISMLLRGNPQFQRQPQWDDPVVCIALVTNAWVFLLLYIVPELCILYRSCRQE<br>CPLQGNACPVTAYQHSFQVENQELSRARDSDGAEEDVALTSYGTPIQPQTVDPTQECFIPQAKLSPQQ<br>DAGGV | 0.18 | 0.57 | |
| GPRC6A | 222545 | NM_148963 | NP_683766 | QPCQTPDDFVAATSPGHIIIGGLFAIHEKMLSSEDSPRRPQIQECVGFEISVFLQTLAMIHSIEMINN<br>STLLPGVKLGYEIYDTCTEVTVAMAATLRFLSKFNCSRETVEFKCDYSSYMPRVKAVIGSGYSEITMA<br>VSRMLNLQLMPQVGYESTAEILSDKIRFPSFLRTVPSDFHQIKAMAHLIQKSGWNWIGIITTDDDYGR<br>LALNTFIIQAEANNVCIAFKEVLPAFLSDNTIEVRINRTLKKIILEAQVNVIVVFLRQFHVFDLFNKA<br>IEMNINKMWIASDNWSTATKITTIPNVKKIGKVVGFAFRRGNISSFHSFLQNLHLLPSDSHKLLHEYA<br>MHLSACAYVKDTDLSQCIFNHSQRTLAYKANKAIERNFVMRNDFLWDYAEPGLIHSIQLAVFALGYAI<br>RDLCQARDCQNPNAFQPWELLGVLKNVTFTDGWNSFHFDAHGDLNTGYDVVLWKEINGHMTVTKMAEY<br>DLQNDVFIIPDQETKNEFRNLKQIQSKCSKECSPGQMKKTTRSQHICCYECQNCPENHYTNQTDMPHC<br>LLCNNKTHWAPVRSTMCFEKEVEYLNWNDSLAILLLILSLLGIIFVLVVGIIFTRNLNTPVVKSSGGL<br>RVCYVILLCHFLNFASTSFFIGEPQDFTCKTRQTMFGVSFTLCISCILTKSLKILLAFSFDPKLQKFL<br>KCLYRPILIIFTCTGIQVVICTLWLIFAAPTVEVNVSLPRVIILECEEGSILAFGTMLGYIAILAFIC<br>FIFAFKGKYENYNEAKFITFGMLIYFIAWITFIPIYATTFGKYVPAVEIIVILISNYGILYCTFIPKC | 0.46 | 0.67 | Intra-cellular |

| | | | | | | |
|---|---|---|---|---|---|---|
| | | | | YVIICKQEINTKSAFLKMIYSYSSHSVSSIALSPASLDSMSGNVTMTNPSSSGKSATWQKSKDLQAQAFAHICRENATSVSKTLPRKRMSSI | | |
| GPR107 | 57720 | NM_020960 | NP_066011 | RVHHLALKDDVRHKVHLNTFGFFKDGYMVVNVSSLSLNEPEDKDVTIGFSLDRTKNDGFSSYLDEDVNYCILKKQSVSVTLLILDISRSEVRVKSPPEAGTQLPKIIFSRDEKVLGQSQEPNVNPASAGNQTQKTQDGGKSKRSTVDSKAMGEKSFSVHNNGGAVSFQFFFNISTDDQEGLYSLYFHKCLGKELPSDKFTFSLDIEITEKNPDSYLSAGEIPLPKLYISMAFFFFLSGTIWIHILRKRRNDVFKIHWLMAALPFTKSLSLVFHAIDYHYISSQGFPIEGWAVVYYITHLLKGALLFITIALIGTGWAFIKHILSDKDKKIFMIVIPLQVLANVAYIIIESTEEGTTEYGLWKDSLFLVDLLCCGAILFPVVWSIRHLQEASATDGKAAINLAKLKLFRHYYVLIVCYIYFTRIIAFLLKLAVPFQWKWLYQLLDETATLVFFVLTGYKFRPASDNPYLQLSQEEEDLEMESVVTTSGVMESMKKVKKVTNGSVEPQGEWEGAV | 0.15 | 0.63 |
| GPR137 | 56834 | NM_020155 | NP_064540 | MESNLSGLVPAAGLVPALPPAVTLGLTAAYTTLYALLFFSVYAQLWLVLLYGHKRLSYQTVFLALCLLWAALRTTLFSFYFRDTPRANRLGPLPFWLLYCCPVCLQFFTLTLMNLYFAQVVFKAKVKRRPEMSRGLLAVRGAFVGASLLFLLVNVLCAVLSHRRRAQPWALLLVRVLVSDSLFVICALSLAACLCLVARRAPSTSIYLEAKGTSVCQAAAMGGAMVLLYASRACYNLTALALAPQSRLDTFDYDWYNVSDQADLVNDLGNKGYLVFGLILFVWELLPTTLLVGFFRVHRPPQDLSTSHILNGQVFASRSYFFDRAGHCEDEGCSWEHSRGESTRCQDQAATTTVSTPPHRRDPPPSPTEYPGPSPPHPRPLCQVCLPLLAQDPGGRGYPLLWPAPCCSCHSELVPSP | 0.25 | 0.54 |
| TPRA1 | 131601 | NM_001136053 | NP_001129525 | MDTLEEVTWANGSTALPPPLAPNISVPHRCLLLLYEDIGTSRVRYWDLLLLIPNVLFLIFLLWKLPSARAKIRITSSPIFITFYILVFVVALVGIARAVVSMTVSTSNAATVADKILWEITRFFLLAIELSVIILGLAFGHLESKSSIKRVLAITTVLSLAYSVTQGTLEILYPDAHLSAEDFNIYGHGGRQFWLVSSCFFFLVYSLVVILPKTPLKERISLPSRRSFYVYAGILALLNLLQGLGSVLLCFDIIEGLCCVDATTFLYFSFFAPLIYVAFLRGFFGSEPKILFSYKCQVDETEEPDVHLPQPYAVARREGLEAAGAAGASAASYSSTQFDSAGGVAYLDDIASMPCHTGSINSTDSERWKAINA | 0.14 | 0.51 |
| GPR143 | 4935 | NM_000273 | NP_000264 | MASPRLGTFCCPTRDAATQLVLSFQPRAFHALCLGSGGLRLALGLLQLLPGRRPAGPGSPATSPPASVRILRAAAACDLLGCLGMVIRSTVWLGFPNFVDSVSDMNHTEIWPAAFCVGSAMWIQLLYSACFWWLFCYAVDAYLVIRRSAGLSTILLYHIMAWGLATLLCVEGAAMLYYPSVSRCERGLDHAIPHYVTMYLPLLLVLVANPILFQKTVTAVASLLKGRQGIYTENERRMGAVIKIRFFKIMLVLIICWLSNIINESLLFYLEMQTDINGGSLKPVRTAAKTTWFIMGILNPAQGFLLSLAFYGWTGCSLGFQSPRKEIQWESLTTSAAEGAHPSPLMPHENPASGKVSQVGGQTSDEALSMLSEGSDASTIEIHTASESCNKNEGDPALPTHGDL | 0.37 | 0.64 |
| GPR151 | 134391 | NM_194251 | NP_919227 | MLAAAFADSNSSSMNVSFAHLHFAGGYLPSDSQDWRTIIPALLVAVCLVGFVGNLCVIGILLHNAWKGKPSMIHSLILNLSLADLSLLLFSAPIRATAYSKSVWDLGWFVCKSSDWFIHTCMAAKSLTIVVVAKVCFMYASDPAKQVSIHNYTIWSVLVAIWTVASLLPLPEWFFSTIRHHEGVEMCLVDVPAVAEEFMSMFGKLYPLLAFGLPLFFASFYFWRAYDQCKKRGTKTQNLRNQIRSKQVTVMLLSIAIISALLWLPEWVAWLWVWHLKAAGPAPPQGFIALSQVLMFSISSANPLIFLVMSEEFREGLKGVWKWMITKKPPTVSESQETPAGNSEGLPDKVPSPESPASIPEKEKPSSPSSGKGKTEKAEIPILPDVEQFWHERDTVPSVQDNDPIPWEHEDQETGEGVK | 0.24 | 0.6 |

**Table S4.** AlphaFold 3-predicted possible binding of CXCL17–GPR25 pairs from different species.

| Species | Name | Gene ID | mRNA ID | Protein ID | Amino acid sequence of mature protein (without signal peptide) | ipTM | pTM |
|---|---|---|---|---|---|---|---|
| *Homo sapiens* | CXCL17 | 284340 | NM_198477 | NP_940879 | SSLNPGVARGHRDRGQASRRWLQEGGQECECKDWFLRAPRRKFMTVSGLPKKQCPCDHFKGNVKKTRHQRHHRKPNKHSRACQQFLKQCQLRSFALPL | 0.67 | 0.72 |
| | GPR25 | 2848 | NM_005298 | NP_005289 | MAPTEPWSPSPGSAPWDYSGLDGLEELELCPAGDLPYGYVYIPALYLAAFAVGLLGNAFVVWLLAGRRGPRRLVDTFVLHLAAADLGFVLTLPLWAAAAALGGRWPFGDGLCKLSSFALAGTRCAGALLLAGMSVDRYLAVVKLLEARPLRTPRCALASCCGVWAVALLAGLPSLVYRGLQPLPGGQDSQCGEEPSHAFQGLSLLLLLLTFVLPLVVTLFCYCRISRRLRRPPHVGRARRNSLRIIFAIESTFVGSWLPFSALRAVFHLARLGALPLPCPLLLALRWGLTIATCLAFVNSCANPLIYLLLDRSFRARALDGACGRTGRLARRISSASSLSRDDSSVFRCRAQAANTASASW | | |
| *Mus musculus* | CXCL17 | 232983 | NM_153576 | NP_705804 | SPNPGVARSHGDQHLAPRRWLLEGGQECECKDWFLQAPKRKATAVLGPPRKQCPCDHVKGREKKNRHQKHHRKSQRPSRACQQFLKRCHLASFALPL | 0.69 | 0.74 |
| | GPR25 | 383563 | NM_001101516 | NP_001094986 | MQSTEPWSPSWGTLSWDYSGSGSLDQVELCPAWNLPYGHAIIPALYLAAFAVGLPGNAFVVWLLSRQRGPRRLVDTFVLHLAAADLGFVLTLPLWAAAEARGGLWPFGDGLCKVSSFALAVTRCAGALLLAGMSVDRYLAVGRPLSARPLRSARCVRAVCGAAWAAAFLAGLPALLYRGLQPSLDGVGSQCAEEPWEALQGVGLLLLLLTFALPLAVTLICYWRVSRRLPRVGRARSNSLRIIFTVESVFVGCWLPFGVLRSLFHLARLQALPLPCSLLLALRWGLTVTTCLAFVNSSANPVIYLLLDRSFRARARFGLCARAGRQVRRISSASSLSRDDSSVFRGRSPKVNSASATW | | |
| *Rattus norvegicus* | CXCL17 | 308436 | NM_001107491 | NP_001100961 | SSPNQEVARHHGDQHQAPRRWLWEGGQECDCKDWSLRVSKRKTTAVLEPPRKQCPCDHVKGSEKKNRRQKHHRKSQRPSRTCQQFLKRCQLASFTLPL | 0.64 | 0.73 |
| | GPR25 | 363993 | NM_001398594 | NP_001385523 | MQSTESWNSSWGTTMSWDYSGSGSLDQVELCPAWELPYSHAIISALYLAAFVVGLTGNTFVVWLLSRRRRPRRLVDTFVLHLAAADLGFVLTLPLWAVAEARGGLWPFGDGLCKISSFALAATRCAGALLLAGMSVDRYLAVGRPLNARPLRSARCVRASCSAVWAAAFLAGLPTLLYRRLQPSLDGEGSQCAEEPSDALQGVGLLLLLLTFALPLAVTLTCYWRVSRRLLRVRRARGNSLRIIFTVESVFVGSWLPFSILRALFYLARLRALPLPCSLLLALRWGLTVATCLAFVHSCANPVIYLLLDRSFRARVRFGPCTRVGRQVRRISSASSLSRDDSSVFWGRSSKANSALAAW | | |
| *Cavia porcellus* | CXCL17 | 100734695 | XM_013146847 | XP_013002301 | SSPKSGVARAHGEQRQASRRWLQEGSRECECKDWFQRALRRKPMTVPALPKKQCPCDHLKVNMKKSRHHKHQRKPNKHSRACQEFLKQCQLASIALPL | 0.69 | 0.75 |
| | GPR25 | 106028732 | XM_063239898 | XP_063095968 | MVATEPWSLLPGTEFWDYSGSGVLGDELESCPSWELPYSYAYVPALYLAAFSVGLLGNTFVVWLLAGRRGPRRLVDTFVLHLAAADLGFVLTLPLWAAAAARGGRWPFGAGLCKLSSFALAGTRCAGALLLAGLSVDRYLAVVKLLEARPLRTPRCALATCCGVWAVALLAGLPSLVFRGLQPFPEGPGSQCGEEPSDAFQAISLLLLLLTFVLPLAVTVFCYWRISRRLRKPPHVGRARRNSLRIIFAVVGTFVGSWLPLSALRAVFHLARLGALPLPCPLVLALRWGLTVATCLAFVHSCANPLIYLLLDRSFRARALQRVCAHAGRQVPRISSASSLSRDDDSVFWGRRQSPNTASASW | | |
| *Felis catus* | CXCL17 | 101086020 | XM_003997765 | XP_003997814 | SSRNPGVARGHRDQRQAPRRWLQEGSQECECKDWFLRAPKRKLMTVPGLPKKQCPCDHFKGSVKKTRHQRHHRKPNKHSRACQQFLTRCQLESFALPL | 0.68 | 0.74 |
| | GPR25 | 101098111 | XM_023247599 | XP_023103367 | MHPTEPRSPSPETASWDYLGSGALEELEPCLAPDLPYGYAYIPALYLAAFVVGLLGNAFVVWLLAARPGPRRLVDTFVLHLAAADLGFVLTLPLWAAAAASGGRWPFGEGLCKLSGFALAGTRCAGALLLAGLSVDRYLAVVKLLDARPLRTPRCALAVCCGVWAVALLAGLPSLAYRGLQPLPGGGGSQCGEEPSDAFQGLSLLVLLLTCVLPLGVTLVCYCRISCRLRRPPHLGRARRNSLRIIFAVESAFVGSWLPFGALRAVFHLARLGAVPLPCGLLLALRWGLSIATCLAFVNSCANPLIYLLLDRSFRARAWRGVCLRADRPARRGSSVSSLCRDDSSVFRSPAGSWERARPANAGSAVP | | |
| *Lontra canadensis* | CXCL17 | 116857606 | XM_032841857 | XP_032697748 | SSPNPGVARGHREVRQAPRTGLHRGSQECDCKDWFLRAPKRKLMAMPEMPKKQCPCDQFKASVKKTRNQRHHRKPNKHWRACQEFLKQCQLASFALPL | 0.67 | 0.72 |
| | GPR25 | 116879773 | XM_032878221 | XP_032734112 | MPLTDPWIPTPRTPSWDYSGSGDLEDLEPCRVRDLPYSYAYVPVLYLAAFVVGLLGNSFVVWLLAGRRGPRRLVDTFVLHLAAADLGLVLTLPLWAVATARGGRWPFGEGLCKLSGFALAGTRCAGALLLAGLSVDRYLAVVKLLDARPLRTPRCALAACCGVWATALLAGLPSLAYRELQPLPGGRGSQCGEEPSDAFQALSLLLLLLTCALPLGVSLVCYCLISRRLRRPPHLGRARRNSLRIIFAVEGAFVGSWLPFGALRAVFHLARLGALPLPCGLLLALRWGLSIATCLAFVNSCANPLIYLLLDRSFRARAWRGVCGRPDRPDRPARGGSSASSLSRDDSSVFRSPAGSWERARTASAGPALP | | |
| *Meles meles* | CXCL17 | 123931218 | XM_045988111 | XP_045844067 | SSPNPGVARGHRDDRQAPRMGLHKGSQECDCKDWFLRAPKRKLMAMSEMPKKQCPCDQFKGGVKKTRHQRHHRKPNKHWRACQEFLKQCQLASFALPL | 0.67 | 0.73 |
| | GPR25 | 123928346 | XM_045983463 | XP_045839419 | MPFTEPWISSPGTPSWDYSGSGDLEDLEPCRFRDLPYSYAYVPVLYLAAFVVGLLGNSFVVWLLAGRRGPRRLVDTFVLHLAAADLGLVLTLPLWAVATARGGRWPFGEGLCKLSGFALACTRCAGALLLAGLSVDRYLAVVKLLDARPLRTPRCALTACCGVWATALLAGLPSLAYRELQPLRGGPGSQCGEEPSDAFQALSLLLLLLTCALPLGVSLACYCLISRRLRRPPHLGRARRNSLRIIFAVEGAFVGSWLPFGALRAVFHLARLGALPLPCGLLLALRWGLSIATCLAFVNSCANPLIYLLLDRSFRARAWRGVCGRPDRPARGGSSASSLSRDDSSVFRSPAGSWERARTANAGPALL | | |
| *Loxodonta africana* | CXCL17 | 100668516 | XM_003406663 | XP_003406711 | SSPNPGVARGHRDQRQASRRSLQKDSQECECKDWLLGAPKRKSMTVTVPGLPKKPCPCDHFKGNVKKIRHQRHHKKPNKHSRACQQFLQRCQLASFALPI | 0.6 | 0.7 |
| | GPR25 | 100672521 | XM_064273428 | XP_064129498 | MHPTEPWSPSPWAGSEDYSGSGALEELEELKLCPSWDLPYSYAYIPVLYLVAFAVGLLGNAFVVWLLAGRRGPRRLVDTFVLHLAAADLGFVLTLPLWAVAAALGGRWPFGESLCKLSSFALAGTRCAGALLLAGMSVDRYLAVVKLLDARPLRTPRCALATCCAVWAVALLAGLPALAYRGLQPLPGGQGSQCGEEASDAFQGLSLLLLLLTFMLPLGVTLFCYCRISRRLRRPPHLGRARRNSLRIIFAIESTFVGSWLPFSALRAIFYLARLGALPLPCSLVLALRWGLTIATCLAFVNSCANPVIYLLLDRSFRARARRGACGTIGRLARRVSSASSLSGDDSSLFRSGACSWGSFQPVQSVSGPL | | |
| *Bos taurus* | CXCL17 | 788717 | NM_001083799 | NP_001077268 | SSHTGVARGQRDQRQASGRWLREGGQECECQDWFLRAPRRTLMAAPRLTKPCPCDHFKGRMKKTRHQRHHRKSNKPSRACQQFLTRCLLESFALPL | 0.65 | 0.73 |
| | GPR25 | 107133259 | XM_015475319 | XP_015330805 | MRPTEPWSPSAGTAPWDYYSGSGAPDELEPEELCAARDLPYSHAYIPALYLAAFAVGLPGNAFVLWLLSGARGRRRLVDTFVQHLAAADLGFVLTLPLWAAAAARGGRWPFGEGLCKLSSFALAGTRCAGALLLAGLSVDRYLAVGRPLAARSPRSRRCALAACAGVWAAALAAGLPSLAFRRLRPLPGQDRGSQCGEESSDAFQGLSLLLLLLTLVLPLAVTVVCYCRVSRRLRGPPHLGRARSRSLRIILAVEGAFVGSWLPFCALRAVFHLASLGALPLPCRLLLALRWGLTVATCLAFVNSCANPLIYLLLDRSFRAQLRQRGACGRADRPARGSSSASSLSGEDGSPFRSPARAGGRAQTASAPSAVGP | | |
| *Physeter catodon* | CXCL17 | 102983204 | XM_024132473 | XP_023988241 | SSNPGIARGHMDQRQASGRWLQDGGQECECKDWFLRAPKRKLMTVPGLPKKPCPCDHFKGRVKKTRHQRHHKKPTKPSRACQQFLRRCQLASFALPL | 0.63 | 0.71 |
| | GPR25 | 112066689 | XM_024130611 | XP_023986379 | MRPTEPWSPSAGTAPWDYSGSGAPEELELEPGPCAARDLPYGYAYIPALYLAAFAVGLPGNAFVLWLLVGRSGPRRLVDTFVQHLAAADLGFVLSLPLWAAAAARGGLWPFGEGLCKLSSFAMAGTRCAGALLLAGLSVDRYLAVGRPLAARARRTRRCALAACGAVWATALAAGLPSLAFRRLRPLPGGRGSQCAEEPSDAFQGLGLLSLLLTLALPLAVTVVCYCLVSRRLRGPALLGRARGRSLRIIVAVEGAFVGSWLPFCALRAVFHLASLRALPLPCGLLLALRWGLSVATCLAFVSSCANPLIYLLLDRSFRAQLRRRGACWRAARAARGGSSASSLSQDYGSPFRSPGRGAGRAQTTNASSAVRP | | |
| *Equus caballus* | CXCL17 | 100629292 | XM_003362327 | XP_003362375 | SSPNPGVAKGHRDRHQASGRWLQAGGQECECKDWLLRAPKRKLMTVPGLPQKQCPCDHFKGNVKKTRHRRHHRKPNKRSRACQQFLKRCQLASLALPL | 0.64 | 0.71 |
| | GPR25 | 111771301 | XM_023632516 | XP_023488284 | MSPTAPWSPSPGATSWDYSGWGALEELELCPSGDLPYAYAYVPVLYLAAFAVGLPGNAFVLWLLAGRRGPRRLVDTFVRHLAAADLGLVLTLPLWAVAAARGGRWPFGEGLCKLSSFALAGTRCAGALLLAALSVARALAVGARARRCPRRVGAVCCGAWVLALLAGLPSLASRGLQPLPGGHGSQCGDESSDAVQGLGLLVLLLTCALPLAVSVLCYCRVSCHLRRPPALGRARARSLRIIFAVEGAFVGSWLPFGALRAVFHLARLGALPLRCRLLLALRWGLTLATCLAFVNSCANPLIYLLLDRTFRARARLWACGRADRLARTVSSASSLSRDDGPAFRGLSSGAGRARTASRAAASL | | |
| *Phascolarctos cinereus* | CXCL17 | 110219727 | XM_021003326 | XP_020858985 | SSNPEGAEGQRDHKLSAKRHSRGSRQACQCEDVFQNIHGGKTVLEAKPPARQCPCDRLKPKQKNAGLGHRKHWRRSCLRFLKQCQLRDITLPL | 0.57 | 0.72 |

| | | | | | | | |
|---|---|---|---|---|---|---|---|
| | GPR25 | 110219523 | XM_021002910 | XP_020858569 | MTTESWSPSPEEGDENYEWSGDYGEPVFCPTPALPFGHIFIPLLYLAAFGAGLMGNSFVVWLLAGQQGPQRLVDTFVLHLAVADLMFVVTLPFWATAVGLRGQWIFGEDLCKVSSFIIAVTRCASTLLLTGMSVDRYLAVVKLLDARPLRNRGCMLMTCAVIWTISILAGVPSLVYRKLLPMPVGPGSLCGDDPTDIFQWLSLLVLFLTFVLPLGIIFFCYSLISQRLWNPPQVGRGRKNNSLRIIFAIVGTFTCSWLPYSILKTFFHIAHLRALPLDCPLLLALRWGLTLSTCLAFVNSSANPVIYLLLDRSFRARAQGTCRRTRPLLASRVSSASSLSGGDSSVFQGWALGRGRNMRLAPRLSILQNS | | |
| *Trichosurus vulpecula* | CXCL17 | 118835868 | XM_036743155 | XP_036599050 | PNPEAAEGQRDHKLPAERHPRGRRQECRCEDIFQNTHGGKRVRVAKPPARQCPCARLKYKRKKPGLGHQKHWRQSCLRFLKQCQLQKINVPL | 0.59 | 0.73 |
| | GPR25 | 118845788 | XM_036753817 | XP_036609712 | MTTESWSPTPGEGDENYEWSGDYGELMFCPTPDLPYGHIFIPLLYLAAFGVGLVGNSFVVWLLAGQQGPRRLVDTFVLHLAMADLMFVVTLPFWATAAGLGGHWTFGEGLCKVSSFIIAVTRCASILLLTGMSVDRYLAVVKLLDARPLRNRGCMLTTCGVIWTISILAGVPSLVYRKLLPMPVGPGSLCGDEPTDIFQWLSLLVLFLTFVLPLGIIIFCYSLISQRLWNPPHLGRGRKNSLRIIFAIVGTFTCSWLPFSILKTFFHIAHLRTLPLDCPLLLVLRWGLTLSTCLAFVNSSANPIIYLLLDRSFRARAHGACKKNHPPLPSRVSSTSSLSGGDSSIFQGWALGWPRNTRLALRLSLLRNP | | |
| *Vombatus ursinus* | CXCL17 | 114040160 | XM_027858170 | XP_027713971 | SSPNPEDAEGQRNHKLPAKRHARGGRQVCRCQDVFQNIHGGKRVQRAKPPARQCPCDRLKYKQKKPGLGHRTHWRRSCLGFLKQCQLRDITLPL | 0.68 | 0.74 |
| | GPR25 | 114029117 | XM_027843464 | XP_027699265 | MTTESWSPSPDEGDENYEWSGDYGEPALCPTPALPFGHVFIPLLYLAAFGAGLIGNSFVVWLLAGQQGPRRLVDTFVLHLAVADLMFVVTLPFWATAVGLHGQWIFGEELCKVSSFIIAVTRCASTLLLAGMSVDRYLAVVKLLDARPLRNRGCMLTTCAVIWIISILAGVPSLVYRKLLPMPVGPGSLCGDDPADIFQGLSLLVLSLTFVLPLGIIFFCYSLISQRLWNPLQLGRGRKNSLRIIFAIVGTFTCSWLPYSILKTFFHIAHLRALPLDCPLLLALRWGLTLSTCLAFVNSSANPIIYLLLDRSFRARARGSCRRTRPPLASRVSSASSLSGGDNSIFQGWALGRGRNTRLAPRLSLLQNP | | |
| *Ornithorhynchus anatinus* | CXCL17 | 103166888 | XM_029066640 | XP_028922473 | SVPHNPGGSRSHGERRQEARRRLGQITHKCRCPGLSQELQRESRARRLQSPRGECPCDNLKVNLKKRPVWHHKGKEGRVHYRHKEACRLFWKQCELEGLSLPI | 0.56 | 0.62 |
| | GPR25 | 100093467 | XM_029069673 | XP_028925506 | MNFAAKNSYDLTEEVPELHPKALFNCPFLLEVASIKCTVSSPLLSLPPARRPSAHPPRGLLHLPLPREPVATAVAAARAVVTAAAAASAGSGASTTSAPGPAAAALSMTSQTWGPSSPGGLGPYSGWGDYEEVGGTVMPWCPTPELPYGYAYIPALYLAAFVVGLVGNGFVVALMVGQPGPRRLVDTFVLHLAVADLVFVCTLPLWAAAAGLGNHWPFGEGLCKLSSFAMAVNRCSSALLLAGMSLDRYLVVVKMLEVRPLRTPGCVVSACGVIWAISLLAGIPSLVFRKLQPAARASGSLCGEEASDAFQSLSLALLVFTFALPLAVILFCYCLISRRLRQHHGLGRHRKNSLRIIFAVVGTFVGSWLPLSLLRAFYHLSRLGVLELPCPLLLALRWGLTLSACLAFINSCANPIIYLSLDRSFRVRALRGACGKASRLVRRASSVSSSRDSRDSGSLFQARGRTRAGSAPGGPG | | |
| *Tachyglossus aculeatus* | CXCL17 | 119946135 | XM_038767563 | XP_038623491 | SVPQNPGDRRSHGERRLAARRRLGELTHKCRCPGSSQELQRESRARRLQSPRGECPCDDLKVNLKKRPMWHHKGKEGRVHYRRKEACRLFWKQCELEGLSLPI | 0.66 | 0.62 |
| | GPR25 | 119930418 | XM_038748745 | XP_038604673 | MTTKISVKSDAVSEPWNSEAGAVVKGDSGSKIKKCTENGSDWQARGSDASKEQDSPIPHRTPLMIAVAAVRAAAVVAVTAASKGGGASATSAPGPAAAALSMTSQTWGPSSPGGLGPYSGWGDYEDVGSTVTPWCPTPELPYGYAYIPALYLAAFVVGLVGNGFVVALMVGQPGPRRLVDTFVLHLAVADLVFVCTLPLWAAAAGLGNHWPFGEGLCKLSSFAMAVNRCSSALLLAGMSLDRYLVVVKMLEVRPLRTPGCVVSACGVIWAISLLAGIPSLVFRKLQPAPRASGSLCGEEVSDAFQSLSLALLVFTFALPLAVILFCYCLISRRLRQHHGLGRQRKNSLRIIFAVVGTFVGSWLPLSLFRVFYHLSRLRVLELPCSLLLALRWGLTLSACLAFINSCANPIIYLSLDRSFRVRALRGACGKASRLVRRASSVSSSRDSQDSGSLFQARGRTRAGSGPCGPGWGPEDP | | |

**Table S5.** Primers and vectors used for generation of human GPR25 expression constructs.

| Expression constructs | Vector for cloning | Enzymes for cleaving cloning vector | Oligo primers for PCR amplification (5' to 3') | Template for PCR amplification | Enzymes for cleaving PCR product | Approach for construct generation |
|---|---|---|---|---|---|---|
| pcDNA6/<br>GPR25 | pcDNA6 | NheI<br>AgeI | Forward: G GCT AGC ATG GCC CCC ACA GAG CCC TGG AGC CCC AGC<br>Reverse: G AC CGG TGG CCA GGA GGC CGA GGC AGT GTT CGC GGC CTG | Genomic DNA from HEK293T cells | NheI<br>AgeI | Ligation by T4 DNA ligase |
| pTRE3G-BI/<br>GPR25-LgBiT:<br>SmBiT-ARRB2 | pTRE3G-BI/GPR83-LgBiT:SmBiT-ARRB2 | EcoRI<br>AgeI<br>(removing GP83) | Forward: AAAG GAA TTC ATG GCC CCC ACA GAG CCC TGG<br>Reverse: GG ACC GGT CCA GGA GGC CGA GGC AGT GTT | pcDNA6/GPR25 | EcoRI<br>AgeI | Ligation by T4 DNA ligase |
| pTRE3G-BI/<br>[W95A]GPR25-LgBiT:<br>SmBiT-ARRB2 | pTRE3G-BI/GPR83-LgBiT:SmBiT-ARRB2 | EcoRI<br>AgeI<br>(removing GP83) | Forward 1: GAA CCG TCA GAT CGC CTG GAG AAT T<br>Reverse 1: TAG CGC CGC CGC CGC GGC CGC CAG CGG CAG CGT GAG CAC<br><br>Forward 2: GTG CTC ACG CTG CCG CTG GCG GCC GCG GCG GCG GCG CTA<br>Reverse 2: GCC ACT GCT AGA CCC TCC GCC | pTRE3G-BI/GPR25-LgBiT:SmBiT-ARRB2 | No cleavage | Gibso assembly of three fragments |
| pTRE3G-BI/<br>[R178A]GPR25-LgBiT:<br>SmBiT-ARRB2 | pTRE3G-BI/GPR83-LgBiT:SmBiT-ARRB2 | EcoRI<br>AgeI<br>(removing GP83) | Forward 1: GAA CCG TCA GAT CGC CTG GAG AAT T<br>Reverse 1: AGG CAG GGG CTG CAA CCC CGC GTA GAC CAG GGA GGG CAG<br><br>Forward 2: CTG CCC TCC CTG GTA TAC GCG GGG TTG CAG CCC CTG CCT<br>Reverse 2: GCC ACT GCT AGA CCC TCC GCC | pTRE3G-BI/GPR25-LgBiT:SmBiT-ARRB2 | No cleavage | Gibso assembly of three fragments |
| pTRE3G-BI/<br>GPR25-LgBiT:<br>SmBiT-ARRB1 | pTRE3G-BI/GPR25-LgBiT:SmBiT-ARRB2 | NheI<br>EcoRV<br>(removing SmBiT-ARRB2) | Forward 1: TCT AGT GGT GGA GGG GGC AGC GGC GGA GGT GGC GAC AAA GGG ACC CGA GTG<br>Forward 2: TAC CGG CTG TTC GAG GAG ATT CTG GGC GGC TCT AGT GGT GGA GGG GGC AGC<br>Forward 3: GCT AGC ATG GGA GTG ACC GGC TAC CGG CTG TTC GAG GAG ATT<br>Forward 4: GTA AAG TCG ACA CCG GGG CCC GCT AGC ATG GGA GTG ACC GGC TAC CGG<br><br>Reverse: GGC TGA TTA TGA TCC TCT GGA GAT ATC GCC GGC GGC CGC CTA TCT GTT GTT GAG CTG TGG AGA | pENTER/ARRB1 | No cleavage | Gibso assembly of two fragments |

pTRE3G-BI/GPR83-LgBiT:SmBiT-ARRB2 was generated in our previous study, reference 31.

The construct pENTER/ARRB1 was purchased from WZ Bio (Jinan, Shandong, China), but it contained a frame-shift mutation in the coding region of human ARRB1 as identified by DNA sequencing. This mutation was corrected via QuikChange mutagenesis in our laboratory.

For oligo primers, the restriction enzyme cleavage site or the fragment pairing with vector is highlighted in yellow, the fraction pairing with PCR template is underlined, the mutation site is shown in red.

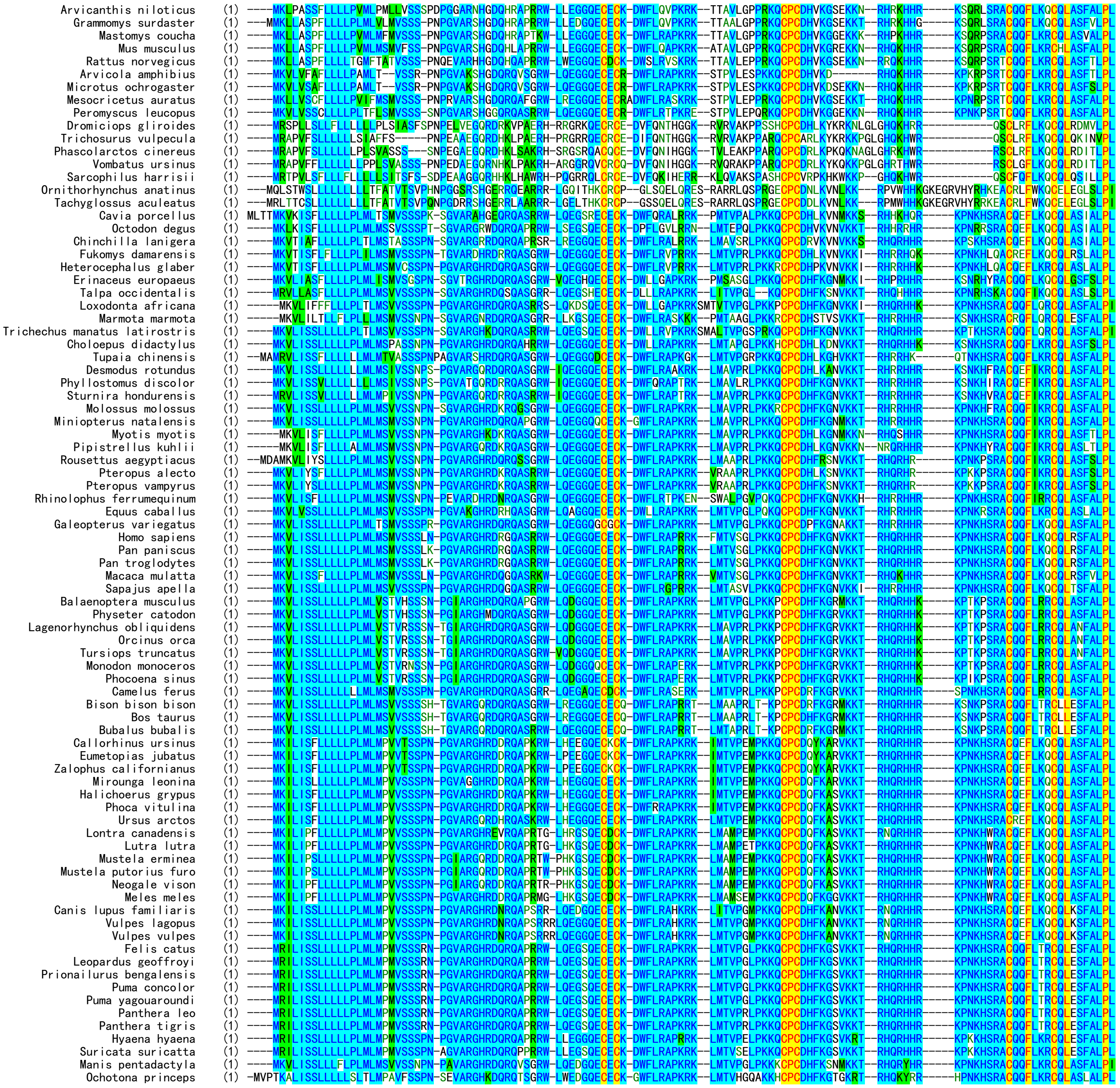

**Fig. S1.** Amino acid sequence alignment of CXCL17 orthologs. Accession numbers of these orthologs are listed in Table S1. They were manually downloaded from NCBI (https://ncbi.nlm.nih.gov/gene) and aligned via AlignX algorithm using the Vector NTI 11.5.1 software.

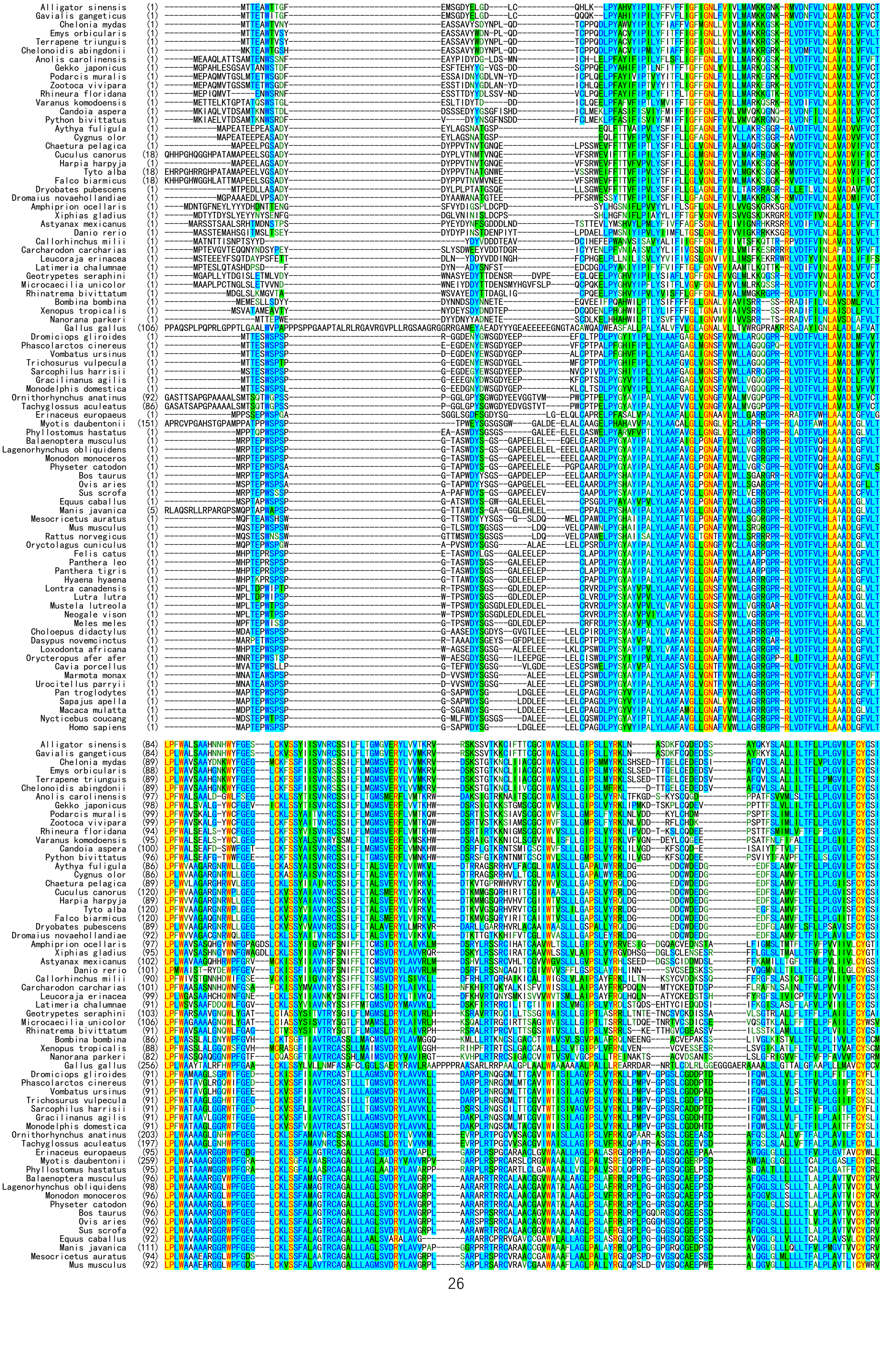

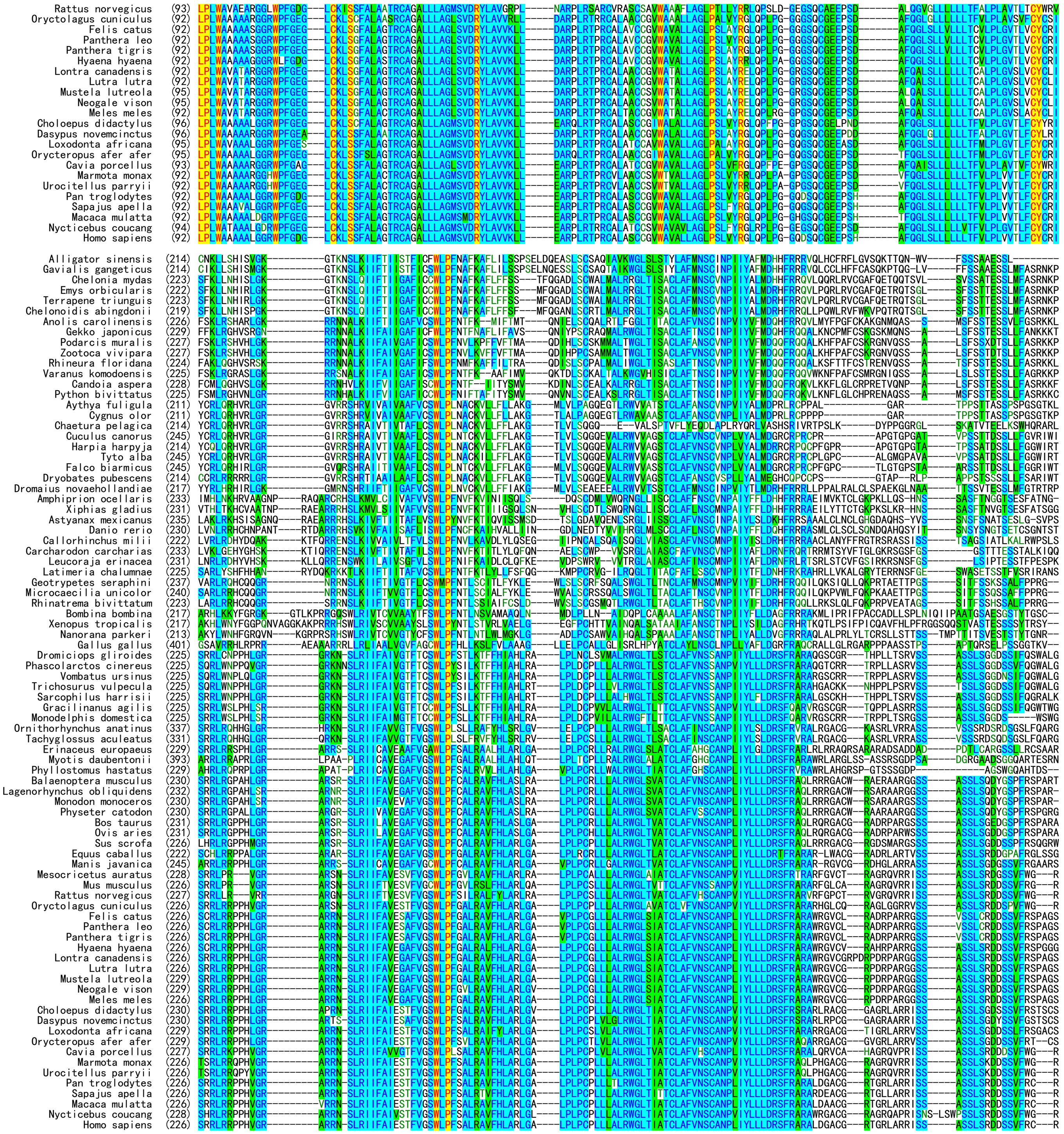

**Fig. S2.** Amino acid sequence alignment of GPR25 orthologs. Accession numbers of these orthologs are listed in Table S2. They were manually downloaded from NCBI (https://ncbi.nlm.nih.gov/gene) and aligned via AlignX algorithm using the Vector NTI 11.5.1 software.

```
                                NdeI
1    ATG CAT CAC CAT CAC CAC CAT ATG GAT GAC GAT GAC AAA AGC AGT CTG AAT CCA GGT GTC GCA CGT GGC CAC CGC
     TAC GTA GTG GTA GTG GTG GTA TAC CTA CTG CTA CTG TTT TCG TCA GAC TTA GGT CCA CAG CGT GCA CCG GTG GCG
      M   H   H   H   H   H   H   M   D   D   D   D   K   S   S   L   N   P   G   V   A   R   G   H   R

76   GAC CGT GGC CAG GCT TCT CGC CGT TGG CTC CAG GAA GGT GGC CAA GAA TGT GAG TGC AAA GAT TGG TTC CTG CGT
     CTG GCA CCG GTC CGA AGA GCG GCA ACC GAG GTC CTT CCA CCG GTT CTT ACA CTC ACG TTT CTA ACC AAG GAC GCA
      D   R   G   Q   A   S   R   R   W   L   Q   E   G   G   Q   E   C   E   C   K   D   W   F   L   R

151  GCC CCG CGC CGT AAA TTC ATG ACC GTG TCC GGT CTG CCG AAG AAA CAG TGC CCT TGT GAT CAT TTC AAG GGC AAT
     CGG GGC GCG GCA TTT AAG TAC TGG CAC AGG CCA GAC GGC TTC TTT GTC ACG GGA ACA CTA GTA AAG TTC CCG TTA
      A   P   R   R   K   F   M   T   V   S   G   L   P   K   K   Q   C   P   C   D   H   F   K   G   N

226  GTT AAG AAA ACG CGC CAC CAA CGT CAT CAC CGT AAG CCG AAC AAA CAT TCA CGC GCG TGC CAG CAA TTT CTC AAA
     CAA TTC TTT TGC GCG GTG GTT GCA GTA GTG GCA TTC GGC TTG TTT GTA AGT GCG CGC ACG GTC GTT AAA GAG TTT
      V   K   K   T   R   H   Q   R   H   H   R   K   P   N   K   H   S   R   A   C   Q   Q   F   L   K

                                                    EcoRI
301  CAA TGT CAG CTT CGT AGC TTT GCT CTG CCT TTG TAA GAA TTC
     GTT ACA GTC GAA GCA TCG AAA CGA GAC GGA AAC ATT CTT AAG
      Q   C   Q   L   R   S   F   A   L   P   L   *
```

**Fig. S3.** The nucleotide and amino acid sequences of the designed CXCL17 precursor. The amino acid sequence of the mature human CXCL17 is shown in red, the enterokinase cleavage site for removal of the N-terminal tag is shaded. The restriction enzyme cleavage sites for molecular cloning are shaded and indicated. The three C-terminal residues removed in [desC3]CXCL17 mutant were highlighted in yellow.

### Untagged human GPR25 in pcDNA6 vector

```
        NheI
1       GCT AGC ATG GCC CCC ACA GAG CCC TGG AGC CCC AGC CCG GGG TCA GCG CCC TGG GAC TAC TCG GGG TTG GAC GGC
        CGA TCG TAC CGG GGG TGT CTC GGG ACC TCG GGG TCG GGC CCC AGT CGC GGG ACC CTG ATG AGC CCC AAC CTG CCG
                 M   A   P   T   E   P   W   S   P   S   P   G   S   A   P   W   D   Y   S   G   L   D   G

76      CTG GAG GAG CTG GAG CTG TGT CCG GCC GGG GAC CTG CCC TAC GGC TAC GTC TAC ATC CCC GCG CTC TAC CTG GCG
        GAC CTC CTC GAC CTC GAC ACA GGC CGG CCC CTG GAC GGG ATG CCG ATG CAG ATG TAG GGG CGC GAG ATG GAC CGC
         L   E   E   L   E   L   C   P   A   G   D   L   P   Y   G   Y   V   Y   I   P   A   L   Y   L   A

151     GCC TTC GCC GTG GGC CTG CTG GGC AAC GCC TTT GTG GTG TGG CTG CTG GCC GGG CGG CGG GGC CCG CGG CGG CTG
        CGG AAG CGG CAC CCG GAC GAC CCG TTG CGG AAA CAC CAC ACC GAC GAC CGG CCC GCC GCC CCG GGC GCC GCC GAC
         A   F   A   V   G   L   L   G   N   A   F   V   V   W   L   L   A   G   R   R   G   P   R   R   L

226     GTG GAT ACC TTC GTG CTG CAC CTG GCG GCA GCT GAC CTG GGC TTC GTG CTC ACG CTG CCG CTG TGG GCC GCG GCG
        CAC CTA TGG AAG CAC GAC GTG GAC CGC CGT CGA CTG GAC CCG AAG CAC GAG TGC GAC GGC GAC ACC CGG CGC CGC
         V   D   T   F   V   L   H   L   A   A   A   D   L   G   F   V   L   T   L   P   L   W   A   A   A

301     GCG GCG CTA GGC GGC CGC TGG CCG TTC GGC GAT GGC CTC TGC AAG CTC AGC AGC TTC GCG CTG GCG GGC ACG CGC
        CGC CGC GAT CCG CCG GCG ACC GGC AAG CCG CTA CCG GAG ACG TTC GAG TCG TCG AAG CGC GAC CGC CCG TGC GCG
         A   A   L   G   G   R   W   P   F   G   D   G   L   C   K   L   S   S   F   A   L   A   G   T   R

376     TGC GCG GGC GCG CTG CTG CTG GCG GGC ATG AGC GTG GAC CGC TAC CTG GCC GTG GTG AAG CTG CTC GAG GCG AGG
        ACG CGC CCG CGC GAC GAC GAC CGC CCG TAC TCG CAC CTG GCG ATG GAC CGG CAC CAC TTC GAC GAG CTC CGC TCC
         C   A   G   A   L   L   L   A   G   M   S   V   D   R   Y   L   A   V   V   K   L   L   E   A   R

451     CCA CTG CGC ACC CCG CGC TGC GCG CTG GCC TCG TGC TGC GGC GTC TGG GCC GTG GCG CTG CTG GCC GGC CTG CCC
        GGT GAC GCG TGG GGC GCG ACG CGC GAC CGG AGC ACG ACG CCG CAG ACC CGG CAC CGC GAC GAC CGG CCG GAC GGG
         P   L   R   T   P   R   C   A   L   A   S   C   C   G   V   W   A   V   A   L   L   A   G   L   P

526     TCC CTG GTC TAC CGG GGG TTG CAG CCC CTG CCT GGG GGC CAG GAC AGC CAG TGC GGC GAG GAG CCC TCC CAC GCC
        AGG GAC CAG ATG GCC CCC AAC GTC GGG GAC GGA CCC CCG GTC CTG TCG GTC ACG CCG CTC CTC GGG AGG GTG CGG
         S   L   V   Y   R   G   L   Q   P   L   P   G   G   Q   D   S   Q   C   G   E   E   P   S   H   A

601     TTC CAG GGC CTC AGC TTG CTG CTG CTG CTG CTG ACC TTC GTG CTG CCC CTG GTC GTC ACC CTC TTC TGC TAC TGC
        AAG GTC CCG GAG TCG AAC GAC GAC GAC GAC GAC TGG AAG CAC GAC GGG GAC CAG CAG TGG GAG AAG ACG ATG ACG
         F   Q   G   L   S   L   L   L   L   L   L   T   F   V   L   P   L   V   V   T   L   F   C   Y   C

676     CGC ATC TCG CGC CGC CTG CGA CGG CCG CCG CAC GTG GGT CGG GCC CGG AGG AAC TCG CTG CGC ATC ATC TTC GCC
        GCG TAG AGC GCG GCG GAC GCT GCC GGC GGC GTG CAC CCA GCC CGG GCC TCC TTG AGC GAC GCG TAG TAG AAG CGG
         R   I   S   R   R   L   R   R   P   P   H   V   G   R   A   R   R   N   S   L   R   I   I   F   A

751     ATC GAG AGC ACG TTT GTG GGC TCC TGG CTG CCC TTC AGC GCC CTG CGG GCC GTC TTC CAC CTG GCG CGT CTG GGG
        TAG CTC TCG TGC AAA CAC CCG AGG ACC GAC GGG AAG TCG CGG GAC GCC CGG CAG AAG GTG GAC CGC GCA GAC CCC
         I   E   S   T   F   V   G   S   W   L   P   F   S   A   L   R   A   V   F   H   L   A   R   L   G

826     GCG CTG CCG CTG CCG TGC CCC CTG CTG CTG GCG CTG CGC TGG GGC CTC ACC ATT GCC ACC TGC CTG GCC TTC GTC
        CGC GAC GGC GAC GGC ACG GGG GAC GAC GAC CGC GAC GCG ACC CCG GAG TGG TAA CGG TGG ACG GAC CGG AAG CAG
         A   L   P   L   P   C   P   L   L   L   A   L   R   W   G   L   T   I   A   T   C   L   A   F   V

901     AAC AGC TGC GCC AAC CCG CTC ATC TAC CTC CTG CTG GAC CGC TCA TTC CGA GCC CGG GCG CTG GAC GGG GCC TGC
        TTG TCG ACG CGG TTG GGC GAG TAG ATG GAG GAC GAC CTG GCG AGT AAG GCT CGG GCC CGC GAC CTG CCC CGG ACG
         N   S   C   A   N   P   L   I   Y   L   L   L   D   R   S   F   R   A   R   A   L   D   G   A   C

976     GGG CGC ACC GGC CGC CTG GCG CGA AGG ATC AGC TCA GCC TCC TCG CTC TCC AGG GAC GAC AGT TCC GTG TTC CGT
        CCC GCG TGG CCG GCG GAC CGC GCT TCC TAG TCG AGT CGG AGG AGC GAG AGG TCC CTG CTG TCA AGG CAC AAG GCA
         G   R   T   G   R   L   A   R   R   I   S   S   A   S   S   L   S   R   D   D   S   S   V   F   R

                                                                    AgeI
1051    TGC CGG GCC CAG GCC GCG AAC ACT GCC TCG GCC TCC TGG CCA CCG GTC TGA TAA
        ACG GCC CGG GTC CGG CGC TTG TGA CGG AGC CGG AGG ACC GGT GGC CAG ACT ATT
         C   R   A   Q   A   A   N   T   A   S   A   S   W   P   P   V   *   *
```

### GPR25-LgBiT in MCS2 of pTRE3G-BI (SmBiT-ARRB1 or SmBiT-ARRB2 in MCS1 of this vector)

```
        EcoRI
1       GAA TTC ATG GCC CCC ACA GAG CCC TGG AGC CCC AGC CCG GGG TCA GCG CCC TGG GAC TAC TCG GGG TTG GAC GGC
        CTT AAG TAC CGG GGG TGT CTC GGG ACC TCG GGG TCG GGC CCC AGT CGC GGG ACC CTG ATG AGC CCC AAC CTG CCG
                 M   A   P   T   E   P   W   S   P   S   P   G   S   A   P   W   D   Y   S   G   L   D   G

76      CTG GAG GAG CTG GAG CTG TGT CCG GCC GGG GAC CTG CCC TAC GGC TAC GTC TAC ATC CCC GCG CTC TAC CTG GCG
        GAC CTC CTC GAC CTC GAC ACA GGC CGG CCC CTG GAC GGG ATG CCG ATG CAG ATG TAG GGG CGC GAG ATG GAC CGC
         L   E   E   L   E   L   C   P   A   G   D   L   P   Y   G   Y   V   Y   I   P   A   L   Y   L   A

151     GCC TTC GCC GTG GGC CTG CTG GGC AAC GCC TTT GTG GTG TGG CTG CTG GCC GGG CGG CGG GGC CCG CGG CGG CTG
        CGG AAG CGG CAC CCG GAC GAC CCG TTG CGG AAA CAC CAC ACC GAC GAC CGG CCC GCC GCC CCG GGC GCC GCC GAC
         A   F   A   V   G   L   L   G   N   A   F   V   V   W   L   L   A   G   R   R   G   P   R   R   L

226     GTG GAT ACC TTC GTG CTG CAC CTG GCG GCA GCT GAC CTG GGC TTC GTG CTC ACG CTG CCG CTG TGG GCC GCG GCG
        CAC CTA TGG AAG CAC GAC GTG GAC CGC CGT CGA CTG GAC CCG AAG CAC GAG TGC GAC GGC GAC ACC CGG CGC CGC
         V   D   T   F   V   L   H   L   A   A   A   D   L   G   F   V   L   T   L   P   L   W   A   A   A

301     GCG GCG CTA GGC GGC CGC TGG CCG TTC GGC GAT GGC CTC TGC AAG CTC AGC AGC TTC GCG CTG GCG GGC ACG CGC
        CGC CGC GAT CCG CCG GCG ACC GGC AAG CCG CTA CCG GAG ACG TTC GAG TCG TCG AAG CGC GAC CGC CCG TGC GCG
         A   A   L   G   G   R   W   P   F   G   D   G   L   C   K   L   S   S   F   A   L   A   G   T   R

376     TGC GCG GGC GCG CTG CTG CTG GCG GGC ATG AGC GTG GAC CGC TAC CTG GCC GTG GTG AAG CTG CTC GAG GCG AGG
```

```
      ACG CGC CCG CGC GAC GAC GAC CGC CCG TAC TCG CAC CTG GCG ATG GAC CGG CAC CAC TTC GAC GAG CTC CGC TCC
       C   A   G   A   L   L   L   A   G   M   S   V   D   R   Y   L   A   V   V   K   L   L   E   A   R

451   CCA CTG CGC ACC CCG CGC TGC GCG CTG GCC TCG TGC TGC GGC GTC TGG GCC GTG GCG CTG CTG GCC GGC CTG CCC
      GGT GAC GCG TGG GGC GCG ACG CGC GAC CGG AGC ACG ACG CCG CAG ACC CGG CAC CGC GAC GAC CGG CCG GAC GGG
       P   L   R   T   P   R   C   A   L   A   S   C   C   G   V   W   A   V   A   L   L   A   G   L   P

526   TCC CTG GTC TAC CGG GGG TTG CAG CCC CTG CCT GGG GGC CAG GAC AGC CAG TGC GGC GAG GAG CCC TCC CAC GCC
      AGG GAC CAG ATG GCC CCC AAC GTC GGG GAC GGA CCC CCG GTC CTG TCG GTC ACG CCG CTC CTC GGG AGG GTG CGG
       S   L   V   Y   R   G   L   Q   P   L   P   G   G   Q   D   S   Q   C   G   E   E   P   S   H   A

601   TTC CAG GGC CTC AGC TTG CTG CTG CTG CTG CTG ACC TTC GTG CTG CCC CTG GTC GTC ACC CTC TTC TGC TAC TGC
      AAG GTC CCG GAG TCG AAC GAC GAC GAC GAC GAC TGG AAG CAC GAC GGG GAC CAG CAG TGG GAG AAG ACG ATG ACG
       F   Q   G   L   S   L   L   L   L   L   L   T   F   V   L   P   L   V   V   T   L   F   C   Y   C

676   CGC ATC TCG CGC CGC CTG CGA CGG CCG CCG CAC GTG GGT CGG GCC CGG AGG AAC TCG CTG CGC ATC ATC TTC GCC
      GCG TAG AGC GCG GCG GAC GCT GCC GGC GGC GTG CAC CCA GCC CGG GCC TCC TTG AGC GAC GCG TAG TAG AAG CGG
       R   I   S   R   R   L   R   R   P   P   H   V   G   R   A   R   R   N   S   L   R   I   I   F   A

751   ATC GAG AGC ACG TTT GTG GGC TCC TGG CTG CCC TTC AGC GCC CTG CGG GCC GTC TTC CAC CTG GCG CGT CTG GGG
      TAG CTC TCG TGC AAA CAC CCG AGG ACC GAC GGG AAG TCG CGG GAC GCC CGG CAG AAG GTG GAC CGC GCA GAC CCC
       I   E   S   T   F   V   G   S   W   L   P   F   S   A   L   R   A   V   F   H   L   A   R   L   G

826   GCG CTG CCG CTG CCG TGC CCC CTG CTG CTG GCG CTG CGC TGG GGC CTC ACC ATT GCC ACC TGC CTG GCC TTC GTC
      CGC GAC GGC GAC GGC ACG GGG GAC GAC GAC CGC GAC GCG ACC CCG GAG TGG TAA CGG TGG ACG GAC CGG AAG CAG
       A   L   P   L   P   C   P   L   L   L   A   L   R   W   G   L   T   I   A   T   C   L   A   F   V

901   AAC AGC TGC GCC AAC CCG CTC ATC TAC CTC CTG CTG GAC CGC TCA TTC CGA GCC CGG GCG CTG GAC GGG GCC TGC
      TTG TCG ACG CGG TTG GGC GAG TAG ATG GAG GAC GAC CTG GCG AGT AAG GCT CGG GCC CGC GAC CTG CCC CGG ACG
       N   S   C   A   N   P   L   I   Y   L   L   L   D   R   S   F   R   A   R   A   L   D   G   A   C

976   GGG CGC ACC GGC CGC CTG GCG CGA AGG ATC AGC TCA GCC TCC TCG CTC TCC AGG GAC GAC AGT TCC GTG TTC CGT
      CCC GCG TGG CCG GCG GAC CGC GCT TCC TAG TCG AGT CGG AGG AGC GAG AGG TCC CTG CTG TCA AGG CAC AAG GCA
       G   R   T   G   R   L   A   R   R   I   S   S   A   S   S   L   S   R   D   D   S   S   V   F   R

                                                          AgeI
1051  TGC CGG GCC CAG GCC GCG AAC ACT GCC TCG GCC TCC TGG ACC GGT ACC GGC GGA GGG TCT AGC AGT GGC GGT GGG
      ACG GCC CGG GTC CGG CGC TTG TGA CGG AGC CGG AGG ACC TGG CCA TGG CCG CCT CCC AGA TCG TCA CCG CCA CCC
       C   R   A   Q   A   A   N   T   A   S   A   S   W   T   G   T   G   G   G   S   S   S   G   G   G

1126  ATG GTC TTC ACA CTC GAA GAT TTC GTT GGG GAC TGG GAA CAG ACA GCC GCC TAC AAC CTG GAC CAA GTC CTT GAA
      TAC CAG AAG TGT GAG CTT CTA AAG CAA CCC CTG ACC CTT GTC TGT CGG CGG ATG TTG GAC CTG GTT CAG GAA CTT
       M   V   F   T   L   E   D   F   V   G   D   W   E   Q   T   A   A   Y   N   L   D   Q   V   L   E

1201  CAG GGA GGT GTG TCC AGT TTG CTG CAG AAT CTC GCC GTG TCC GTA ACT CCG ATC CAA AGG ATT GTC CGG AGC GGT
      GTC CCT CCA CAC AGG TCA AAC GAC GTC TTA GAG CGG CAC AGG CAT TGA GGC TAG GTT TCC TAA CAG GCC TCG CCA
       Q   G   G   V   S   S   L   L   Q   N   L   A   V   S   V   T   P   I   Q   R   I   V   R   S   G

1276  GAA AAT GCC CTG AAG ATC GAC ATC CAT GTC ATC ATC CCG TAT GAA GGT CTG AGC GCC GAC CAA ATG GCC CAG ATC
      CTT TTA CGG GAC TTC TAG CTG TAG GTA CAG TAG TAG GGC ATA CTT CCA GAC TCG CGG CTG GTT TAC CGG GTC TAG
       E   N   A   L   K   I   D   I   H   V   I   I   P   Y   E   G   L   S   A   D   Q   M   A   Q   I

1351  GAA GAG GTG TTT AAG GTG GTG TAC CCT GTG GAT GAT CAT CAC TTT AAG GTG ATC CTG CCC TAT GGC ACA CTG GTA
      CTT CTC CAC AAA TTC CAC CAC ATG GGA CAC CTA CTA GTA GTG AAA TTC CAC TAG GAC GGG ATA CCG TGT GAC CAT
       E   E   V   F   K   V   V   Y   P   V   D   D   H   H   F   K   V   I   L   P   Y   G   T   L   V

1426  ATC GAC GGG GTT ACG CCG AAC ATG CTG AAC TAT TTC GGA CGG CCG TAT GAA GGC ATC GCC GTG TTC GAC GGC AAA
      TAG CTG CCC CAA TGC GGC TTG TAC GAC TTG ATA AAG CCT GCC GGC ATA CTT CCG TAG CGG CAC AAG CTG CCG TTT
       I   D   G   V   T   P   N   M   L   N   Y   F   G   R   P   Y   E   G   I   A   V   F   D   G   K

1501  AAG ATC ACT GTA ACA GGG ACC CTG TGG AAC GGC AAC AAA ATT ATC GAC GAG CGC CTG ATC ACC CCC GAC GGC TCC
      TTC TAG TGA CAT TGT CCC TGG GAC ACC TTG CCG TTG TTT TAA TAG CTG CTC GCG GAC TAG TGG GGG CTG CCG AGG
       K   I   T   V   T   G   T   L   W   N   G   N   K   I   I   D   E   R   L   I   T   P   D   G   S

                                              NdeI
1576  ATG CTG TTC CGA GTA ACC ATC AAC AGT TAA CAT ATG
      TAC GAC AAG GCT CAT TGG TAG TTG TCA ATT GTA TAC
       M   L   F   R   V   T   I   N   S   *
```

**Fig. S4.** The nucleotide and amino acid sequences of human GPR25 constructs. The amino acid sequence of human GPR25 is shown in red, that of LgBiT in blue. The two mutated residues (W95 and R178) are highlighted in yellow.
